# Insights into the role of C3d dimers in B cell activation and Staphylococcal immune evasion

**DOI:** 10.1101/2020.04.22.054700

**Authors:** A.A. Wahid, R.W. Dunphy, A. Macpherson, B.G. Gibson, L. Kulik, K. Whale, C.R. Back, T.M. Hallam, B. Alkhawaja, R.L. Martin, I.P. Meschede, A.D.G. Lawson, V.M. Holers, A.G. Watts, S.J. Crennell, C.L. Harris, K.J. Marchbank, J.M.H. van den Elsen

**Author notes:** Corresponding author addresses: Ayla A. Wahid and Jean M.H. van den Elsen.

## Abstract

Cleavage of C3 to C3a and C3b plays a central role in the generation of complement-mediated defences. Although the thioester-mediated surface deposition of C3b has been well-studied, fluid-phase dimers of C3 fragments remain largely unexplored. Here we present the first X-ray crystal structures of disulphide-linked human C3d dimers and show they undergo structurally-stabilising N-terminal domain swapping when in complex with the *Staphylococcus aureus* immunomodulator Sbi. Through binding studies and flow cytometric analyses we uncover the physiologically-relevant roles of these dimers in crosslinking complement receptor 2 and modulating B cell activation to potentially promote anergy. This potential induction of cellular tolerance by C3d dimers could contribute to Sbi-mediated *S. aureus* immune evasion as well as limit autoreactive immune responses under physiological conditions. Thus, insights gained from our findings could inform the design of novel therapies for autoimmune disorders and enhance our understanding surrounding the importance of complement in the fluid phase.

## Introduction

Activation of the central complement component C3 to C3a and C3b by classical/lectin (C4bC2a) or alternative (C3bBb) pathway C3 convertases plays an essential role in the generation of complement-mediated defence mechanisms against invading microbial pathogens. While the circulating C3a anaphylatoxin is involved in inducing inflammatory immune responses, C3b facilitates opsonophagocytosis and clearance of immune complexes through thioester-mediated opsonisation of primary amine- or hydroxyl-containing antigenic and self surfaces. Attachment of multiple copies of C3b and its breakdown products to antigenic surfaces in this way can result in C3d-complement receptor 2 (CR2/CD21) and antigen-B cell receptor (BCR) co-ligation which generates co-stimulatory signals for B cell activation in a C3d copy-dependent manner^1,2^ and has been widely explored in vaccine design^3-8^.

Structure determination of native C3, C3b and C3c has provided crucial insights into the mechanistic basis behind the activation of C3 to C3b^9,10^ while complexes of C3b with factor I (FI) and the short consensus repeat (SCR) domains 1-4 of its cofactor factor H (FH_1-4_) have revealed the processes through which C3b is proteolytically cleaved into its successive opsonin fragments iC3b and C3dg^11^. Crystal structures have also shed light upon the molecular basis underlying the thioester-mediated attachment of C3d to antigenic surfaces^12^, provided explanations of how the interactions of C3d with its receptors (CR2^13^ and CR3^14^) facilitate the recognition of opsonised antigens, and the mechanisms by which pathogens such as *Staphylococcus aureus* utilise C3d-binding proteins (e.g. Sbi^15^, Efb-C^16,17^ and Ecb/Ehp^18,19^) to inhibit these interactions and evade the immune system. Furthermore, complexes of C3d with FH SCR domains 19 and 20 have been pivotal in understanding the regulatory measures in place to protect host tissues against the indiscriminate attachment of C3d to self versus non-self surfaces^20,21^.

However, while these seminal structural studies alongside an abundance of functional investigations have advanced our knowledge surrounding the interaction of C3 fragments with self and non-self surfaces at a molecular level, our understanding of the structural and functional aspects of fluid phase C3 activation products remains incomplete. During activation in the fluid phase, most of the C3 molecules do not covalently attach to surface-exposed hydroxyl- or amine-nucleophiles but instead the highly reactive Cys–Gln thioester moiety within the thioester-containing C3d domain (TED) of C3 undergoes hydrolysis resulting in the generation of C3(H_2_O) and formation of the C3(H_2_O)Bb alternative pathway (AP) C3 convertase. Only approximately 10% of C3b (∼1 mg mL^-1^) generated by these fluid phase or surface-associated convertases is deposited onto reactive surfaces^22^, leaving the remaining 90% to react with water wherein exposure of the cysteine free sulfhydryl can lead to dimerisation of C3b and its subsequent breakdown products^23,24^.

Disulphide or thioester-linked C3b dimers^25^ have been found to bind CR1 with 25-fold higher affinity than monomeric C3b^26^, induce histaminase release from human polymorphonuclear leukocytes^27^, serve as binding platforms for factor B fragment Bb during formation of AP C5 convertases^28,29^ and act as potent AP activators in complex with IgG^30^. Dimers of C3dg have also been isolated from C3-activated human serum following omission of N-ethylmaleimide^31^ and the propensity of recombinant C3d to dimerise has been reported previously^32,33^. A crystal structure of dimeric C3dg purified from rat serum^34^ provides further crucial evidence of the endogenous existence of these dimers. However, aside from this severely truncated C3dg dimer which is believed to have undergone proteolytic truncation during the crystallisation process^35^, there is currently a gap in knowledge surrounding the structural significance of disulphide-linked dimers of C3 fragments as the thioester cysteine sulfhydryl is routinely removed prior to structural analyses. For instance, the free cysteine of C3b has been reacted with iodoacetamide prior to structural determination^10,36,37^ and the vast majority of C3d constructs used for structural studies to date have harboured a cysteine to alanine substitution in order to prevent dimerisation^12,13,20,21^.

In this study we therefore aimed to delineate the molecular details and explore the functional significance of dimeric human C3 fragments that can form following activation of C3 in the fluid phase. We present the first crystal structure of a human C3d dimer at 2.0 Å where dimerisation is mediated by disulphide linkage of the thioester cysteine residues. In addition, through X-ray crystallography, we show this C3d dimer undergoes structurally-stabilising N-terminal 3D domain swapping when bound to domain IV of the *S. aureus* C3-activating immune evasion protein Sbi, thereby providing the first example of this ligand-induced structural phenomenon in a complement protein. Through surface plasmon resonance (SPR) binding studies and cell experiments using mouse splenocytes and human PBMC we show how dimeric C3d crosslinks surface-bound CR2 and can modulate B cell activation to promote the establishment of tolerance/anergy by downregulating CD40 in a more robust manner than C3d monomers. Thus, in the future these newly-discovered physiologically-relevant roles of C3 fragment dimers could inform the design of autoimmune therapies and help to further elucidate the significance of complement in the fluid phase as it interacts with cells of the adaptive immune system and beyond.

## Results

In order to elucidate the importance of dimeric human C3 fragments, here we determined the crystal structures of disulphide-linked human C3d dimers in the absence or presence of Sbi domain IV. Following SPR studies comparing the binding kinetics and avidity of dimeric and monomeric C3d, C3d-induced changes in the activation state of B cells were explored using flow cytometric analyses.

### Disulphide linkage of the thioester cysteine results in C3d dimerisation

A crystal structure of wild-type human C3d, harbouring a cysteine at position 17/1010 (C3d numbering/intact pre-pro C3 numbering) (C3d^17C^), was obtained at 2.0 Å resolution (**Figure 1, Supplementary Tables S1** and **S2**). The structure clearly shows the formation of a dimer mediated by partial disulphide linkage of the thioester cysteine residues at position 17/1010 in both monomeric chains. This 17C-17C disulphide creates a link between the two C3d monomers at the C-terminus of helix α1, causing the convex molecular surfaces of the monomers to orient towards each other whilst simultaneously exposing their concave binding faces (**Figure 1a**). Closer examination of the C3d^17C^ dimer interaction surface (**Figure 1b**) confirms that the overall (α-α)_6_ barrel configuration of both monomers remains unchanged and comparable to previously-published structures of monomeric C3d (0.61 Å (chain A)/0.40 Å (chain B) main chain (M1-P294) RMSD relative to PDB: 1C3D). The 2Fo-Fc electron density map at the C3d^17C^ dimer interface shows that chain B residue 17C (**Figure 1b** inset) adopts a dual conformation with one conformer existing in an unpaired oxidized form. This indicates the disulphide bond linking the two C3d monomers can occur in a partially disconnected state which is consistent with results from size exclusion chromatography experiments suggesting C3d^17C^ exists in a monomer-dimer equilibrium in solution (**Supplementary Figure S1**).

**Figure 1.**
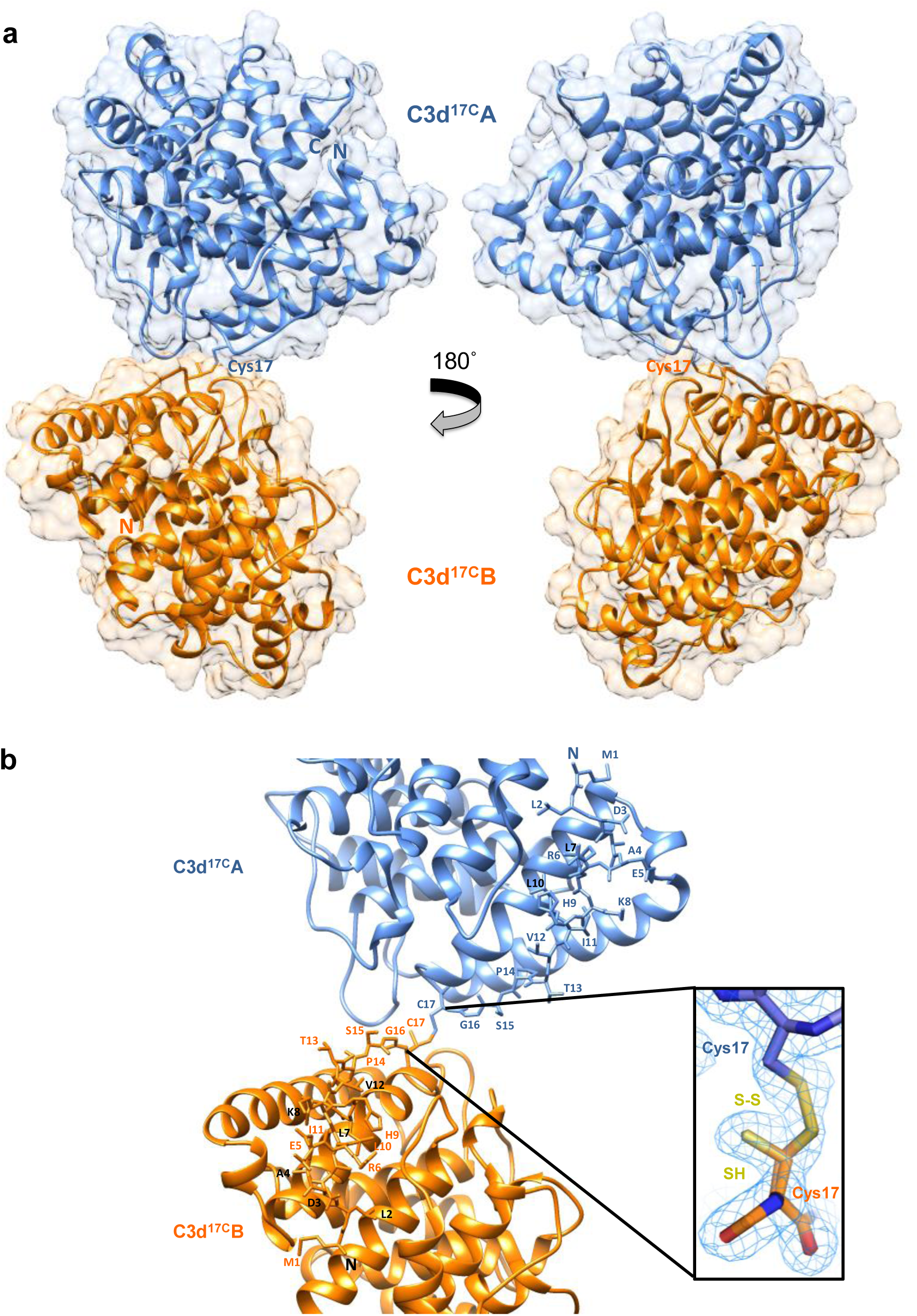
Structure of a disulphide-linked human C3d^17C^ dimer at 2.0 Å resolution. (**a**) The ribbon diagram shows disulphide linkage of the monomeric subunits at position Cys17 results in the formation of a dimer 92.37 Å in length with a 0.61 Å (chain A)/0.40 Å (chain B) main chain (M1-P294) RMSD relative to the structure of C3d^17A^ (PDB:1C3D). (**b**) Enlarged view of the C3d^17C^ dimer interface showing the side chains of helix α1 residues M1-C17. Inset: electron density contoured at 1.0 σ of the partially broken C17-C17 interchain disulphide bond (2.07 Å) resulting from oxidation of one conformer of Chain B Cys17. PDB submission code: 6RMT. See Supplementary Tables S1 and S2 for data collection and refinement statistics.

Superimposition of the ligand-binding domains of CR2 (SCR1-2), the α_M_I integrin domain of CR3 or SCRs 19-20 of FH on to the dimeric C3d^17C^ structure fails to generate any molecular clashes (**Supplementary Figures S2a-d**). This important observation suggests dimerisation does not cause any interference in the formation of complexes between C3d or C3dg and their most physiologically-relevant binding partners. *Staphylococcus aureus* immune evasion proteins such as Efb-C, Ecb/Ehp and Sbi-IV are also predicted to bind the C3d^17C^ dimer without any hindrance as the concave surfaces of both monomers are exposed and accessible. Significantly, as CR2 and CR3 interact with the C3d^17C^ dimer via opposing surfaces, complement receptor crosslinking could play an important role in the function of C3d dimers (**Supplementary Figure S2a-c**). Moreover, the absence of steric hindrance following superimposition of the C3d^17C^ dimer onto the C3b TED domain (**Supplementary Figure S2e**), suggests dimerisation of C3b, as proposed previously^23,24^, could occur in a similar fashion to C3d without affecting the ability of C3b to interact with the complement regulators FH and FI.

### Sbi-IV induces N-terminal helix swapping of dimeric C3d

Following on from our structural analyses of the C3d^17C^ dimer alone, we used co-crystallisation to determine the structure of a complex between C3d^17C^ and Sbi-IV at 2.4 Å (**Figure 2, Supplementary Tables S1** and **S2**) in the interests of gaining an understanding of how dimeric C3d(g) could be exploited by pathogens such as *S. aureus* to modulate immune responses. Previous studies have shown Sbi-IV, which adopts a three-helix bundle fold^38^, binds the acidic residue-lined concave face of C3d^15^ and together with Sbi domain III can be harnessed as a vaccine adjuvant that promotes the opsonisation of antigens with a ‘natural’ coat of C3 breakdown products^8^ via AP activation-mediated consumptive cleavage of C3^39^.

**Figure 2.**
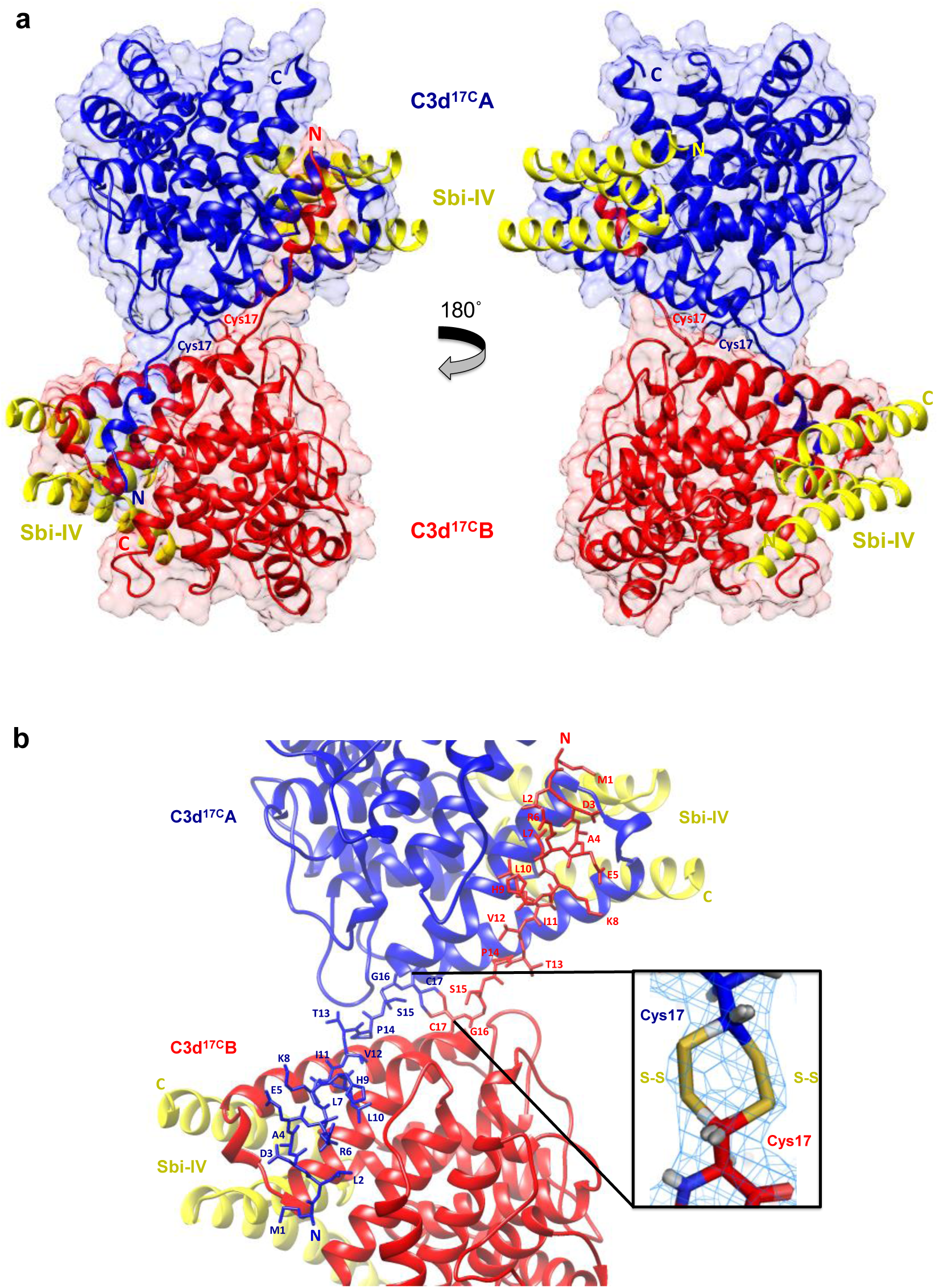
Structure of an N-terminal helix-swapped human C3d^17C^ dimer in complex with Sbi-IV at 2.4 Å resolution. (**a**) In addition to a C17-C17 disulphide linkage, the ribbon diagrams show reciprocal swapping of the N-terminal α1 helix region M1-G16 between the two C3d chains resulting in a domain-swapped dimeric complex 92.31 Å in length with a 0.80 Å (chain A)/0.76 Å (chain B) main chain (G16-P294) RMSD relative to the structure of C3d^17A^ (PDB:1C3D). (**b**) Enlarged view of the C3d^17C^ dimer interface when in complex with Sbi-IV showing side chains of the swapped helix α1 region. Inset: electron density contoured at 1.0 σ of two possible C17-C17 interchain disulphide bonds resulting from dual conformation of Chain A and Chain B residue Cys17 (Chain A C17 conformer A-Chain B C17 conformer A: 2.04 Å, Chain A C17 conformer B-Chain B C17 conformer B: 2.06 Å) with a preference for the conformer B disulphide (Chain B C17 conformer B occupancy: 0.66). PDB submission code: 6RMU. See Supplementary Tables S1 and S2 for data collection and refinement statistics.

The C3d^17C^-Sbi-IV complex presented here shows both the opposing concave binding surfaces of the C3d^17C^ dimer are occupied by domain IV of the Sbi immune evasion protein (**Figure 2a**) in the same manner observed previously by Clark and colleagues^15^. In our structure, occlusion caused by the tight dimer interactions prevents secondary binding of Sbi-IV to the convex surface of C3d close to the thioester region described in the same study (**Supplementary Figure S3**). In clear contrast to the unliganded C3d^17C^ dimer shown in **Figure 1a**, the C3d^17C^ dimer bound to Sbi-IV has undergone a helix swap, a relatively rare structural manifestation whereby a reciprocal exchange between the identical N-terminal α1 helices of the two C3d chains has occurred (**Figure 2a**). As depicted in **Figure 2b**, the helix α1 region M1/994 – G16/1009, located N-terminally from C17/1010, has undergone a full 3D domain swap in both C3d^17C^ molecules, recreating identical interactions with the opposing C3d recipient without causing appreciable differences in relative orientation (0.80 Å (chain A)/0.76 Å (chain B) main chain (G16-P294) RMSDs relative to PDB: 1C3D) (**Figure 3**). The fact that the swap is found to occur at a neutral pH of 8 rather than the acidic pH (3.5-4.0) known to induce domain swapping in other proteins^40^, provides support for its physiological relevance. Unlike the unliganded C3d^17C^ dimer, the 2Fo-Fc electron density map of dimeric C3d^17C^ in the Sbi-IV bound state distinctly shows the formation of two alternative 17C-17C inter-chain disulphide bond configurations at the C3d dimer interface that both remain intact (**Figure 2b** inset).

**Figure 3.**
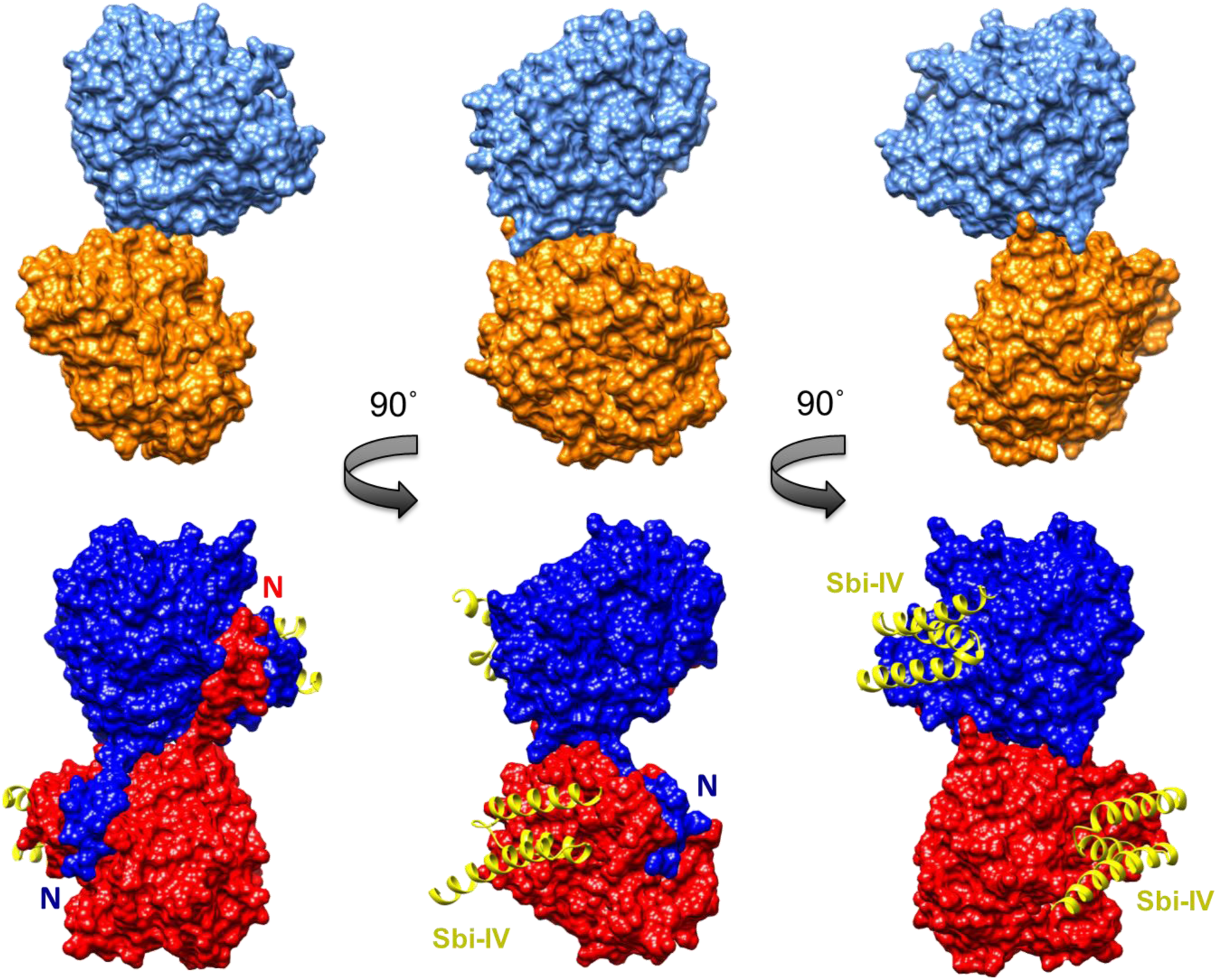
Molecular surface comparison of the C3d^17C^ dimer (top) and the C3d^17C^ dimer in complex with Sbi-IV (bottom). Solid molecular surface representations in three different orientations rotated by 90° angles counter-clockwise are shown where the N-terminal helix swap can be seen in the C3d^17C^ dimer-Sbi-IV complex.

### Helix swapping structurally stabilises dimeric C3d

In addition to stabilising the 17C-17C inter-chain disulphide, the helix α1 swap causes a substantial increase (617%) in joint buried surface area from 720.6 Å^2^ in the C3d^17C^ dimer to 4445.4 Å^2^ in the Sbi-IV-bound helix-swapped C3d^17C^ dimer. This expansion in interface area is supported by molecular interactions including 14 hydrogen bonds and 4 ionic interactions which are absent in the unliganded dimer. Of note, the exact region (M1/994 – G16/1009) located at the extreme N-terminus of C3d^17C^ involved in helix swapping as well as the 17C-17C inter-chain disulphide is absent in the structure of the truncated rat C3d dimer crystallised following proteolytic breakdown of C3dg purified from serum^34^. It is therefore possible the helix swap observed here buries the newly-exposed surface prior to a further degradation step, producing a more compact C3d dimer, C3dt.

Further evidence of an Sbi-IV-induced stabilisation effect was also acquired through near-UV (320-250 nm) circular dichroism (CD) spectroscopic analyses of C3d collected as a function of temperature where the absorption of aromatic side chains and disulphide bonds, absent in Sbi-IV, provides information about the protein’s tertiary structure^41^ (**Supplementary Figure S4**). Here, Sbi-IV was found to significantly increase the melting temperature (ΔTm) of C3d^17C^ by 8.7°C from 50.98 ± 1.41°C in the absence of Sbi-IV (**Supplementary Figure S4a**) to 59.67 ± 1.65°C in the presence of Sbi-IV (**Supplementary Figure S4b**), suggesting that Sbi-IV-mediated helix swapping also enhances the structural stability of dimeric C3d^17C^ in solution. Although Sbi-IV also raises the ΔTm of C3d^17A^ by ∼9.0°C, monomeric C3d^17A^ is less thermally stable (**Supplementary Figure S5a**) and unlike dimeric C3d^17C^ undergoes precipitation at temperatures above 65°C in the presence of wild-type Sbi-IV (**Supplementary Figures S5b**) but not with an N-terminally truncated Sbi-IV mutant (V1_D11del) lacking key residues that were shown to interact with helix α1 on the convex surface of C3d in the crystal structure and NMR analysis of the Sbi-IV:C3d^17A^ complex^15^ (**Supplementary Figure S5c**). This precipitation of C3d^17A^ in the presence of wild-type Sbi-IV, and lack of it in the presence of the Sbi-IV truncation mutant V1_D11del, suggests that the potential displacement of the N-terminal α1 helical region of C3d in the absence of close C3d dimer interactions destabilises the three-dimensional structure of C3d, leading to aggregation and precipitation (see **Supplementary Figures S4c** and **S5d** for sigmoidal fits of thermal denaturation curves used for Tm calculations and **Supplementary Figure S6** for C3d^17C^ and C3d^17A^ far-UV secondary structure analysis).

### C3d dimers can crosslink CR2 and FH_19-20_

As our structural analyses revealed the propensity of C3d^17C^ to dimerise, we next analysed the binding interactions of C3d dimers in comparison to monomeric C3d^17A^ using CR2 and FH_19-20_ as two important known binding partners. Given that both the crystal structure of dimeric C3d^17C^ (**Figure 1b** inset) and size exclusion chromatography experiments (**Supplementary Figure S1**) showed the inter-chain disulphide bond at the dimer interface to be inherently unstable, chemical conjugation of the two C3d^17C^ monomers at position 17C through use of a bromine-based linear linker (N,N’-(propane-1,3-diyl) bis(2-bromoacetamide)) was employed to create a more stable dimer amenable to purification (see Materials and Methods; **Supplementary Figures S7-S9b**). The N,N’-(propane-1,3-diyl) bis(2-bromoacetamide) linker was used as this class of chemical compound has been shown to selectively crosslink cysteine residues located within close spatial proximity^42^. Dimeric C3d^17C^ resulting from this chemical crosslinking reaction was subsequently validated using particle analysis (**Supplementary Figure S9c**), analytical ultracentrifugation (**Supplementary Figure S9d**) and mass spectrometry (**Supplementary Tables S3** and **S4, Supplementary Figure 9e**) and utilised in SPR spectroscopy studies to gain insights into its binding patterns and kinetics.

In contrast to monomeric C3d^17A^ which displays a conventional association-steady state-dissociation binding pattern when flowed over surface-immobilised CR2-Fc and FH_19-20_, the binding of dimeric C3d^17C^ to the same ligands was found to be noticeably distinct and suggestive of a two-state binding interaction (**Figure 4a**). At low concentrations up to the first replicate of 15.63 nM (dashed line), the highly avid interactions with negligible dissociation indicate a bivalent binding mode whereby the C3d^17C^ dimer crosslinks two CR2-Fc or two FH_19-20_ molecules. During the first injection of 15.63 nM C3d^17C^ dimer, 25 RU of material binds to the surface and 10 RU remain avidly bound to the surface after the regeneration. While at the second 15.63 nM injection, 18 RU of material binds to the surface and only 2RU remains avidly bound at the end (**Supplementary Figure S10**). In both cases an equivalent amount of material is eluted during regeneration illustrating that the first injection likely saturates the highly avid binding sites. As the surface cannot be fully regenerated of these high avidity complexes, the subsequent cycle commences at a higher baseline response. At this point and higher concentrations, the high avidity binding sites for dimeric C3d^17C^ on CR2-Fc or FH_19-20_ remain saturated causing the binding mode to switch to less favourable/avid readily-disrupted interactions suggestive of 1:1 binding between the C3d^17C^ dimer and CR2-Fc or FH_19-20_ although some cross-linked C3d^17C^ dimer-CR2-Fc and C3d^17C^ dimer-FH_19-20_ complexes persist (1-2 RU of material remaining bound to the surface after regeneration). At the highest three concentrations of dimeric C3d^17C^ (62.5-250 nM), the less favourable interactions which are readily eluted from the surface dominate (**Figure 4a** inset). Consistent with these results, the unusual binding patterns observed here were also evident in a further two independent experiments (**Supplementary Figure S11**) and cannot be attributed to higher order species of analyte or ligand as the biophysical techniques performed showed the dimeric C3d^17C^ and FH_19-20_ preparations used did not contain aggregates (**Supplementary Figures S9c, S9d** and **S12**). Models illustrating the binding events described here are presented in **Figure 4b** (for CR2-Fc) and **Figure 4c** (for FH_19-20_).

**Figure 4.**
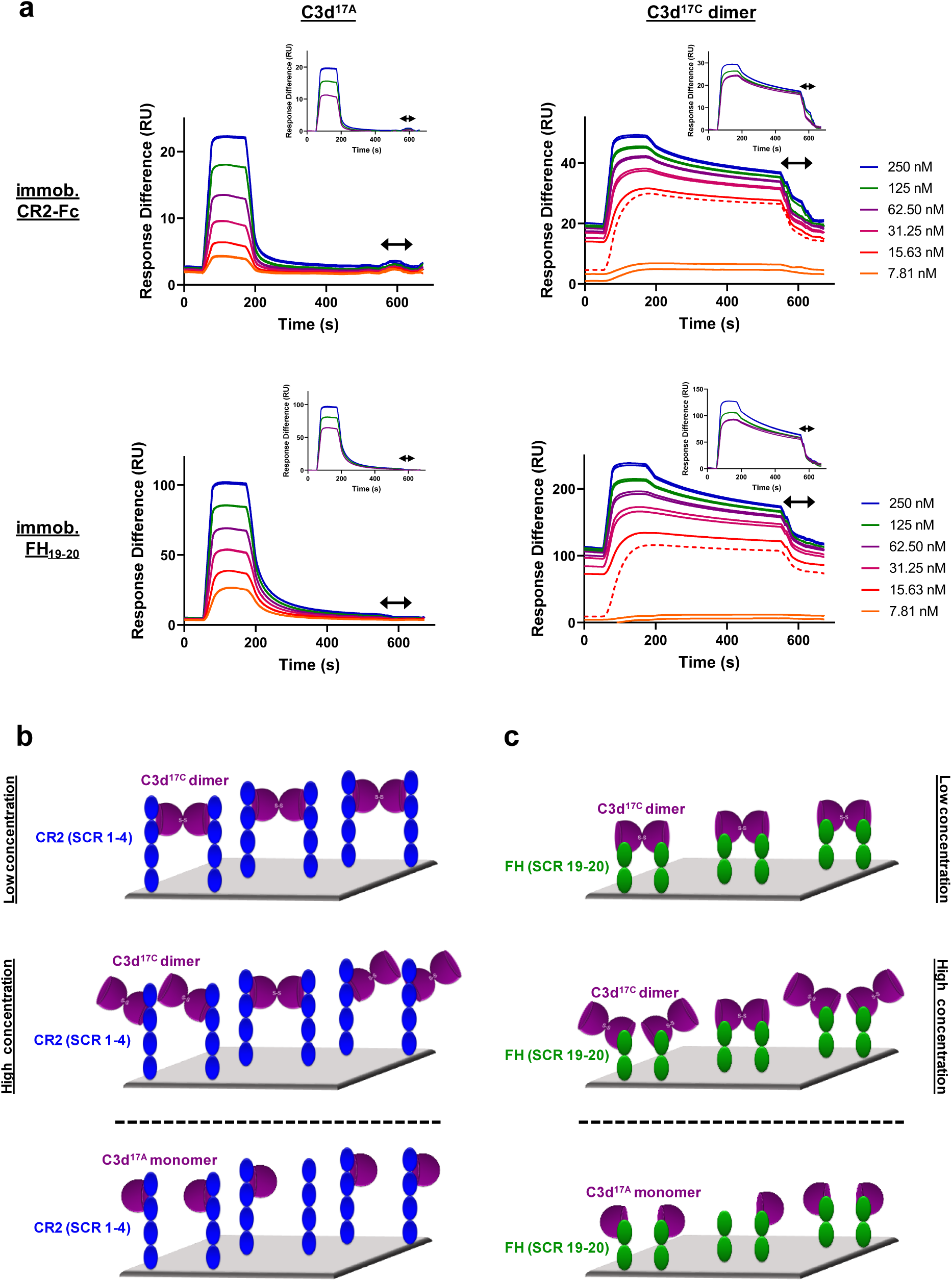
Dimeric C3d^17C^ crosslinks CR2 and FH_19-20_. (**a**) SPR sensorgrams showing serially-diluted concentrations of 250 nM monomeric C3d^17A^ (left) or dimeric C3d^17C^ (right) flowed in duplicate over flow cells of a CM5 senor chip immobilised with CR2-Fc (top) or FH_19-20_ (bottom). The binding of C3d^17A^ to CR2-Fc and FH_19-20_ follows a conventional association-steady state-dissociation pattern while the binding of dimeric C3d^17C^ to the same ligands generates an unusual two-state binding interaction. At concentrations up to the first 15.63 nM replicate (dashed line) the binding patterns depict highly avid interactions suggestive of the formation of dimeric C3d^17C^-CR2-Fc and dimeric C3d^17C^-FH_19-20_ crosslinked complexes which are not fully eluted from the surface. Thus, the subsequent injection cycles commence at a higher baseline response where the high avidity binding sites for dimeric C3d^17C^ on CR2-Fc or FH_19-20_ remain saturated. This causes the binding mode to switch to less favourable, readily-disrupted interactions suggestive of the formation of 1:1 complexes, although some crosslinked complexes persist. Inset: baseline-adjusted sensorgrams showing the less favourable interactions at higher concentrations of dimeric C3d^17C^ which are readily eluted from the surface. Arrows depict the regeneration period. See Supplementary Figure S10 for further details and Supplementary Figure S11 for results from an additional two independent experiments. (**b**,**c**) Schematic models depicting the proposed mechanistic basis behind dimeric C3d^17C^-mediated crosslinking of surface-associated CR2 (SCR 1-4) (**b**) and FH (SCR 19-20) (**c**). At low concentrations, C3d^17C^ dimers crosslink two surface-associated CR2 (SCR 1-4) or FH_19-20_ molecules via highly avid interactions involving the acidic residue-lined concave face of C3d and SCRs 1 and 2 of CR2 or the convex surface and domain 19 of FH (top). Once a critical threshold concentration has been surpassed, the increase in dimeric C3d^17C^ molecules relative to available CR2 or FH_19-20_ binding sites outcompetes the second binding site on C3d^17C^ dimers and favours the formation of 1:1 complexes (middle). Unlike C3d^17C^ dimers, monomeric C3d^17A^ lacks the ability to crosslink CR2 or FH_19-20_ and is restricted to the formation of 1:1 complexes (bottom).

### Dimeric C3d is a more potent modulator of B cell activation than monomeric C3d

Following on from our SPR studies, which indicated dimeric C3d may have the capacity to crosslink CR2, our next aim was to analyse the effects of dimeric compared to monomeric C3d on B cell activation. Flow cytometry was employed to examine changes in the expression of four surface-associated B cell activation markers (CD40, CD69, CD71 and CD86) resulting from stimulation of isolated human B cells with monomeric C3d^17A^ or chemically-linked dimeric C3d^17C^ alone or in the presence of BCR-crosslinking anti-IgM F(ab^’^)_2_. As shown in **Supplementary Figure S13**, although agonism of the BCR by anti-IgM significantly enhances expression of all the activation markers (except CD40), neither monomeric C3d^17A^ nor dimeric C3d^17C^ appears to influence the activation of isolated B cells in an appreciable manner, as measured by the markers examined.

A more general approach, using Ca^2+^ influx as a measure of B cell activation was therefore taken next. Here, incubation of B220^+^ mouse splenocytes with monomeric C3d^17A^ or dimeric C3d^17C^ prior to stimulation with a suboptimal dose of a biotinylated-anti-IgM/C3dg-biotin/streptavidin (a-IgM-b/C3dg-b/ST) BCR/CR2-crosslinking complex was found to inhibit BCR/CR2-mediated Ca^2+^ influx in a concentration-dependent manner (**Supplementary Figure S14**). The observed blocking effect was more pronounced following treatment with dimeric C3d^17C^, particularly at the lower concentration of 4 µg (**Supplementary Figure S14a**), and for both constructs is only evident when C3d is added ahead of the a-IgM-b/C3dg-b/ST complex and when a suboptimal dose of anti-IgM (i.e. unable to trigger Ca^2+^ influx in the absence of CR2 engagement) within the crosslinking complex is used. Thus, the perceived inhibition of Ca^2+^ influx and hence B cell activation is likely to result from sequestration of CR2 by monomeric C3d^17A^, and to a greater extent, due to avidity and possibly via CR2-CR2 crosslinking as suggested by our SPR experiments, dimeric C3d^17C^, reducing the proportion of CR2 available for crosslinking with the BCR.

In order to further investigate C3d-mediated changes in the activation state of B cells within mixed populations of cells, as would occur *in vivo*, flow cytometry was utilised to explore differences in the expression of CD40, CD69, CD71 and CD86 on CD19^+^ cells within donor human PBMC samples (see **Supplementary Figures S15** and **S16** for gating strategy applied). In contrast to the results gathered from isolated human B cells (**Supplementary Figure 13**), a clear dose-dependent relationship between C3d and B cell activation was observed in these experiments indicating other mononuclear cell types may be involved in B cell responsiveness, as measured by expression of the markers analysed, to free C3d (**Figure 5**). At concentrations ≥ 10 nM, both monomeric C3d^17A^ and chemically-linked dimeric C3d^17C^ are able to enhance expression of the early B cell activation markers CD69 and CD86 even in the absence of BCR engagement by anti-IgM. In concert with anti-IgM although both forms of C3d synergistically upregulate expression of these markers in a concentration-dependent manner, dimeric C3d^17C^ is found to be approximately three-fold more effective at enhancing activation than monomeric C3d^17A^ (47nM vs 139 nM (CD69) and 18 nM vs 59 nM (CD86) geometric mean EC50s), perhaps through more avid interactions with CR2.

**Figure 5.**
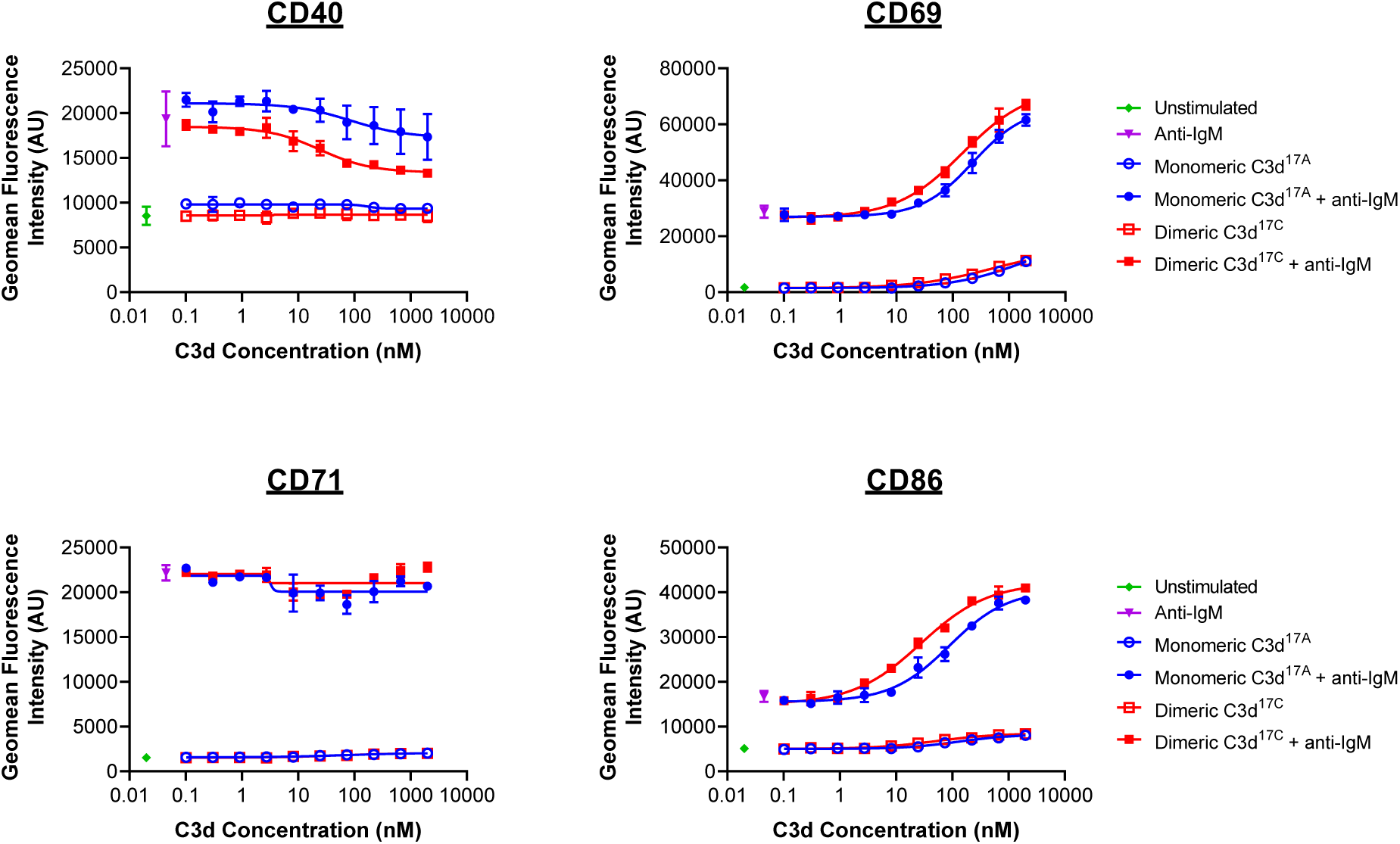
Monomeric C3d^17A^ and dimeric C3d^17C^ induce differential marker-specific changes in the activation state of PBMC B cell populations. Flow cytometric analysis of CD19^+^ B cells stimulated with monomeric C3d^17A^ or dimeric C3d^17C^ in the presence or absence of BCR-crosslinking anti-IgM F(ab^’^)_2_ (10 µg mL^-1^) reveals C3d-induced changes in the expression of surface-associated B cell activation markers. While no C3d-mediated changes in CD71 expression are evident, at higher concentrations (≥ 3 nM) both monomeric C3d^17A^ and dimeric C3d^17C^ appear to downregulate CD40, with a more pronounced reduction in expression in the presence of dimeric C3d^17C^. Conversely, in the presence of anti-IgM, both monomeric C3d^17A^ and to a greater extent dimeric C3d^17C^ synergistically upregulate CD69 and CD86 although at concentrations ≥ 10 nM both forms of C3d are also capable of enhancing expression of these activation markers in the absence of anti-IgM. Data are of PBMC B cell populations from a representative donor and displayed as mean values (n=2) ± standard deviation from the mean with curves fitted using a non-linear regression model. Results from an additional two donors can be found in Supplementary Figure S17.

Interestingly, in contrast to CD69 and CD86, both monomeric C3d^17A^ and dimeric C3d^17C^ appear to downregulate CD40, particularly in the presence of anti-IgM, with a more pronounced reduction in expression evident in the presence of dimeric C3d^17C^. Differently still, despite achieving a substantial increase in expression in the presence of anti-IgM in the experimental time period, CD71 does not appear to be influenced by either form of C3d. Importantly, the differential marker-specific trends observed are consistent across cells gathered from all three donors analysed (data from donors 2 and 3 can be found in **Supplementary Figure S17**) suggesting that free C3d (unattached to an antigen) may regulate B cell activation in a selective manner and that dimeric C3d may have more potent modulatory roles than C3d monomers.

## Discussion

In the past, pre-treatment of C3b with sulfhydryl-alkylating agents and routine use of a recombinant C17A C3d construct has prohibited the structural and functional analysis of disulphide-linked dimers of C3 fragments that can form following activation in the fluid phase. In this study, we present the first X-ray crystal structure of a human C3d dimer where dimerisation is mediated by partial disulphide linkage of the thioester cysteine residues at position 17/1010 (C3d numbering/intact pre-pro C3 numbering) (**Figure 1**) in a manner that would also permit dimerisation of C3b (**Supplementary Figure S2e**). Importantly, this dimer retains the ability of C3d to bind SCR domains 1-2 of CR2, the α_M_I integrin domain of CR3 and SCR domains 19-20 of FH (**Supplementary Figures S2a-d**). We also find that in the presence of domain IV of the *S. aureus* immunomodulator Sbi, dimeric C3d undergoes N-terminal 3D domain swapping, exchanging the N-terminal α1 helices of the two C3d chains (**Figures 2** and **3**). This domain swap structurally stabilises the dimer as evidenced by the expansion in interaction surface, enhanced number of intermolecular interactions and increase in melting temperature (**Supplementary Figure S4**).

Intriguingly, the secondary binding site for Sbi-IV on the convex surface of C3d revealed previously through X-ray crystallographic, NMR chemical shift perturbation^15^ and SAXS analyses^8^, is located near the base or hinge region of the swapped C3d^17C^ α1 helix. It is at this position between helices α1 and α2 where Sbi-IV helix α1 forms strong interactions with the thioester cysteine C17/1010 and surrounding residues S15/1008 and Q20/1013 of C3d (**Supplementary Figure S3**^**15**^). Thus, this alternative binding of Sbi-IV to the hinge region of the N-terminal α1 helix of C3d, which in our structure is occluded by tight dimer interactions, could play an important role in the Sbi-IV-mediated helix swapping observed in the present study, particularly as helix α1 is prone to displacement, which has been shown to occur during the conversion of C3 to C3b^10^. Binding of Sbi-IV could further increase the susceptibility of this specific helix in C3d to displacement such that when two C3d monomers bound by Sbi-IV on their convex surfaces in this way are brought into close proximity, the helix α1-bound Sbi-IV molecules migrate to bind the concave face of the opposing C3d molecules, a favoured higher-affinity interaction, causing a reciprocal exchange of the C3d α1 helices followed by disulphide linkage of the thioester cysteine residues at position 17/1010 (**Figure 6**). Indeed, our near-UV thermal melt analyses, which suggest N-terminal truncation (V1_D11del) abolishes the ability of Sbi-IV to displace the N-terminal α1 helical region of C3d (**Supplementary Figure S5**), provide support for the proposed role of Sbi-IV interactions with the convex binding site of C3d in the molecular mechanism underlying the observed helix swap.

**Figure 6.**
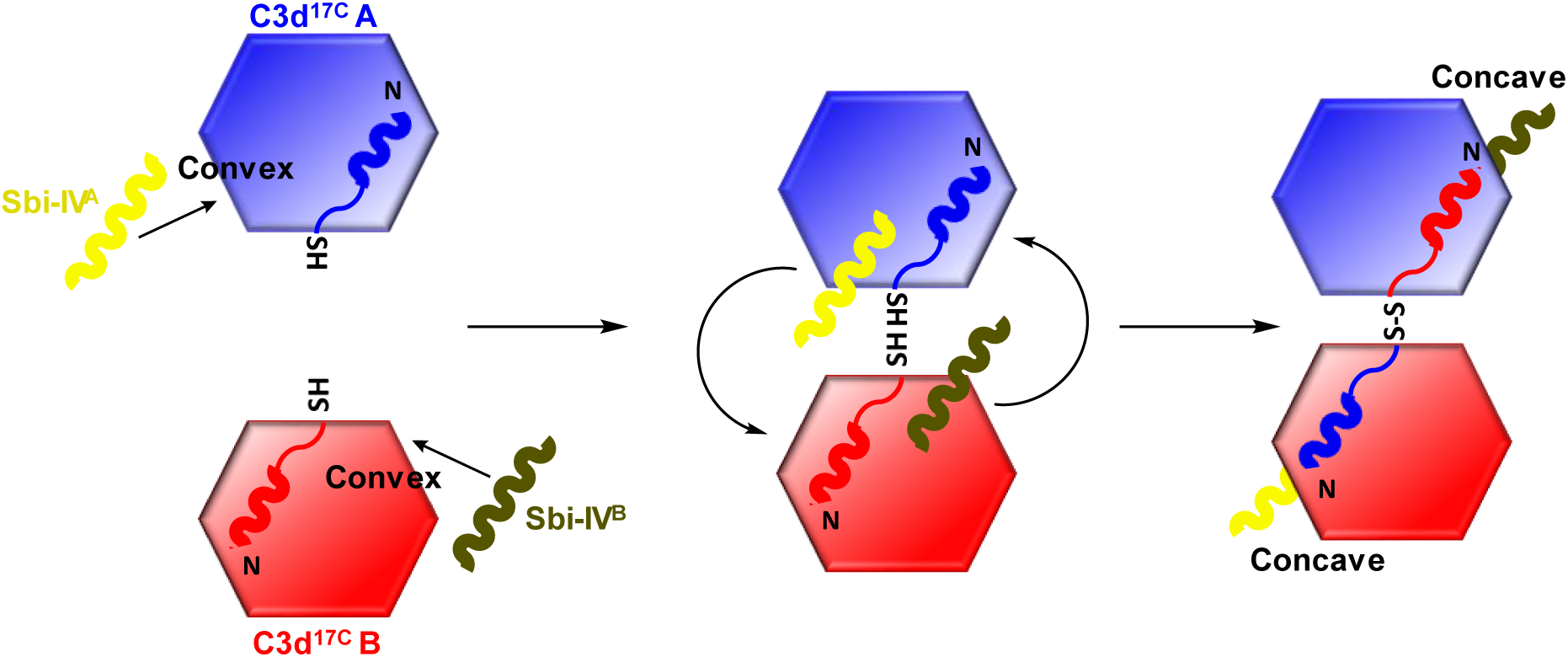
Schematic model depicting the proposed mechanistic basis behind Sbi-IV-induced N-terminal helix swapping of dimeric C3d^17C^. **Left**: in the fluid phase, Sbi-IV^A^ helix α1 binds to the N-terminal helix α1 and its adjoining loop on the convex thioester-containing face of C3d^17C^ monomer A (a short-distance low-affinity interaction) and similarly Sbi-IV^B^ binds the convex face of C3d^17C^B. **Middle and right**: when these two Sbi-IV-bound C3d^17C^ monomers come into close proximity, Sbi-IV^A^ helix α2 is attracted to and binds the concave face of C3d^17C^B (a more favourable higher affinity interaction), pulling C3d^17C^A helix α1 with it. A similar change in binding mode occurs with Sbi-IV^B^ binding the concave face of C3d^17C^A resulting in a reciprocal exchange of the N-terminal α1 helices of the two C3d^17C^ monomers. The switch in Sbi-IV binding mode from the convex to the concave surface of C3d releases the thioester domain allowing the formation of a disulphide bond between the thioester cysteines of the two C3d^17C^ monomers and hence the formation of a helix-swapped C3d dimer. For the purposes of simplicity only the N-terminal helix of C3d^17C^ and one Sbi-IV helix are shown.

Following the swap, the altered placement of the swapped helix α1 could enhance its susceptibility to further degradation and lead to the formation of C3dt, a more compact dimer, as suggested by the previously-published structure of a truncated rat dimer which lacks the exact region of C3d involved in helix swapping (M1/994 – G16/1009)^34^. On a broader perspective, this newly-discovered N-terminal helix swap of dimeric C3d is unique as this relatively rare structural event has not been described in C3d or a complement protein previously, although 3D domain swapping has been documented in a number of disparate proteins^40,43^ and various complement components such as FH-related proteins (FHRs)^44,45^ are known to exist in a dimeric form. To the best of our knowledge this is also the first report of a domain swap facilitated by the binding of a ligand.

In order to complement our structural studies, we next analysed the binding of a stable chemically-linked C3d^17C^ dimer (**Supplementary Figures S8** and **S9**) to CR2 (SCR1-4) and FH_19-20_ using SPR. Here dimeric C3d^17C^ showed higher avidity binding to both of the interacting partners examined, and in contrast to monomeric C3d, was found to crosslink surface-associated CR2 as well as FH_19-20_ (**Figure 4, Supplementary Figure S10** and **S11**). This crosslinking of CR2 by disulphide-linked C3d^17C^ dimers cannot be explained by the formation of higher order aggregates of dimeric C3d^17C^ (**Supplementary Figures S9c** and **S9d**) or FH_19-20_ (**Supplementary Figure S12**) and is a finding that has not been observed previously but could indicate a potential physiologically-relevant role of these dimers. Future investigations will elucidate whether dimeric C3d^17C^ can crosslink its other receptor, CR3, or a combination of CR2 and CR3, as suggested by our structural superpositions (**Supplementary Figures S2b** and **S2c**).

Finally, we investigated the effects of dimeric compared to monomeric C3d on the activation state of primary human and mouse B cells using flow cytometry and Ca^2+^ influx experiments. When assayed in isolation, B cells purified from human PBMCs appeared to be unresponsive to both forms of free C3d (**Supplementary Figure S13**). However, both monomeric C3d^17A^, and to a greater extent dimeric C3d^17C^, inhibited BCR/CR2-mediated Ca^2+^ influx in B220^+^ murine splenocytes when added prior to stimulation with a BCR/CR2-crosslinking complex (**Supplementary Figure S14**). Further to previous reports using biotinylated C3dg (with a C17A mutation) in the presence of streptavidin^46^, these results suggest that pre-ligation of CR2 by naturally-occurring fluid phase C3d(g) dimers could inhibit BCR/CR2 crosslinking-mediated Ca^2+^ responses in B cells by sequestering the CR2/CD19/CD81 receptor complex from the BCR with higher avidity than C3d monomers.

Both dimeric and monomeric C3d were also found to induce changes in the expression of B cell activation markers on human CD19^+^ cells within PBMC samples (**Figure 5, Supplementary Figure S17**). Specifically, in the presence of anti-IgM, both monomeric C3d^17A^ and to a three-fold greater extent dimeric C3d^17C^, synergistically upregulated CD69 and CD86 which is consistent with previous reports showing independent ligation of CR2 and the BCR (i.e. without crosslinking) by simultaneous stimulation with biotinylated-C3dg/streptavidin complexes and anti-IgM can augment B cell activation^47^. Our results, however, additionally show that dimeric C3d^17C^ is more efficient at augmenting CR2/BCR-dependent activation and that BCR engagement may not be necessary for upregulation of certain activation makers as at higher concentrations both forms of C3d were also capable of enhancing expression of the early activation markers CD69 and CD86 in the absence of anti-IgM.

In contrast to CD69 and CD86, both monomeric and dimeric C3d appeared to downregulate CD40, with a more pronounced reduction in expression in the presence of dimeric C3d^17C^. CD40 is involved in the regulation of several B cell processes including germinal centre reactions^48^, isotype switching^49^ and somatic hypermutation^50^ and has also been shown to prevent B cells from becoming anergic^51^. In line with these roles, downregulation of CD40 was shown to be a beneficial outcome of Rituximab treatment of systemic lupus erythematosus (SLE) patients^52^ and CD40/CD40L levels have been linked to anti-DNA autoantibody titres in lupus patients^53^ and mouse models^54^. Although further investigations are required to explain the C3d-mediated downregulation of CD40 expression on B cells observed in our study, it is possible that C3d stimulation of CR2 or CR3 expressed on other PBMC cell types (e.g. T cells^55-58^ and natural killer cells^59^) induces the production of higher levels of soluble CD40L that drive internalisation of CD40 or prevent efficient staining by occluding the receptor. Alternatively, the known binding of CD40L to CR3^60^ could be outcompeted by CR3 interactions with C3d, particularly in its dimeric form, elevating the levels of soluble CD40L available for binding CD40. Further experiments investigating the effects of free monomeric and dimeric C3d on IgG titre and hence B-cell differentiation or antibody class switching following activation of PBMCs with T cell supernatants or co-culture with IL-2 or CD40L-producing feeder cells will help to understand this process further.

On the whole, our cell experiments not only suggest free fluid-phase C3d(g) (unattached to a surface) may regulate B cell activation in a selective manner but also that there are clear functional differences between monomeric and dimeric C3d with the latter being a more potent modulator of the activation state of B cells as a consequence of high avidity receptor interactions or through receptor crosslinking. In addition, they indicate other PBMC cell types play an important role in the responsiveness of B cells, in terms of the activation markers analysed, to C3d, perhaps through the provision of sensitising or synergising co-stimulatory molecules or via CR2-CR2 or CR2-CR3 crosslinking between cells.

Together with the fact that C3d inhibited BCR/CR2-mediated Ca^2+^ influx prior to stimulation with a BCR/CR2-crosslinking complex, the C3d-induced downregulation of CD40 expression could also signify fluid-phase C3d(g), particularly in its dimeric form, may alter the activation of B cells and direct them towards an anergic state. This would be logical in terms of helping to limit the involvement of complement in the generation of humoral immune responses in the absence of a threat and is consistent with reports of CR2 ligation being involved in the anergy of autoreactive B cells^47,61,62^. Thus, C3d(g) dimers could have implications for the development of novel therapies for autoimmune diseases through their effects on CD40/CD40L interactions which along with B cell anergy, are also critical for the maintenance of peripheral self-tolerance by immature dendritic cells (DCs)^63^, known to express CR3^64,65^. By extension, these newly-uncovered functions of C3d could also offer a possible explanation as to why humoral immune responses are inhibited, rather than enhanced, by certain vaccine constructs composed of antigens linked to linear repeats of C3d placed in close proximity to each other^47,66^.

If free C3d(g), more notably in its dimeric form, does in fact play a role in anergising or inducing a state of tolerance in B cells, then in addition to sequestering CR2 away from *S. aureus* surfaces and antigens, the consumptive cleavage of C3 in the fluid phase followed by stabilisation of disulphide-linked C3d dimers through domain swapping mediated by Sbi could promote B cell anergy, as well more widespread tolerance through possible effects on T cells and DCs, and thereby further facilitate evasion of the immune system by *S. aureus*. Future investigations could elucidate whether these interactions between Sbi and C3d and their resultant functional consequences are involved in the known association between *S. aureus* infections and autoimmune diseases^67^. In a wider context, it would also be interesting to explore possible connections between the levels of C3d(g) dimers, their distribution in the body and pathological conditions associated with uncontrolled C3 activation, such as C3 glomerulopathy, as we surmise local upregulation of fluid phase C3d(g) concentrations is likely to enhance C3d(g) dimerisation.

In summary, in this study we present the first structures of fluid-phase disulphide-linked human C3d dimers and through accompanying functional analyses show that these dimers have physiologically-relevant roles in crosslinking CR2 and selectively modulating B cell activation, possibly to trigger tolerogenic pathways, which could have implications for autoimmune diseases. In addition, our new discovery that Sbi domain IV can induce structurally-stabilising domain swapping of dimeric C3d, a unique example in the field of structural biology of the complement system, reveals another dimension in the complex relationship between *S. aureus* and the immune system and could signify a novel staphylococcal B cell subversion strategy. Overall, our findings shed light on a fundamental aspect of complement biology that is often overlooked and have the potential to inform the design of novel therapeutics for autoimmune diseases in the future.

## Materials and Methods

### Expression and purification of recombinant proteins

The DNA sequence of human C3d (residues 1-310) comprised of C3 residues 996-1303 (pre-pro C3 numbering) with a Cys to Ala mutation at position 17(C3d) /1010 (pre-pro C3) (C3d^17A^) was previously cloned into the pET15b expression plasmid^12^. To reproduce the wild-type sequence, the Ala at position 17 of the C3d^17A^ construct was reverted back to a Cys (C3d^17C^) using site-directed mutagenesis. Both C3d constructs were expressed in the *Escherichia coli* BL21(DE3) (Sigma Aldrich) or Shuffle T7 (NEB) cell lines and purified using cation exchange followed by size exclusion chromatography.

The DNA sequence of the Sbi construct composed of domain IV (150-266 of full-length Sbi) bearing an N-terminal 6x His-tag was previously cloned into the pQE30 plasmid^15^ while the DNA sequence of a truncated Sbi-IV construct lacking 11 residues located at the N-terminus of helix α1 (198-208 of full-length Sbi) (V1_D11del) was produced using PCR with designed primers and cloned into the pET28a plasmid. Both Sbi constructs were expressed in *E. coli* BL21(DE3) cells and purified using nickel-affinity and size exclusion chromatography.

The human CR2(SCR1-4)-Fc and FH_19-20_ constructs used in surface plasmon resonance experiments were expressed in Chinese hamster ovary (CHO) cells or *Pichia pastoris* respectively and purified as described previously (CR2(SCR1-4)-Fc: ^7^; FH_19-20_: ^68^). The monomeric state of FH_19-20_ was confirmed using analytical ultracentrifugation (**Supplementary Figure S12**).

### Crystallisation, data collection and structure determination of dimeric C3d^17C^ and a C3d^17C^ dimer-Sbi-IV complex

Crystallisation was performed at 18°C using the hanging drop vapour diffusion method. For the dimeric C3d^17C^ structure, a 15 mg mL^-1^ (432 µM) C3d^17C^ solution was subjected to a grid screen containing 100 mM Tris pH 8, 50-300 mM NaCl and 16-26% PEG 4000. Crystals appeared in the condition containing 100 mM Tris pH 8, 200 mM NaCl, 24% PEG 4000. For the C3d^17C^ dimer-Sbi-IV complex structure, needle-like co-crystals at a 1:1 molar (288 µM) ratio of Sbi-IV (2.66 mg mL^-1^) and C3d^17C^ (10 mg mL^-1^) appeared in the condition containing 0.2 M Sodium citrate tribasic dihydrate, 20% PEG 3350 (PACT *premier* condition E11, Molecular Dimensions) (measured pH: 8.07).

Crystals were mounted on loops, flash-frozen in liquid nitrogen and X-ray diffraction data collected on the IO4 beamline at the Diamond Light Source synchrotron (Oxfordshire, UK) (See **Supplementary Table S1** for data collection statistics). Integration of Dectris PILATUS 6M pixel detector diffraction images and data reduction were performed using Xia2-DIALS and AIMLESS respectively. The automated BALBES pipeline and COOT were used for molecular replacement and model building. Refinement was carried out in REFMAC and Phenix (refinement statistics can be found in **Supplementary Table S2**) and UCSF Chimera was used for superpositioning and generation of images. Bond numbers and buried surface area were calculated with PDBePISA. Both structures have been submitted to the PDB with the following submission codes: 6RMT (C3d^17C^ dimer) and 6RMU (C3d^17C^ dimer-Sbi-IV complex).

### Circular dichroism (CD) spectroscopy

For tertiary structure thermal melt analyses, buffer-corrected CD measurements of 87.5 µM C3d^17C^ or C3d^17A^ alone or in the presence of an equimolar concentration of wild-type Sbi-IV or an Sbi-IV truncation mutant (V1_D11del) in 10 mM phosphate pH 7.4 were acquired at wavelengths between 250 and 320 nm in 1 nm increments. 50 μg mL^−1^ C3d^17C^ or C3d^17A^ and wavelengths between 190 and 280 nm were used for secondary structure analyses. All measurements were obtained on a Chirascan spectrometer (Applied Photophysics) with a bandwidth of 2 nm and 2 second time per point at temperatures ranging from 5°C to 85°C in 5°C increments followed by a final reading at 20°C. Millidegree values were converted to units of mean residue ellipticity and deconvolution of secondary structural data was performed using DichroWeb. Melting temperature values were estimated using sigmoidal fits of thermal denaturation curves at different wavelengths performed using the Boltzmann function in GraphPad Prism (version 8.4.1). See **Supplementary Figures S4** and **S5** for near-UV thermal melt analysis of C3d^17C^ and C3d^17A^ and sigmoidal fits of thermal denaturation curves used for Tm calculations and **Supplementary Figure S6** for far-UV thermal melt studies of C3d^17C^ and C3d^17A^.

### Synthesis and characterisation of N,N’-(propane-1,3-diyl) bis(2-bromoacetamide) linker

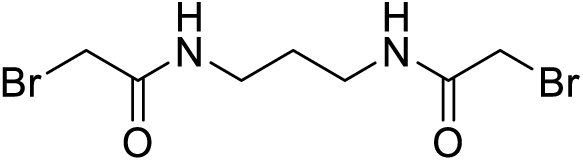

A solution of K_2_CO_3_ (5.92 g, 42.8 mmol) in water (21 mL) was added to a solution of 1,3-diaminopropane (1.06 g, 14.3 mmol) in chloroform (35 mL) at 5°C with stirring. A solution of bromoacetyl bromide (8.65 g, 42.8 mmol) in anhydrous chloroform (15 mL) was then added dropwise to the mixture and the reaction was left stirring at room temperature for 18 hours. The resultant precipitate was filtered, washed with water (6 x 10 mL), and dried under vacuum to yield *N,N’*-(propane-1,3-diyl)bis(2-bromoacetamide) as a white solid (2.45 g, 55%). Subsequent characterisation of the linker was performed using ^1^H-NMR (**Supplementary Figure S7a**) and ^13^C-NMR (**Supplementary Figure S7b**). High resolution electrospray ionisation time-of-flight mass spectrometry m/z: [M + Na]^+^ calculated for C_7_H_12_Br_2_N_2_O_2_Na = 338.9143 Da, 338.9143 Da was observed.

### Production, purification and characterisation of chemically-linked C3d dimers

For the generation of chemically-linked C3d^17C^ dimers, small-scale trials were performed involving combination of C3d^17C^ with the *N,N’*-(propane-1,3-diyl)bis(2-bromoacetamide) linker in 0.1 M Tris, 0.15 M NaCl, 5 mM EDTA pH 7.5 at 0.55, 0.75 or 1.0 molar equivalences. Following overnight incubation at room temperature (21°C), linker-mediated C3d^17C^ dimerisation was confirmed using reducing SDS-PAGE and electrospray time-of-flight mass spectrometry (**Supplementary Figure S8**). A larger scale reaction at 0.75 molar equivalence (3.75 mg C3d^17C^, 0.026 mg linker) was subsequently carried out as described above and subjected to size exclusion chromatography to separate the chemically-linked dimeric C3d^17C^ from monomeric C3d^17C^ (**Supplementary Figures S9a** and **S9b**). Particle size analysis yielded a single species (**Supplementary Figure S9c**), analytical ultracentrifugation confirmed the dimeric state of the chemically-linked C3d^17C^ (**Supplementary Figure S9d**) and both biophysical techniques showed a lack of aggregate formation. Chemically-linked dimeric C3d^17C^ was subsequently digested with trypsin (1:50 ratio) at 37°C over a time course (**Supplementary Figure S9e**). The digestion reaction was stopped by addition of a trypsin inhibitor (1:2 ratio). Electrospray ionization time-of-flight mass spectrometry of the trypsin-digested dimeric C3d^17C^ fragments followed by analysis using the Masshunter Qualitative Analysis and BioConfirm (Agilent) software packages was used to confirm chemical linkage at position 17C of C3d (**Supplementary Table S3**) and the presence of an intact internal disulphide bond (**Supplementary Table S4**).

### Surface plasmon resonance

All surface plasmon resonance experiments were performed on a Biacore S200 sensor (GE Healthcare) at 25°C with HBST (10 mM HEPES, 150 mM NaCl, 0.005% Tween-20, pH 7.4) used as the running buffer. CR2-Fc and FH_19-20_ were prepared in 10 mM sodium acetate pH 5 and immobilised at 300 RU (CR2-Fc: 20 µM, FH_19-20_: 240 µM) to different flow cells of CM5 chips (GE Healthcare) using standard amine coupling involving preparation of the dextran matrix with 100 mM N-hydroxysuccinimide (NHS) and 40 mM 1-ethyl-3-(dimethylaminopropyl) carbodiimide (EDC) followed by quenching with 1 M ethanolamine-HCl pH 8.5. Monomeric C3d^17A^ and chemically-linked dimeric C3d^17C^ used as analytes were prepared to a fixed concentration, serially diluted in HBST and injected in duplicate. 10 mM sodium acetate, 1 M NaCl pH 4 was used as the regeneration buffer but could not regenerate the chip surface of the highly avid interactions between dimeric C3d^17C^ and CR2-Fc/FH_19-20_. Data were analysed using the Biacore S200 Evaluation Software 1.0. Responses from blank reference flow cells were subtracted from ligand-immobilised flow cells and all data were double-referenced (buffer inject subtracted).

### Flow cytometric analysis of B cell activation

Frozen human peripheral blood mononuclear cells (PBMCs) from blood donors were thawed and diluted into cold RPMI 1640 medium (Gibco) supplemented with 10% (v/v) foetal bovine serum, 1% (v/v) penicillin-streptomycin (Sigma Aldrich) and 1% (v/v) GlutaMAX (Gibco). For experiments on isolated B cells, B cells were purified from PBMCs by negative selection using the Miltenyi Biotec Human B Cell Isolation Kit II as per the manufacturer’s instructions. Following centrifugation, cells were counted, assessed for viability, which typically exceeded 90% for PBMCs and 80% for B cells, and re-suspended to the desired density in ambient medium.

PBMCs or B cells were then seeded onto sterile V-bottom plates at a density of 150,000 or 40,000 cells/well respectively and allowed to recover at 37°C with 5% CO_2_ for 1h. Monomeric C3d^17A^ or chemically-linked dimeric C3d^17C^ were serially diluted in media and added to the cells in duplicate to give final concentrations ranging from 2 µM to 0.1 nM (based on the molecular weight of monomeric C3d^17A^ for both constructs in order to control for the number of binding sites). Following a 30 min incubation period with the C3d constructs, additions of either goat F(ab’)_2_ anti-human IgM LE/AF (Southern Biotech) at a final concentration of 10 µg mL^-1^ or media were made and the cells incubated for a further 18h at 37°C with 5% CO_2_.

After a period of cooling on ice, the cells were stained for 1h with the LIVE/DEAD™ fixable near-infrared dead cell stain (1:1000 dilution, Invitrogen) along with the following labelled antibodies diluted in an ice-cold staining buffer (PBS supplemented with 1% BSA, 2mM EDTA and 0.05% NaN_3_): PerCP-Cy™5.5 mouse anti-human CD19 (1:40 dilution, Clone HIB19, BD Pharmingen) (for PBMC samples only), FITC mouse anti-human CD40 (1:20 dilution, Clone 5C3, BD Pharmingen), Brilliant Violet 421™ mouse anti-human CD69 (1:40 dilution, Clone FN50, BioLegend), PE mouse anti-human CD71 (1:20 dilution, Clone M-A712, BD Pharmingen) and APC mouse anti-human CD86 (1:20 dilution, Clone 2331, BD Pharmingen). The cells were subsequently washed and analysed on an Intellicyt® iQue Screener PLUS flow cytometer. The gating strategy applied for live, singlet CD19^+^ B cells and activation markers can be found in **Supplementary Figures S15** and **S16**, respectively. Antibody capture beads were used for compensation. Data were expressed as mean values from at least 2 replicates ± standard deviation from the mean and depicted as scatter plots with curves fitted using a four parameter variable slope non-linear regression model in GraphPad Prism (version 8.4.1).

### Ca^2+^ influx experiments

Intracellular Ca^2+^ measurements using flow cytometry were performed as described previously^46,69,70^. Briefly, isolated C57BL/6 mouse splenocytes maintained at 37°C were Indo 1-AM loaded, stained with a rat anti-mouse CD45R/B220-APC antibody (Clone RA3-6B2, BD Pharmingen) and analysed on a BD LSR II flow cytometer (BD Biosciences) at RT. 4 or 10 µg of monomeric C3d^17A^ or chemically-linked dimeric C3d^17C^ were added to the cells 30s after data acquisition. After 90s, cells were stimulated with a suboptimal concentration of pre-mixed complexes composed of 0.056 µg mL^-1^ biotinylated F(ab’)_2_ goat anti-mouse IgM (µ-chain specific) (Jackson ImmunoResearch), ∼3 µg mL^-1^ C3dg-biotin (produced in house) and ∼1.3 µg mL^-1^ streptavidin (aIgM-b/C3dg-b/ST). Experiments were run for 10 min and intracellular Ca^2+^ influx of gated B220^+^ cells was analysed using the FlowJo® software (FlowJo LLC, BD).

## Data Availability

The datasets generated and analysed during the current study are available from the corresponding authors on request.

## Acknowledgements

This research was supported by the Biotechnology and Biological Sciences Research Council Follow On Fund BB/N022165/1. A.A.W. was sponsored by a PhD studentship granted by Raoul and Catherine Hughes and the University of Bath Alumni Fund. R.W.D. was supported by a Medical Research Council GW4 Doctoral Training Partnership. B.G. and K.J.M. were funded by a Northern Counties Kidney Research Grant and Alexion Pharmaceuticals funded T.M.H.’s PhD studentship via Complement UK. K.J.M. also recognises the support from Kidney Research UK grant (RP-006-20270301). C.L.H. was funded by Newcastle University. The authors would like to acknowledge the team at the Diamond Light Source synchrotron (Oxfordshire, UK) for access to the IO4 beamline. Shaun Reeksting from the Material and Chemical Characterisation Facility (MC^2^, University of Bath) is also thanked for his assistance with the mass spectrometry analyses.

## Author Contributions

J.v.d.E, K.J.M., A.M., B.G.G. and A.A.W. designed the experiments. C.R.B. performed preliminary structural studies that lead to the discovery of the helix swap. A.A.W. performed the crystallisation and circular dichroism experiments. S.J.C. and A.A.W. reprocessed the crystallography data and refined the structures. R.W.D. produced the truncated Sbi-IV mutant and carried out the structural analyses and molecular modelling. A.G.W., B.A. and R.L.M. synthesised and characterised the linker and conducted initial linkage experiments. B.G.G. and A.A.W. performed the SPR experiments under the guidance of K.J.M. and with helpful discussions from C.L.H. who also analysed the data. L.K. completed the Ca^2+^ influx experiments. The B cell activation flow cytometry experiments were performed by A.M. and K.W. with the assistance of I.P.M. J.v.d.E. and A.A.W. wrote the manuscript with valuable contributions from all the authors.

## Competing interests

K.J.M. is a consultant for and receives funding or renumeration from Gemini Therapeutics Ltd., Freeline Therapeutics, MPM Capital and Catalyst Biosciences. C.L.H. has recently received consultancy or SAB payments from Freeline Therapeutics, Q32 Bio Inc., Roche, GlaxoSmithKline and Gyroscope Therapeutics and has received research funding from Ra Pharmaceuticals; all funds were donated to Newcastle University. T.M.H. is funded by Alexion Pharmaceuticals. All other authors declare no competing interests.

## Additional information

**Supplementary information** for this publication is available online

## References

1. Carter, R. & Fearon, D. Polymeric C3dg Primes Human B Lymphocytes for Proliferation Induced By Anti-IgM. J. Immunol. 143, 1755–1760 (1989).

2. Dempsey, P., Allison, M., Akkaraju, S., Goodnow, C. & Fearon, D. C3d of complement as a molecular adjuvant: Bridging innate and acquired immunity. Science 271, 348–350, doi:10.1126/science.271.5247.348 (1996).

3. Ross, T., Xu, Y., Bright, R. & Robinson, H. C3d enhancement of antibodies to hemagglutinin accelerates protection against influenza virus challenge. Nat. Immunol. 1, 127–131, doi:10.1038/77802 (2000).

4. Green, T. et al. C3d enhancement of neutralizing, antibodies to measles hemagglutinin. Vaccine 20, 242–248, doi:10.1016/S0264-410X(01)00266-3 (2001).

5. Henson, S., Smith, D., Boackle, S., Holers, V. & Karp, D. Generation of recombinant human C3dg tetramers for the analysis of CD21 binding and function. J. Immunol. Methods 258, 97–109, doi:10.1016/S0022-1759(01)00471-9 (2001).

6. Green, T. D., Montefiori, D. C. & Ross, T. M. Enhancement of antibodies to the human immunodeficiency virus type 1 envelope by using the molecular adjuvant C3d. J. Virol. 77, 2046–2055, doi:10.1128/Jvi.77.3.2046-2055.2003 (2003).

7. He, Y. et al. A novel C3d-containing oligomeric vaccine proVides insight into the viability of testing human C3d-based vaccines in mice. Immunobiology 223, 125–134, doi:10.1016/j.imbio.2017.10.002 (2017).

8. Yang, Y. et al. Utilization of Staphylococcal Immune Evasion Protein Sbi as a Novel Vaccine Adjuvant. Front. Immunol. 9, doi:10.3389/fimmu.2018.03139 (2019).

9. Janssen, B. et al. Structures of complement component C3 provide insights into the function and evolution of immunity. Nature 437, 505–511, doi:10.1038/nature04005 (2005).

10. Janssen, B., Christodoulidou, A., McCarthy, A., Lambris, J. & Gros, P. Structure of C3b reveals conformational changes that underlie complement activity. Nature 444, 213–216, doi:10.1038/nature05172 (2006).

11. Xue, X. et al. Regulator-dependent mechanisms of C3b processing by factor I allow differentiation of immune responses. Nat. Struct. Mol. Biol. 24, 643–651, doi:10.1038/nsmb.3427 (2017).

12. Nagar, B., Jones, R., Diefenbach, R., Isenman, D. & Rini, J. X-ray crystal structure of C3d: A C3 fragment and ligand for complement receptor 2. Science 280, 1277–1281, doi:10.1126/science.280.5367.1277 (1998).

13. van den Elsen, J. & Isenman, D. A Crystal Structure of the Complex Between Human Complement Receptor 2 and Its Ligand C3d. Science 332, 608–611, doi:10.1126/science.1201954 (2011).

14. Vorup-Jensen, T. & Jensen, R. Structural Immunology of Complement Receptors 3 and 4. Front. Immunol. 9, doi:10.3389/fimmu.2018.02716 (2018).

15. Clark, E. et al. A structural basis for Staphylococcal complement subversion: X-ray structure of the complement-binding domain of Staphylococcus aureus protein Sbi in complex with ligand C3d. Mol. Immunol. 48, 452–462, doi:10.1016/j.molimm.2010.09.017 (2011).

16. Hammel, M. et al. A structural basis for complement inhibition by Staphylococcus aureus. Nat. Immunol. 8, 430–437, doi:10.1038/ni1450 (2007).

17. Ricklin, D., Ricklin-Lichtsteiner, S. K., Markiewski, M. M., Geisbrecht, B. V. & Lambris, J. D. Cutting Edge: Members of the Staphylococcus aureus Extracellular Fibrinogen-Binding Protein Family Inhibit the Interaction of C3d with Complement Receptor 2. J. Immunol. 181, 7463–7467, doi:10.4049/jimmunol.181.11.7463 (2008).

18. Hammel, M. et al. Characterization of Ehp, a secreted complement inhibitory protein from Staphylococcus aureus. J. Biol. Chem. 282, 30051–30061, doi:10.1074/jbc.M704247200 (2007).

19. Amdahl, H. et al. Staphylococcal Ecb Protein and Host Complement Regulator Factor H Enhance Functions of Each Other in Bacterial Immune Evasion. J. Immunol. 191, 1775–1784, doi:10.4049/jimmunol.1300638 (2013).

20. Kajander, T. et al. Dual interaction of factor H with C3d and glycosaminoglycans in host-nonhost discrimination by complement. Proc. Natl. Acad. Sci. USA 108, 2897–2902, doi:10.1073/pnas.1017087108 (2011).

21. Morgan, H. et al. Structural basis for engagement by complement factor H of C3b on a self surface. Nat. Struct. Mol. Biol. 18, 463–470, doi:10.1038/nsmb.2018 (2011).

22. Law, S. K. A. & Dodds, A. W. The internal thioester and the covalent binding properties of the complement proteins C3 and C4. Protein Sci. 6, 263–274, doi:10.1002/pro.5560060201 (1997).

23. Bexborn, F., Andersson, P., Chen, H., Nilsson, B. & Ekdahl, K. The tick-over theory revisited: Formation and regulation of the soluble alternative complement C3 convertase (C3(H_2_O)Bb). Mol. Immunol. 45, 2370–2379, doi:10.1016/j.molimm.2007.11.003 (2008).

24. Perkins, S. J. & Sim, R. B. Molecular Modeling of Human-Complement Component-C3 and Its Fragments by Solution Scattering. Eur. J. Biochem. 157, 155–168, doi:10.1111/j.1432-1033.1986.tb09652.x (1986).

25. Rodriguez, E., Nan, R., Li, K., Gor, J. & Perkins, S. A Revised Mechanism for the Activation of Complement C3 to C3b - A Molecular Explanation of a Disease-Associated Polymorphism. J. Biol. Chem. 290, 2334–2350, doi:10.1074/jbc.M114.605691 (2015).

26. Arnaout, M., Melamed, J., Tack, B. & Colten, H. Characterization of the Human Complement (C3b) Receptor with a Fluid Phase C3b Dimer. J. Immunol. 127, 1348–1354 (1981).

27. Melamed, J., Arnaout, M. & Colten, H. Complement (C3b) Interaction with the Human Granulocyte Receptor - Correlation of Binding of Fluid-Phase Radiolabeled Ligand with Histaminase Release. J. Immunol. 128, 2313–2318 (1982).

28. Kinoshita, T. et al. C5 Convertase of the Alternative Complement Pathway - Covalent Linkage Between 2 C3b Molecules within the Trimolecular Complex Enzyme. J. Immunol. 141, 3895–3901 (1988).

29. Hong, K. et al. Reconstitution of C5 Convertase of the Alternative Complement Pathway with Isolated C3b Dimer and Factors B and D. J. Immunol. 146, 1868–1873 (1991).

30. Jelezarova, E., Luginbuehl, A. & Lutz, H. C3b_2_-IgG complexes retain dimeric C3 fragments at all levels of inactivation. J. Biol. Chem. 278, 51806–51812, doi:10.1074/jbc.M304613200 (2003).

31. Shigeoka, A., Gobel, R., Janatova, J. & Hill, H. Neutrophil Mobilization Induced by Complement Fragments during Experimental Group-B Streptococcal (GBS) Infection. Am. J. Pathol. 133, 623–629 (1988).

32. Gilbert, H., Eaton, J., Hannan, J., Holers, V. & Perkins, S. Solution structure of the complex between CR2 SCR 1-2 and C3d of human complement: An X-ray scattering and sedimentation modelling study. J. Mol. Biol. 346, 859–873, doi:10.1016/j.jmb.2004.12.006 (2005).

33. Li, K. et al. Solution Structure of the Complex Formed between Human Complement C3d and Full-length Complement Receptor Type 2. J. Mol. Biol. 384, 137–150, doi:10.1016/j.jmb.2008.08.084 (2008).

34. Zanotti, G. et al. Structure at 1.44 angstrom resolution of an N-terminally truncated form of the rat serum complement C3d fragment. Biochim. Biophys. Acta, Protein Struct. Mol. Enzymol. 1478, 232–238, doi:10.1016/S0167-4838(00)00040-6 (2000).

35. Isenman, D. E. & van den Elsen, J. M. H. in Structural Biology of the Complement System (eds D. Morikis & J.D. Lambris) Ch. 5, 111-142 (CRC Press, 2005).

36. Forneris, F. et al. Structures of C3b in Complex with Factors B and D Give Insight into Complement Convertase Formation. Science 330, 1816–1820, doi:10.1126/science.1195821 (2010).

37. Forneris, F. et al. Regulators of complement activity mediate inhibitory mechanisms through a common C3b-binding mode. EMBO J. 35, 1133–1149, doi:10.15252/embj.201593673 (2016).

38. Upadhyay, A. et al. Structure-function analysis of the C3 binding region of Staphylococcus aureus immune subversion protein Sbi. J. Biol. Chem. 283, 22113–22120, doi:10.1074/jbc.M802636200 (2008).

39. Burman, J. et al. Interaction of human complement with Sbi, a staphylococcal immunoglobulin-binding protein - Indications of a novel mechanism of complement evasion by Staphylococcus aureus. J. Biol. Chem. 283, 17579–17593, doi:10.1074/jbc.M800265200 (2008).

40. Liu, Y. & Eisenberg, D. 3D domain swapping: As domains continue to swap. Protein Sci. 11, 1285–1299, doi:10.1110/ps.0201402 (2002).

41. Kelly, S. M. & Price, N. C. The Use of Circular Dichroism in the Investigation of Protein Structure and Function. Curr. Protein Pept. Sc. 1, 349–384, doi: 10.2174/1389203003381315 (2000).

42. Yang, P. et al. Engineering a long-acting, potent GLP-1 analog for microstructure-based transdermal delivery. Proc. Natl. Acad. Sci. USA 113, 4140–4145, doi:10.1073/pnas.1601653113 (2016).

43. Rousseau, F., Schymkowitz, J. & Itzhaki, L. S. in Protein Dimerization and Oligomerization in Biology (ed Jacqueline M. Matthews) 137–152 (Springer New York, 2012).

44. de Jorge, E. et al. Dimerization of complement factor H-related proteins modulates complement activation in vivo. Proc. Natl. Acad. Sci. USA 110, 4685–4690, doi:10.1073/pnas.1219260110 (2013).

45. van Beek, A. et al. Factor H-Related (FHR)-1 and FHR-2 Form Homo- and Heterodimers, while FHR-5 Circulates Only As Homodimer in Human Plasma. Front. Immunol. 8, doi:10.3389/fimmu.2017.01328 (2017).

46. Lyubchenko, T., dal Porto, J., Cambier, J. C. & Holers, V. M. Coligation of the B cell receptor with complement receptor type 2 (CR2/CD21) using its natural ligand C3dg: Activation without engagement of an inhibitory signaling pathway. J. Immunol. 174, 3264–3272, doi:10.4049/jimmunol.174.6.3264 (2005).

47. Lee, Y. et al. Complement component C3d-antigen complexes can either augment or inhibit B lymphocyte activation and humoral immunity in mice depending on the degree of CD21/CD19 complex engagement. J. Immunol. 175, 8011–8023, doi:10.4049/jimmunol.175.12.8011 (2005).

48. Han, S. H. et al. Cellular Interaction in Germinal Centers - Roles of CD40 Ligand and B7-2 in Established Germinal Centers. J. Immunol. 155, 556–567 (1995).

49. Aversa, G., Punnonen, J., Carballido, J., Cocks, B. & de Vries, J. CD40 ligand- CD40 interaction in Ig isotype switching in mature and immature human B cells. Semin. Immunol. 6, 295–301, doi:10.1006/smim.1994.1038 (1994).

50. Zan, H. et al. Induction of Ig somatic hypermutation and class switching in a human monoclonal IgM^+^ IgD^+^ B cell line in vitro: Definition of the requirements and modalities of hypermutation. J. Immunol. 162, 3437–3447 (1999).

51. Eris, J. M. et al. Anergic Self-Reactive B-Cells Present Self Antigen and Respond Normally to Cd40-Dependent T-Cell Signals but Are Defective in Antigen-Receptor-Mediated Functions. Proc. Natl. Acad. Sci. USA 91, 4392–4396, doi:10.1073/pnas.91.10.4392 (1994).

52. Tokunaga, M. et al. Down-regulation of CD40 and CD80 on B cells in patients with life-threatening systemic lupus erythematosus after successful treatment with rituximab. Rheumatology 44, 176–182, doi: 10.1093/rheumatology/keh443 (2005).

53. Desai-Mehta, A., Lu, L. J., Ramsey-Goldman, R. & Datta, S. K. Hyperexpression of CD40 ligand by B and T cells in human lupus and its role in pathogenic autoantibody production. J. Clin. Invest. 97, 2063–2073, doi:10.1172/JCI118643 (1996).

54. Kaneko, Y. et al. CD-40-mediated stimulated of B1 and B2 cells: Implication in autoantibody production in murine lupus. Eur. J. Immunol. 26, 3061–3065, doi:10.1002/eji.1830261236 (1996).

55. Tsoukas, C. D. & Lambris, J. D. Expression of CR2/EBV Receptors on Human Thymocytes Detected by Monoclonal-Antibodies. Eur. J. Immunol. 18, 1299–1302, doi:10.1002/eji.1830180823 (1988).

56. Fischer, E., Delibrias, C. & Kazatchkine, M. Expression of CR2 (the C3dg/EBV receptor, CD21) on normal human peripheral blood T lymphocytes. J. Immunol. 146, 865–869 (1991).

57. Levy, E. et al. Lymphocyte-T Expression of Complement Receptor-2 (Cr-2 Cd21) - a Role in Adhesive Cell Cell-Interactions and Dysregulation in a Patient with Systemic Lupus-Erythematosus (Sle). Clin. Exp. Immunol. 90, 235–244, doi:10.1111/j.1365-2249.1992.tb07935.x (1992).

58. Wagner, C. et al. The complement receptor 3, CR3 (CD11b/CD18), on T lymphocytes: activation-dependent up-regulation and regulatory function. Eur. J. Immunol. 31, 1173–1180, doi:10.1002/1521-4141(200104)31:4<1173::aid- immu1173^#x003E;3.0.co;2-9 (2001).

59. Min, X. Y. et al. Expression and regulation of complement receptors by human natural killer cells. Immunobiology 219, 671–679, doi:10.1016/j.imbio.2014.03.018 (2014).

60. Zirlik, A. et al. CD40 ligand mediates inflammation independently of CD40 by interaction with Mac-1. Circulation 115, 1571–1580, doi:10.1161/CIRCULATIONAHA.106.683201 (2007).

61. Prodeus, A. P. et al. A critical role for complement in maintenance of self-tolerance. Immunity 9, 721–731, doi:10.1016/S1074-7613(00)80669-X (1998).

62. Birrell, L., Kulik, L., Morgan, B. P., Holers, V. M. & Marchbank, K. J. B cells from mice prematurely expressing human complement receptor type 2 are unresponsive to T-dependent antigens. J. Immunol. 174, 6974–6982, doi: 10.4049/jimmunol.174.11.6974 (2005).

63. Quezada, S. A., Jarvinen, L. Z., Lind, E. E. & Noelle, R. J. CD40/CD154 interactions at the interface of tolerance and immunity. Annu. Rev. Immunol. 22, 307–328, doi:10.1146/annurev.immunol.22.012703.104533 (2004).

64. Verbovetski, I. et al. Opsonization of apoptotic cells by autologous iC3b facilitates clearance by immature dendritic cells, down-regulates DR and CD86, and up-regulates CC chemokine receptor 7. J. Exp. Med. 196, 1553–1561, doi: 10.1084/jem.20020263 (2002).

65. Bajtay, Z., Speth, C., Erdei, A. & Dierich, M. P. Cutting edge: Productive HIV-1 infection of dendritic cells via complement receptor type 3 (CR3, CD11b/CD18). J. Immunol. 173, 4775–4778, doi:10.4049/jimmunol.173.8.4775 (2004).

66. Suradhat, S. et al. Fusion of C3d molecule with bovine rotavirus VP7 or bovine herpesvirus type 1 glycoprotein D inhibits immune responses following DNA immunization. Vet Immunol. Immunop. 83, 79–92, doi: 10.1016/s0165-2427(01)00369-5 (2001).

67. Mousavi, M. N. S. et al. The pathogenesis of Staphylococcus aureus in autoimmune diseases. Microb Pathog. 111, 503–507, doi:10.1016/j.micpath.2017.09.028 (2017).

68. Herbert, A. P., Uhrin, D., Lyon, M., Pangburn, M. K. & Barlow, P. N. Disease-associated sequence variations congregate in a polyanion recognition patch on human factor H revealed in three-dimensional structure. J. Biol. Chem. 281, 16512–16520, doi:10.1074/jbc.M513611200 (2006).

69. Kulik, L. et al. Intrinsic B cell hypo-responsiveness in mice prematurely expressing human CR2/CD21 during B cell development. Eur. J. Immunol. 37, 623–633, doi:10.1002/eji.200636248 (2007).

70. Kulik, L., Chen, K. A., Huber, B. T. & Holers, V. M. Human complement receptor type 2 (CR2/CD21) transgenic mice provide an in vivo model to study immunoregulatory effects of receptor antagonists. Mol. Immunol. 48, 883–894, doi:10.1016/j.molimm.2010.12.019 (2011).

