## Supplementary Information for "Insights into the role of C3d dimers in B cell activation and Staphylococcal immune evasion"

This PDF file includes:

Supplementary Figures S1-S17

Supplementary Tables S1-S4

**Contents of Supplementary Information, Wahid *et al*.,:**

***Insights into the role of C3d dimers in B cell activation and Staphylococcal immune evasion.***


**Supplementary Figures**

**Supplementary Figure S1** **$C3d^{17C}$ dimers exist in equilibrium with monomers in solution and cannot be separated by size exclusion chromatography.**

**Supplementary Figure S2** **Superpositions of the $C3d^{17C}$ dimer with ligand-binding domains of CR2 (a ,c), CR3 (b, c), FH (d) and with C3b (e).**

**Supplementary Figure S3** **Two binding modes of Sbi-IV to C3d.**

**Supplementary Figure S4** **Sbi-IV-mediated helix swapping stabilises dimeric $C3d^{17C}$ in solution.**

**Supplementary Figure S5** **Near-UV thermal melt circular dichroism spectroscopic studies of $C3d^{17A}$ tertiary structure in the absence or presence of wild-type Sbi-IV or a truncated Sbi-IV mutant.**

**Supplementary Figure S6** **Far-UV circular dichroism spectroscopic studies of $C3d^{17C}$ (a) and $C3d^{17A}$ (b) secondary structure as a function of temperature.**

**Supplementary Figure S7** **Characterisation of *N,N'*-(propane-1,3-diyl) bis(2-bromoacetamide) linker.**

**Supplementary Figure S8** **Induced dimerisation of $C3d^{17C}$ using chemical linkage.**

**Supplementary Figure S9** **Purification and characterisation of chemically-linked $C3d^{17C}$ dimers.**

**Supplementary Figure S10** **Analysis of $C3d^{17C}$ dimer-CR2-Fc sensorgram.**

**Supplementary Figure S11** **SPR sensorgrams of $C3d^{17C}$ dimer binding interactions with immobilised CR2-Fc and $FH_{19\text{-}20}$.**

**Supplementary Figure S12** **Analytical ultracentrifugation analysis of $FH_{19\text{-}20}$.**

**Supplementary Figure S13** **Flow cytometric analysis of C3d-mediated changes in the activation state of purified B cells.**

**Supplementary Figure S14** **Preligation of CR2 with monomeric $C3d^{17A}$ or dimeric $C3d^{17C}$ inhibits BCR/CR2-dependent $Ca^{2+}$ influx in murine B cells.**

**Supplementary Figure S15** **Flow cytometry gating strategy for B cells in PBMC samples.**

| | |
|---|---|
| **Supplementary Figure S16** | **Flow cytometry channel gating of B cell activation markers.** |
| **Supplementary Figure S17** | **Flow cytometric analysis of C3d-mediated changes in the activation state of PBMC B cell populations from an additional two donors.** |

**<u>Supplementary Tables</u>**

| | |
|---|---|
| **Supplementary Table S1** | **Data collection statistics.** |
| **Supplementary Table S2** | **Refinement statistics.** |
| **Supplementary Table S3** | **Mass spectrometry species of trypsin-digested dimeric C3d$^{17C}$ confirming chemical linkage of C3d$^{17C}$ at position 17C.** |
| **Supplementary Table S4** | **Trypsin digest mass spectrometry data confirming presence of an intact internal disulphide bond in chemically-linked dimeric C3d$^{17C}$.** |

#### Supplementary Figure S1

a

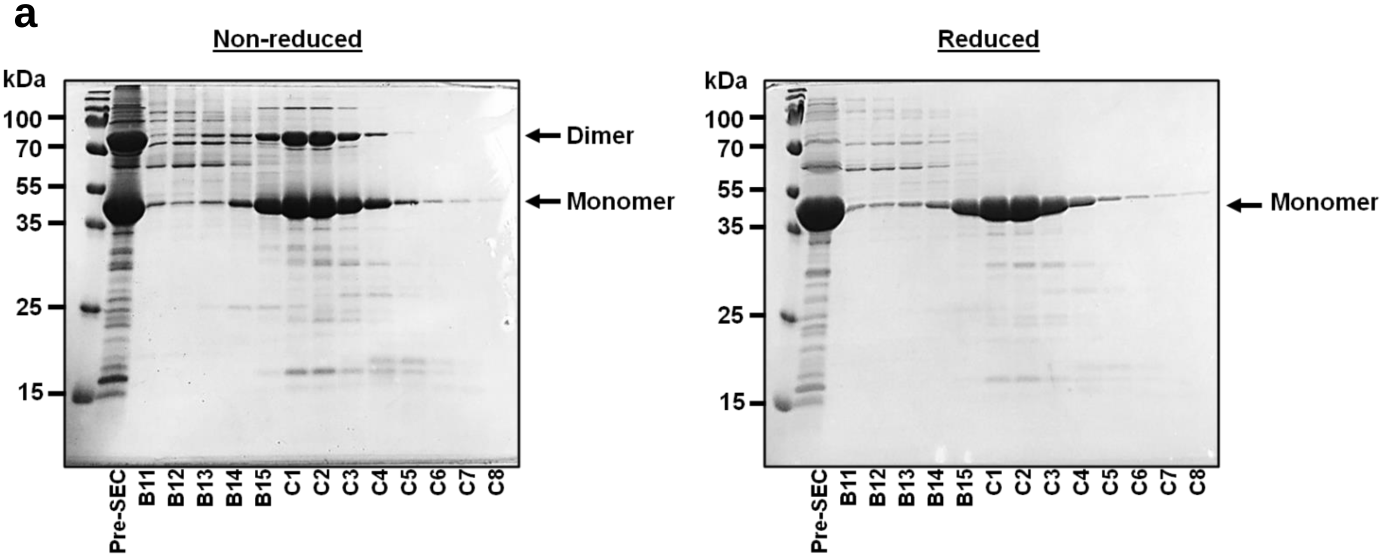


b

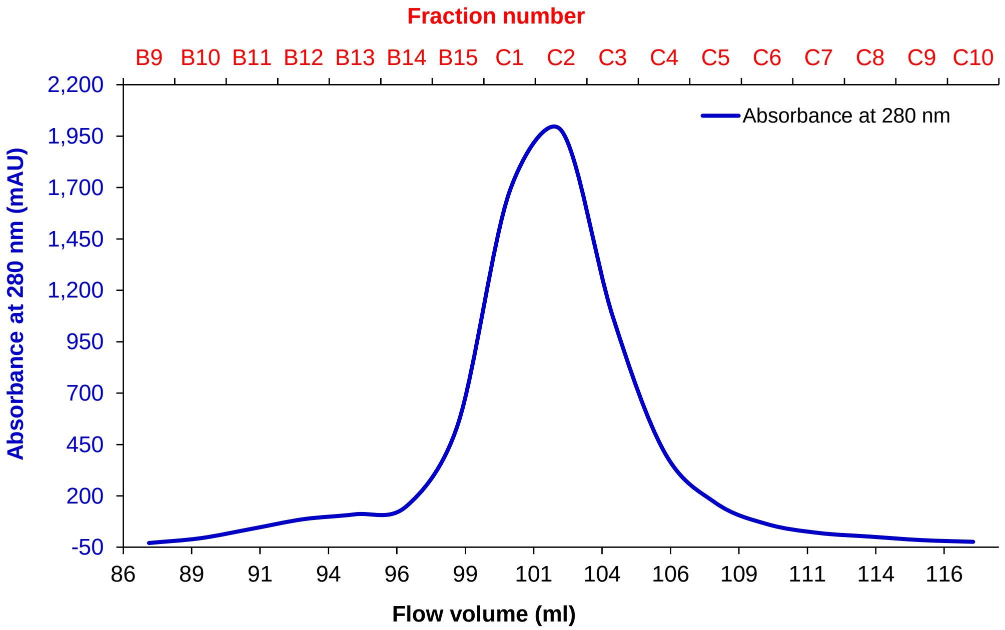


**C3d$^{17C}$ dimers exist in equilibrium with monomers in solution and cannot be separated by size exclusion chromatography. a:** non-reducing (L) and reducing (R) SDS-PAGE of elution fractions from C3d$^{17C}$ size exclusion chromatography. Elution of monomeric and dimeric C3d$^{17C}$ occurs simultaneously suggesting a monomer-dimer equilibrium in solution (L). Disulphide-linked dimers are converted to monomeric C3d$^{17C}$ in the presence of β-mercaptoethanol (R). **b**: Chromatogram showing elution peak of C3d$^{17C}$ size exclusion chromatography. Pre-SEC: pre-size exclusion chromatography.

#### Supplementary Figure S2

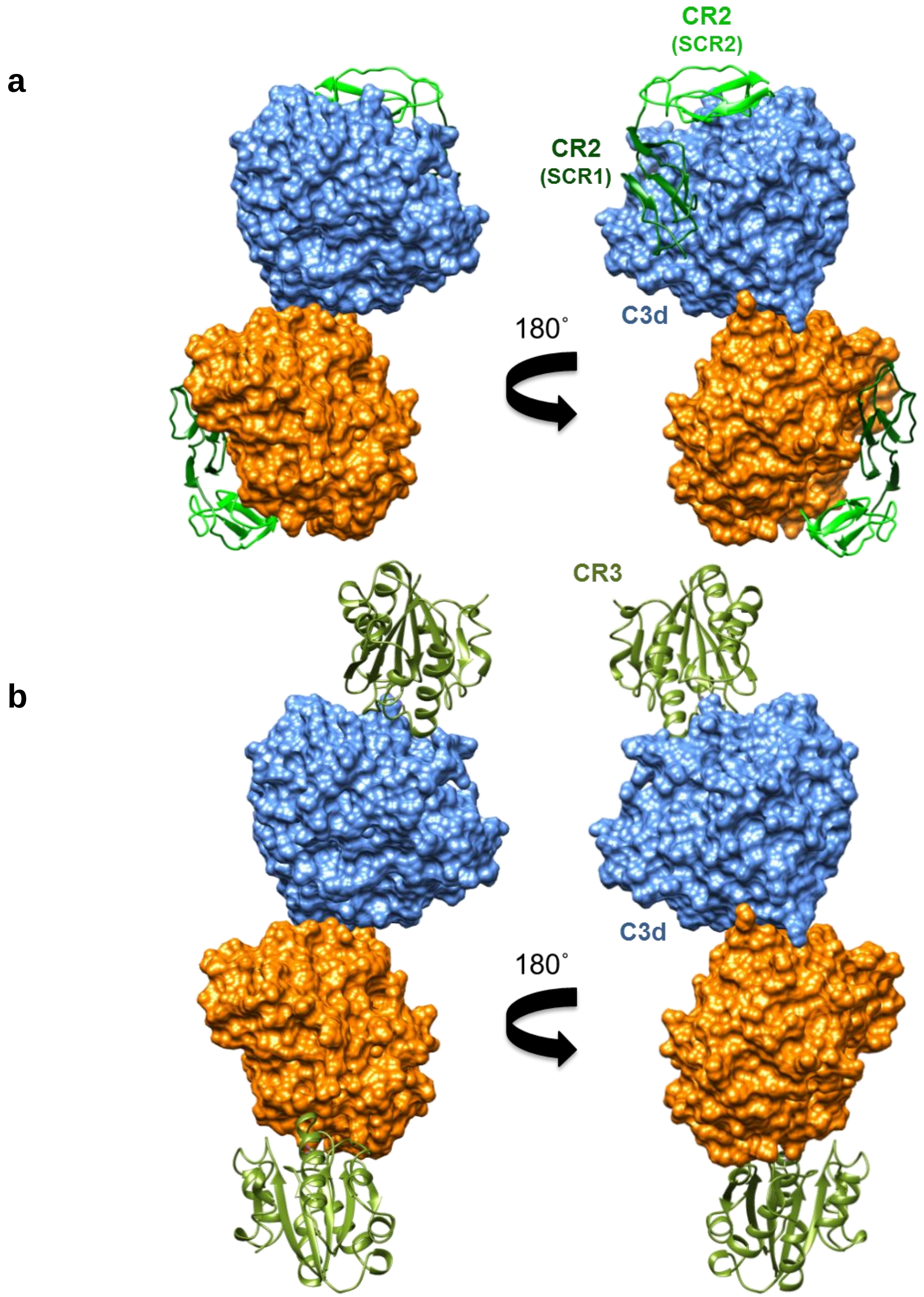


**Superpositions of the C3d$^{17C}$ dimer with ligand-binding domains of CR2 (**a, c**), CR3 (**b, c**), FH (**d**) and with C3b (**e**). a, b:** Superpositioning of C3d-binding domains SCR1-2 of CR2 (PDB accession code: 3OED) (**a**) and $\alpha_M$I integrin domain of CR3 (PDB accession code: 4M76) (**b**) onto their opposing binding sites in the C3d$^{17C}$ dimer.

**c**

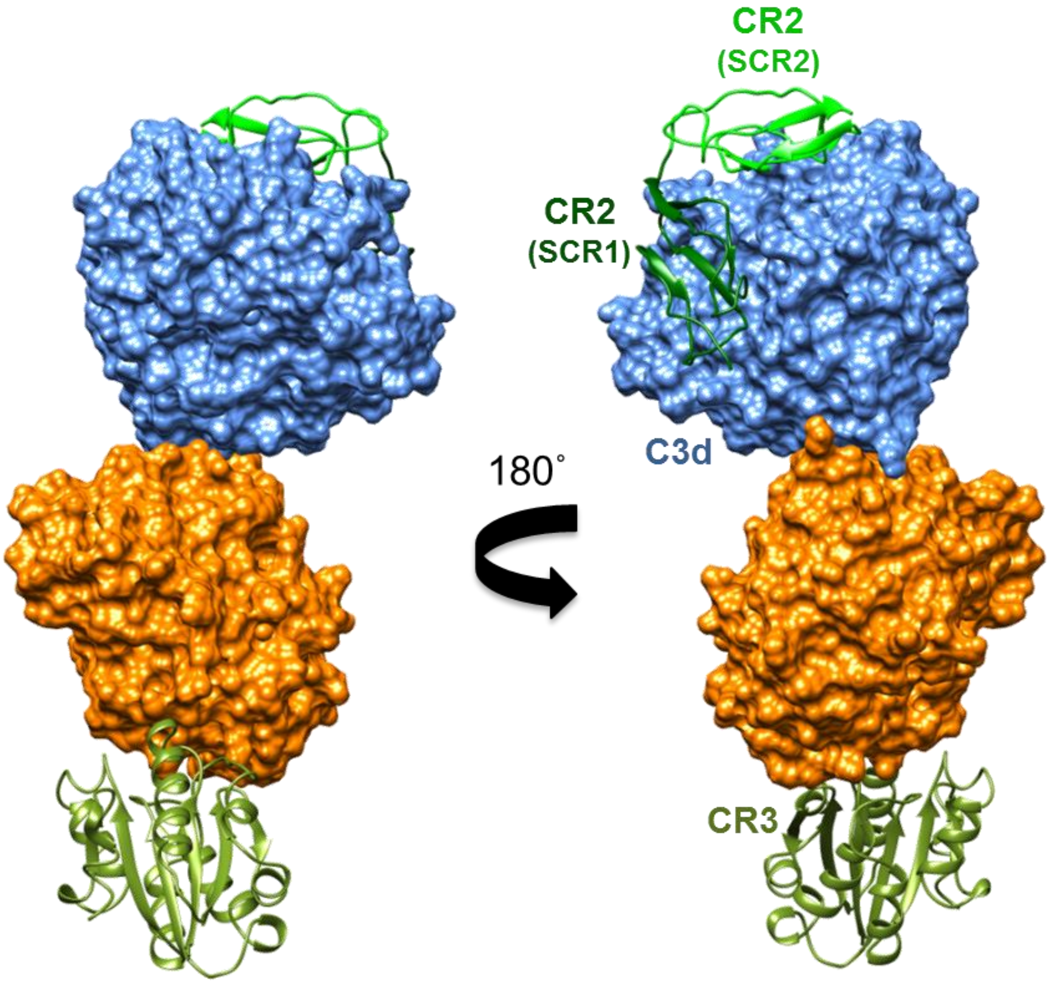


**c:** Superpositioning of CR2 SCR1-2 (PDB accession code: 3OED) and CR3 αMI integrin domain (PDB accession code: 4M76) onto the opposing concave and convex binding sites of the same C3d[17C] dimer indicating dimeric C3d could play a role in crosslinking of the receptors.

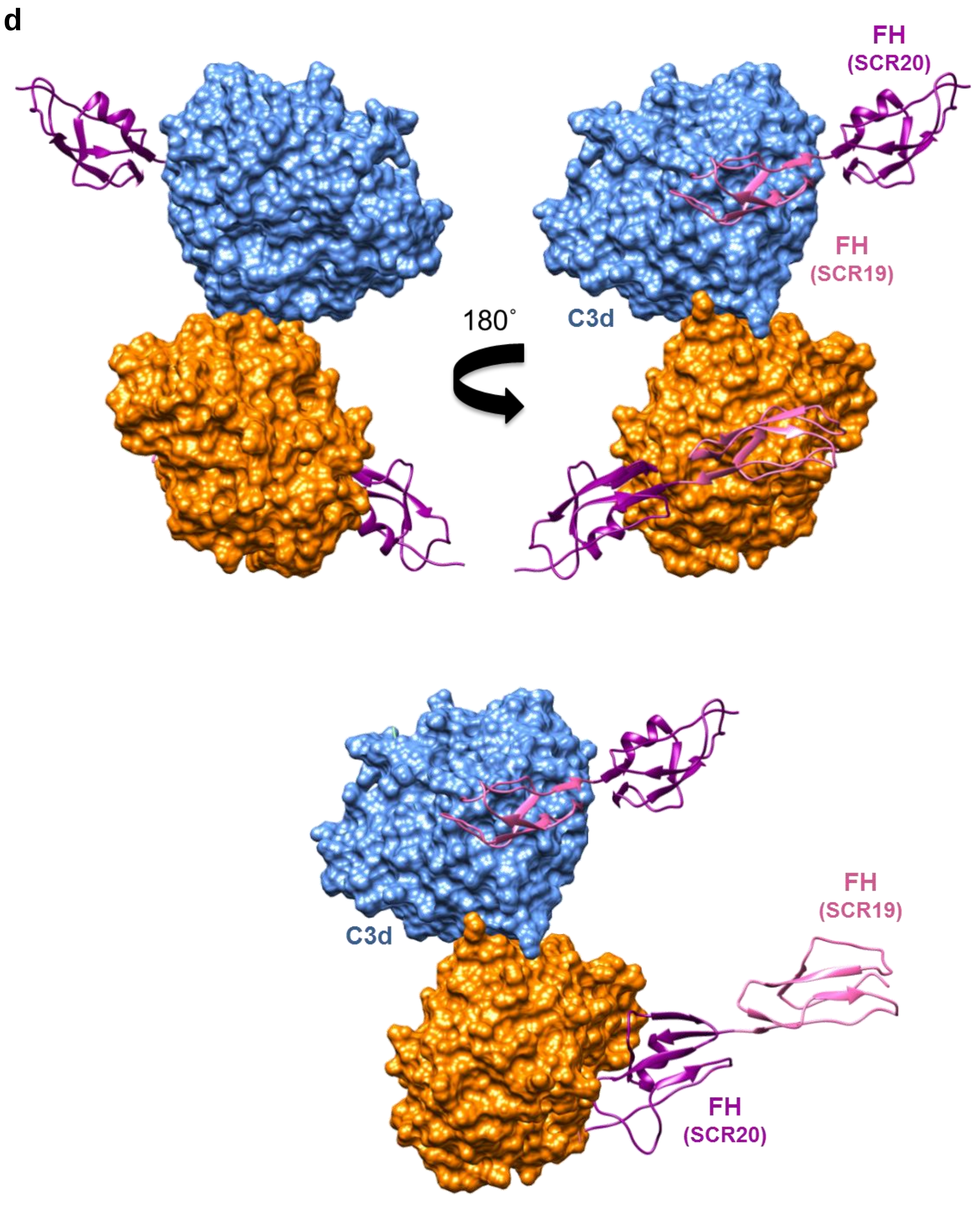


**d:** Superpositioning of C3d-binding domains (SCR19-20) of FH (PDB accession code: 2XQW) onto the C3d$^{17C}$ dimer. Shown are the interactions via SCR domain 19 (top, bottom) as well as SCR 20 (bottom).

**e**

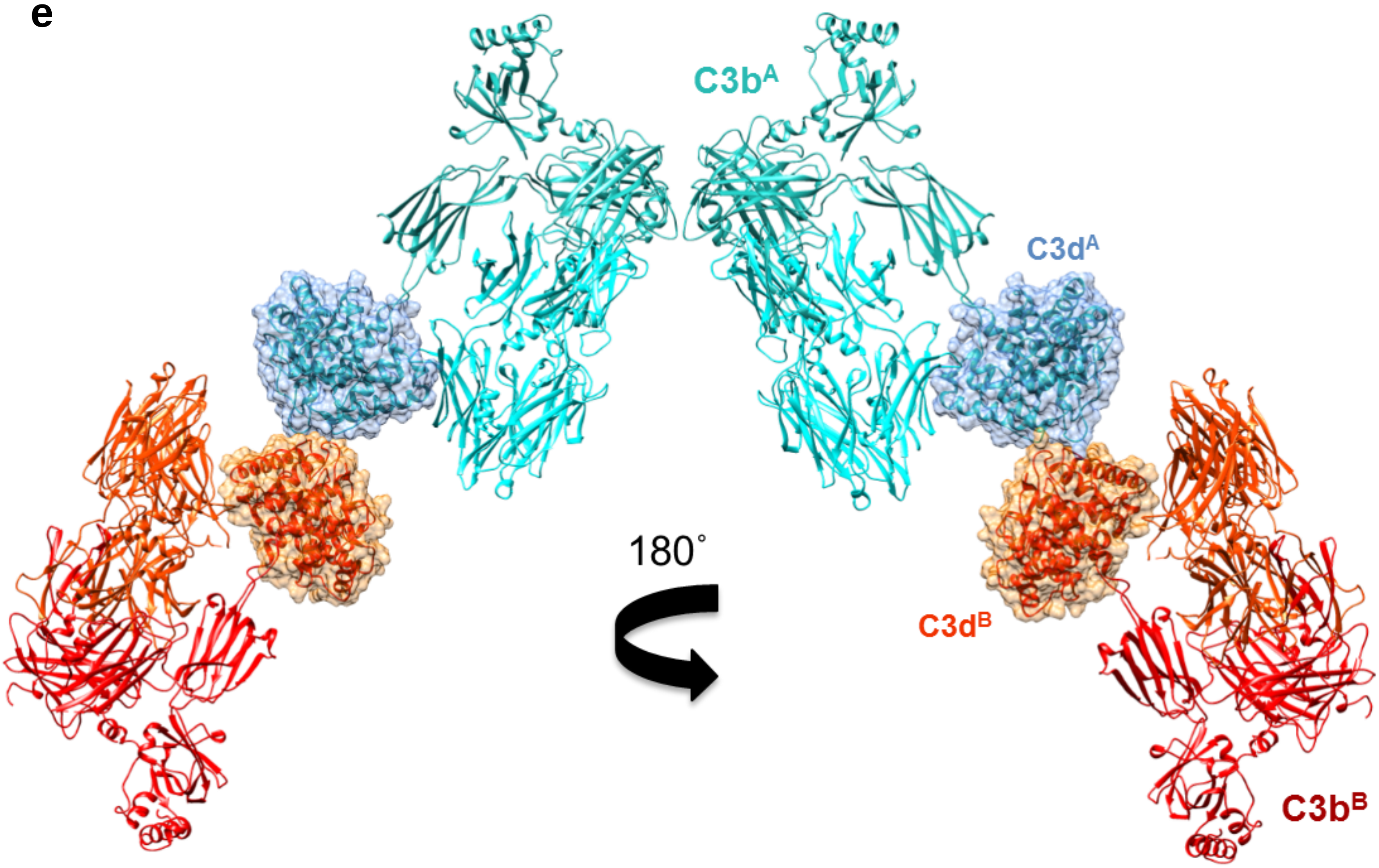


**e:** Superpositioning of the C3d$^{17C}$ dimer onto the structure of C3b (PDB accession code: 2WII) showing the potential of C3b dimer formation.

#### Supplementary Figure S3

**Two binding modes of Sbi-IV to C3d.** Shown are the two binding modes of Sbi-IV observed in the previously published crystal structure of the Sbi-IV:C3d$^{17A}$ complex (Clark *et al.*, 2010). Shown in red is Sbi-IV bound to the concave surface of C3d (in green) and in turquoise Sbi-IV bound to the convex surface of C3d (respective PDB accession codes: 2WY8 and 2WY7). The inset shows a close-up of the interactions of Sbi-IV with the hinge region at the base of the swapped α1 helix in the C3d$^{17C}$ dimer-Sbi-IV complex (adapted from Clark *et al.*, 2010).

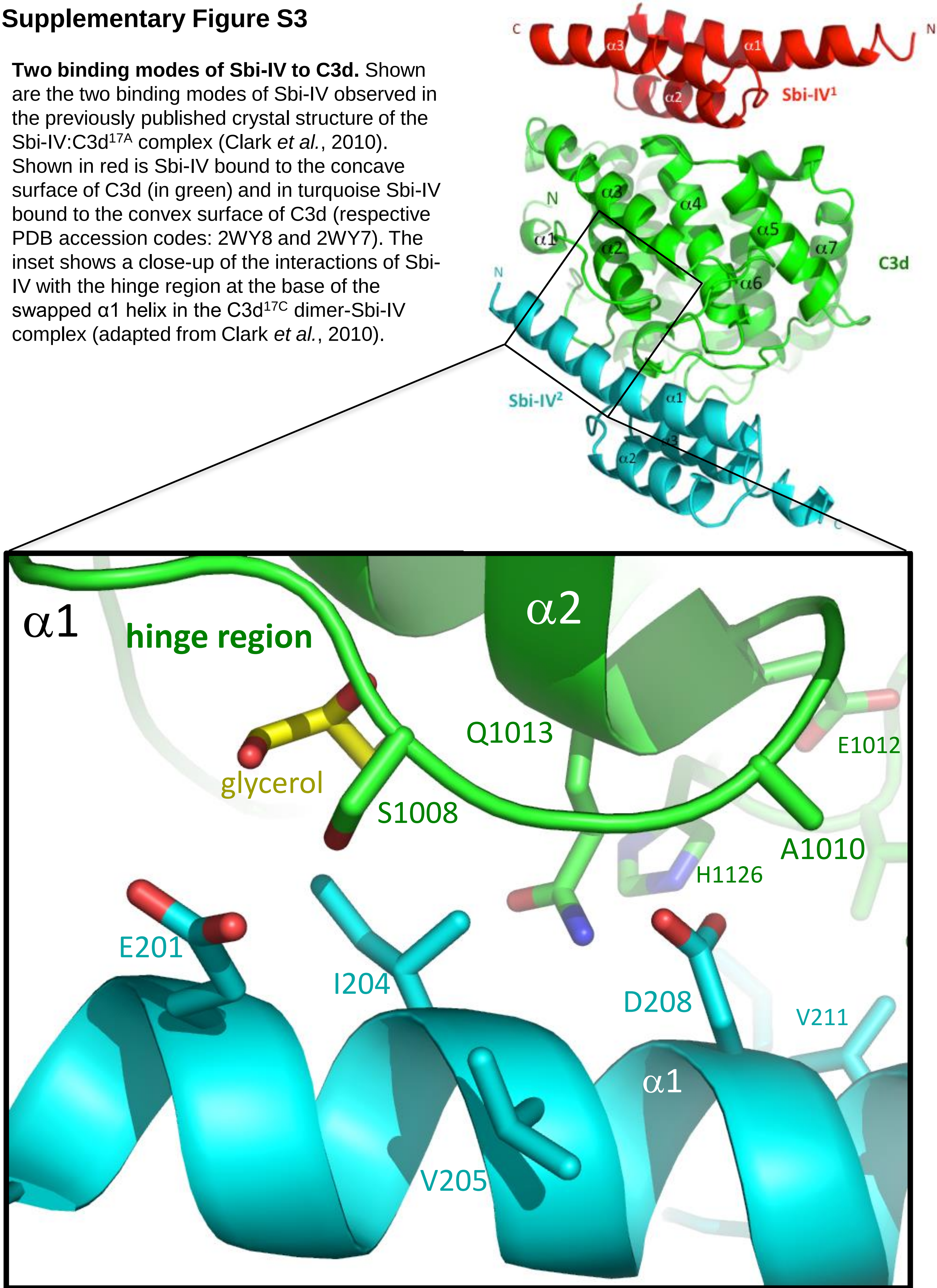

#### Supplementary Figure S4

**a**

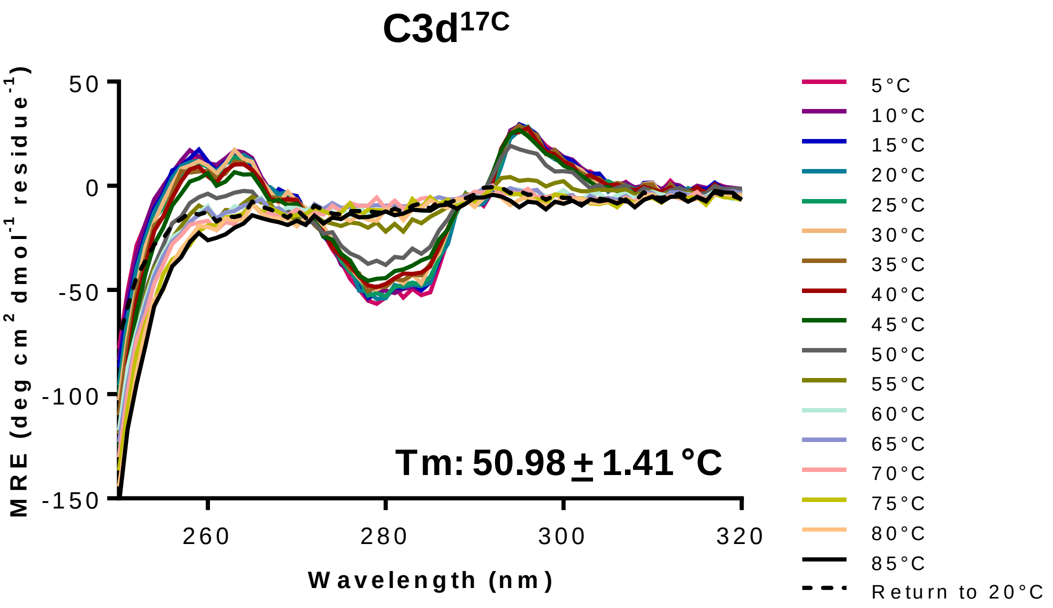


**b**

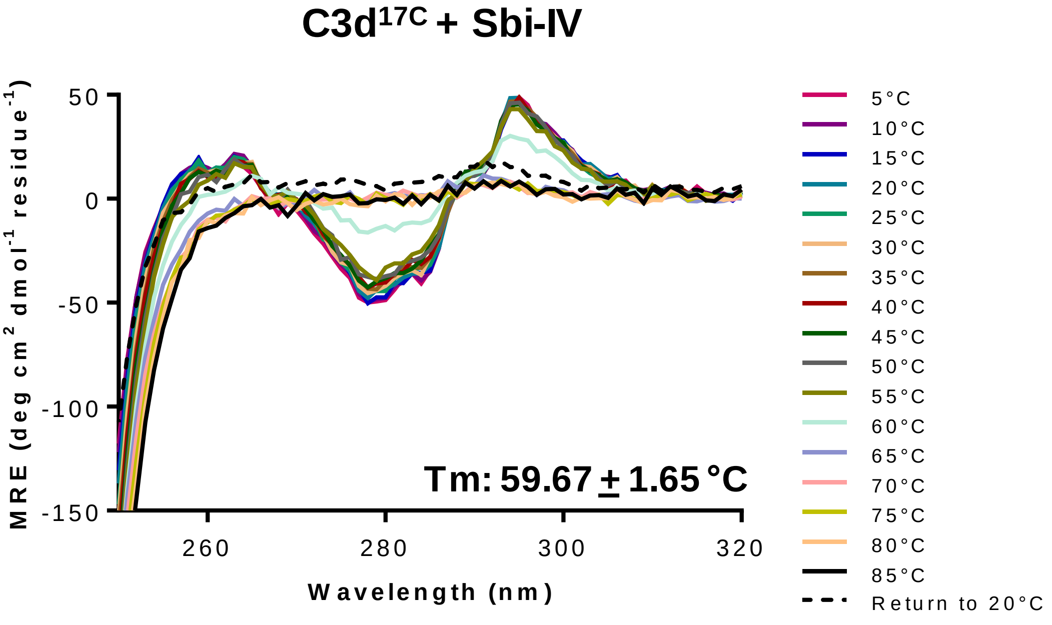


**Sbi-IV-mediated helix swapping stabilises dimeric C3d$^{17C}$ in solution.** Near-UV thermal melt circular dichroism spectroscopic studies of C3d$^{17C}$ in the absence (**a**) or presence (**b**) of Sbi-IV showing changes in tertiary structure as a function of temperature. Stabilisation of C3d$^{17C}$ in the presence of Sbi-IV is indicated by a ~8.7°C increase in melting temperature (Tm).

**c**

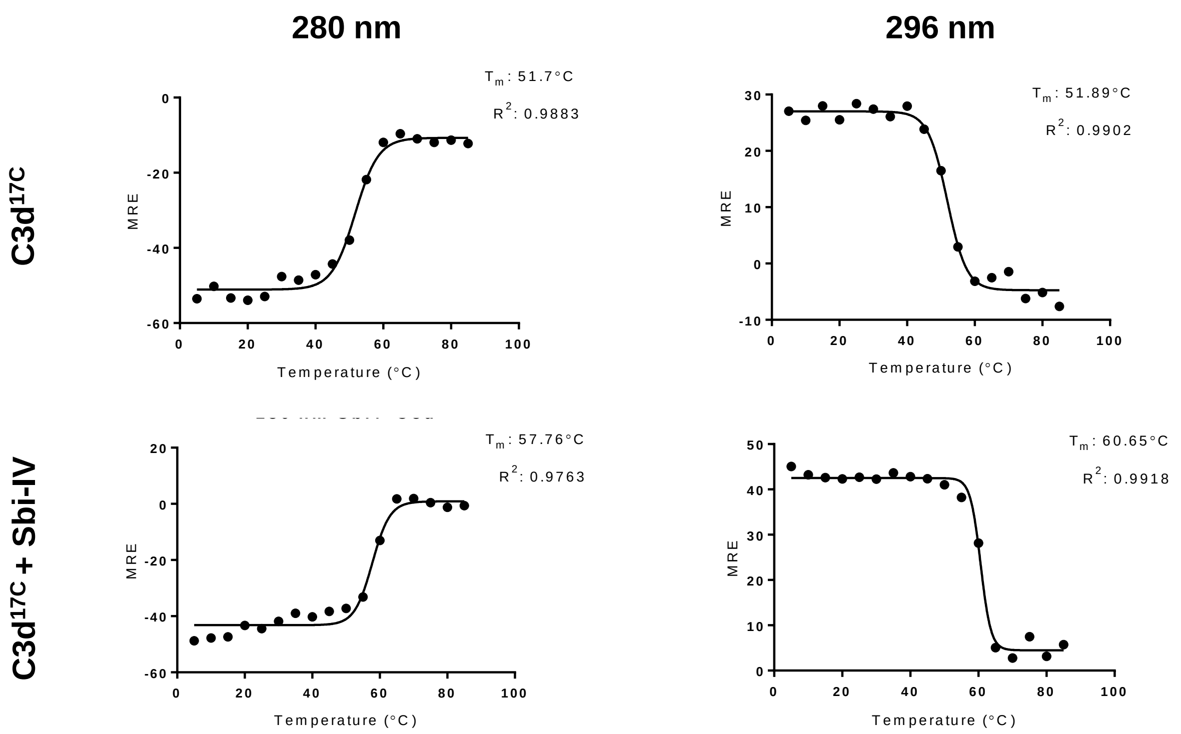


**c:** Boltzmann sigmoidal fits of thermal denaturation curves at 280 and 296 nm used for melting temperature (Tm) calculations of C3d$^{17C}$ in the absence (top) or presence (bottom) of Sbi-IV following near-UV circular dichroism spectroscopic studies performed as a function of temperature. MRE: mean residue ellipticity. See Supplementary Figure S5 for near-UV thermal melt analysis of C3d$^{17A}$. Far-UV thermal unfolding analysis of C3d$^{17C}$ and C3d$^{17A}$ secondary structure can be found in Supplementary Figure S6.

#### Supplementary Figure S5

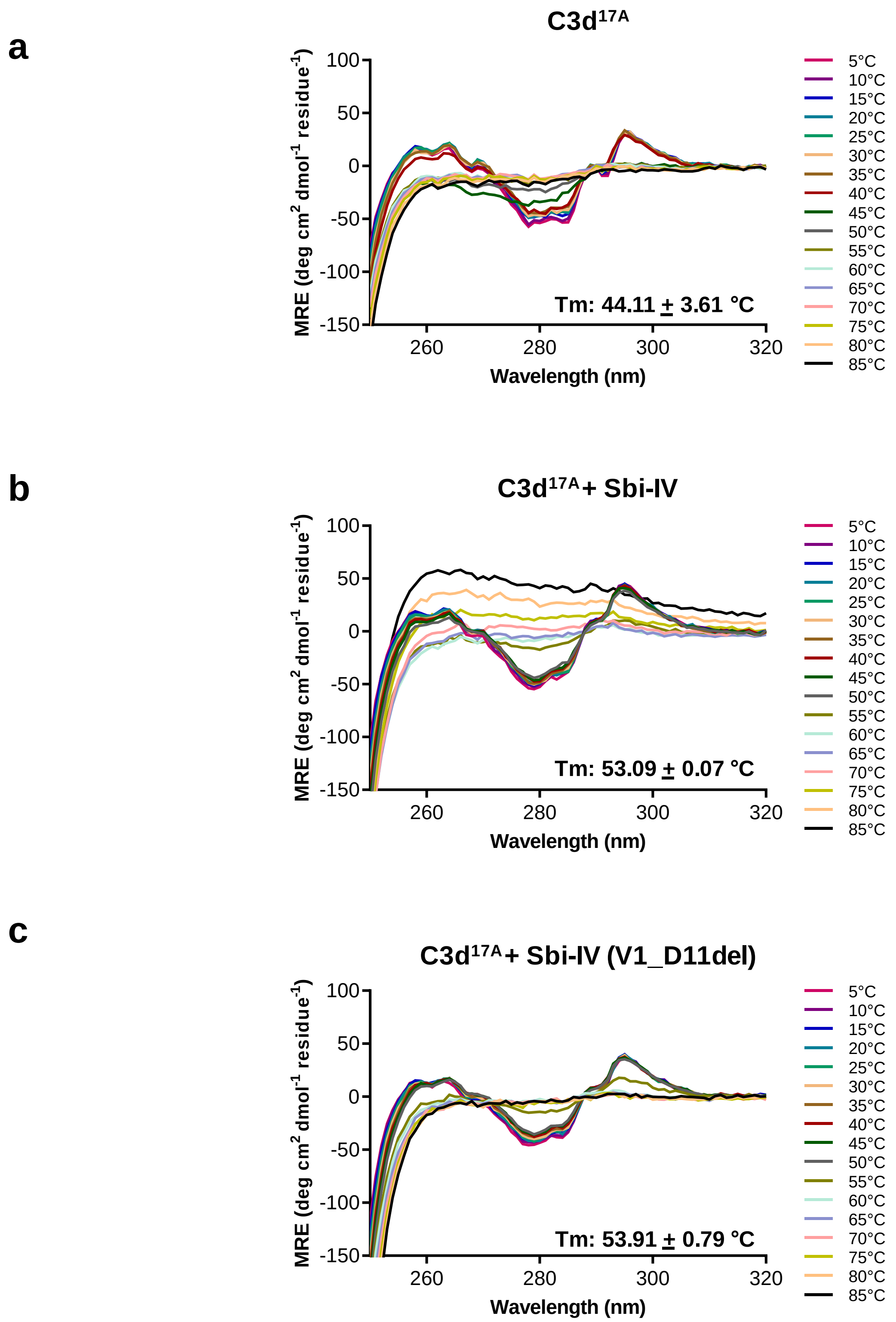


**Near-UV thermal melt circular dichroism spectroscopic studies of C3d$^{17A}$ tertiary structure in the absence or presence of wild-type Sbi-IV or a truncated Sbi-IV mutant. a, b:** In comparison to the absence of Sbi-IV (**a**), C3d$^{17A}$ in the presence of wild-type Sbi-IV (**b**), displays a ~9 °C increase in melting temperature (Tm) but also undergoes aggregation and precipitation at high temperatures. This aggregation of C3d$^{17A}$ is not evident in the absence of Sbi-IV (**a**) or in the presence of a truncated Sbi-IV mutant (V1-D11del) lacking the N-terminal region of the Sbi-IV α1 helix known to interact with helix α1 located on the convex surface of C3d (**c**).

d

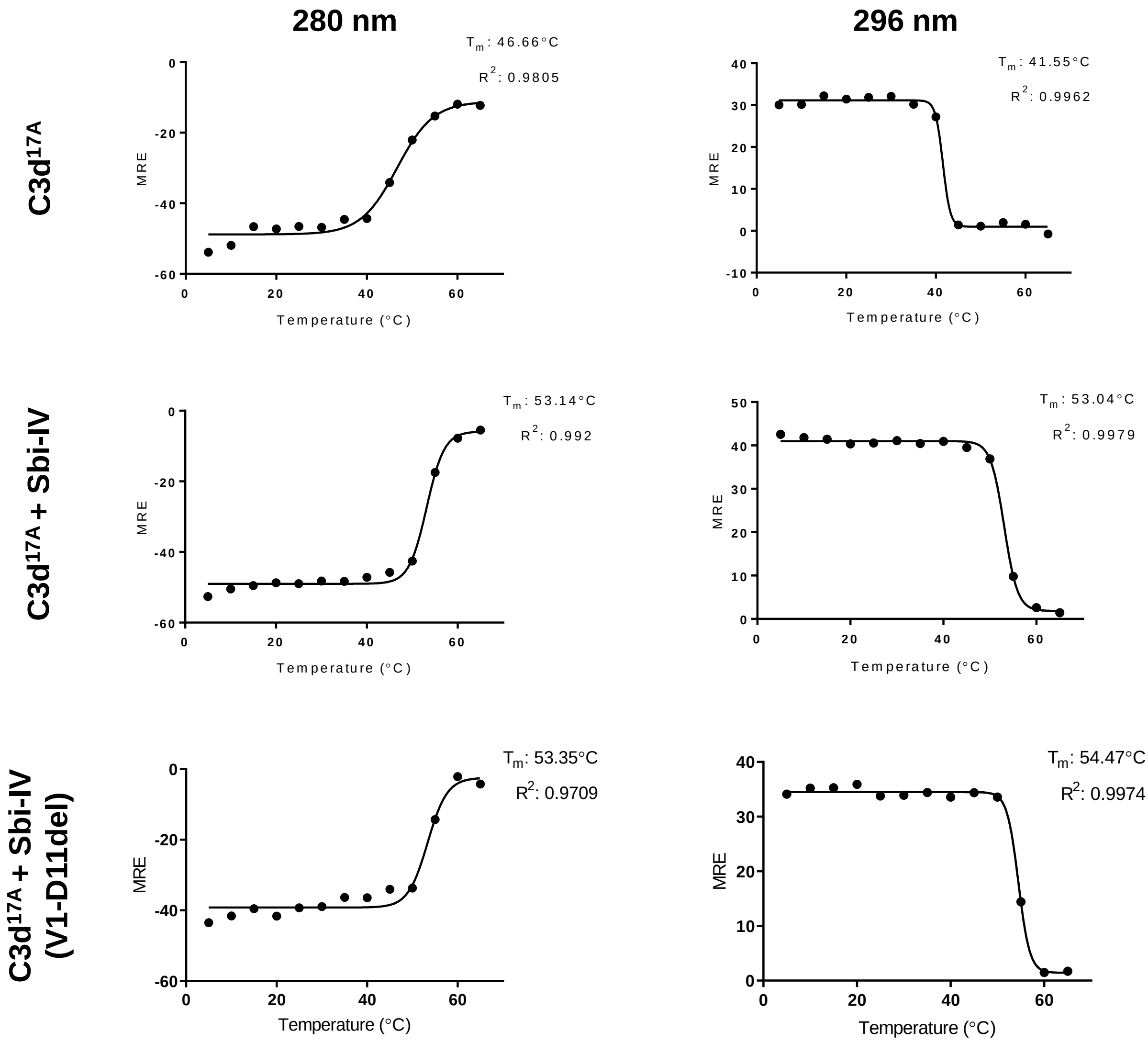


**d:** Boltzmann sigmoidal fits of thermal denaturation curves at 280 and 296 nm used for melting temperature (Tm) calculations of $C3d^{17A}$ in the absence (top) or presence of wild-type Sbi-IV (middle) or a truncated Sbi-IV mutant (V1-D11del) (bottom) following near-UV circular dichroism spectroscopic studies performed as a function of temperature. Data gathered at 65-85 °C was excluded from calculations. MRE: mean residue ellipticity.

#### Supplementary Figure S6

**a**

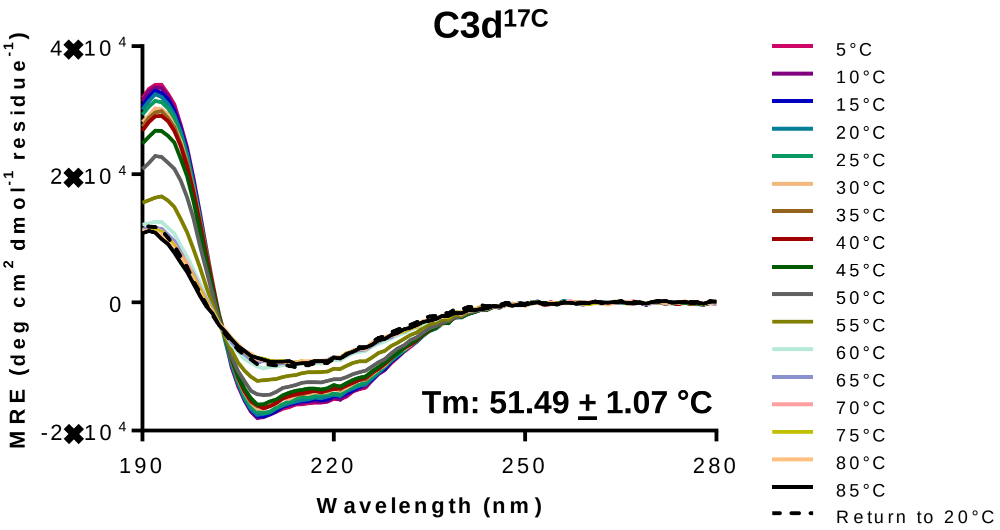


| | Reference set for CDSSTR | α helices | β sheets | Turns | Unordered | Total | NRMSD |
|---|---|---|---|---|---|---|---|
| **5°C** | 7 | 52% | 11% | 14% | 23% | 100% | 0.010 |
| **20°C** | 7 | 51% | 12% | 13% | 24% | 100% | 0.010 |
| **85°C** | 7 | 18% | 26% | 20% | 35% | 99% | 0.023 |
| **Return to 20°C** | 7 | 19% | 28% | 21% | 33% | 101% | 0.017 |

**b**

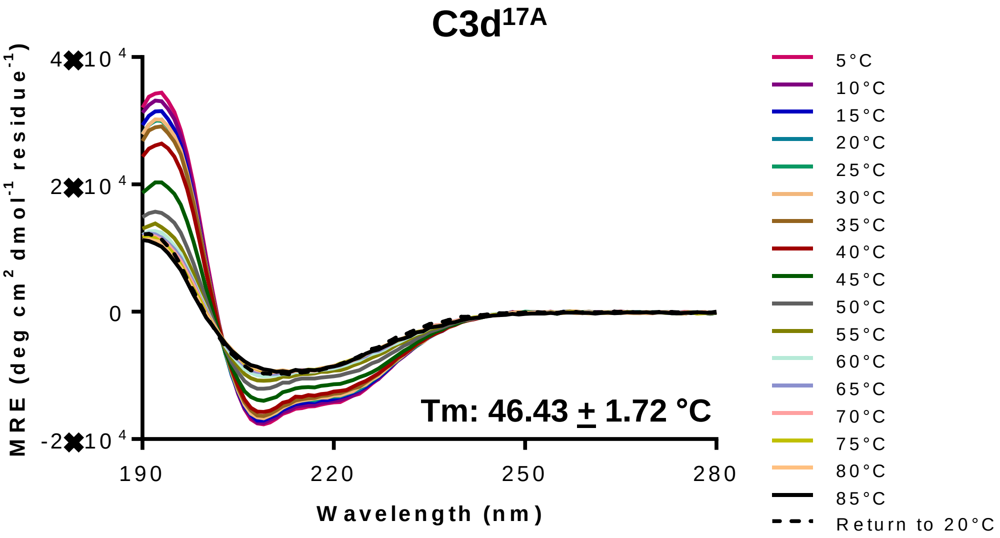


| | Reference set for CDSSTR | α helices | β sheets | Turns | Unordered | Total | NRMSD |
|---|---|---|---|---|---|---|---|
| **5°C** | 7 | 52% | 11% | 14% | 24% | 101% | 0.007 |
| **20°C** | 7 | 48% | 11% | 15% | 26% | 100% | 0.008 |
| **85°C** | 7 | 19% | 25% | 20% | 36% | 100% | 0.019 |
| **Return to 20°C** | 7 | 18% | 29% | 21% | 33% | 101% | 0.014 |

**Far-UV circular dichroism spectroscopic studies of C3d$^{17C}$** (a) **and C3d$^{17A}$** (b) **secondary structure as a function of temperature. a, b:** Both C3d$^{17C}$ (**a**) and C3d$^{17A}$ (**b**) show irreversible thermal unfolding characterised by a significant loss of α-helicity and possess melting temperatures (Tm) of 51.49 $\pm$ 1.07 °C and 46.43 + 1.72 °C respectively. Deconvolution was performed using Dichroweb.

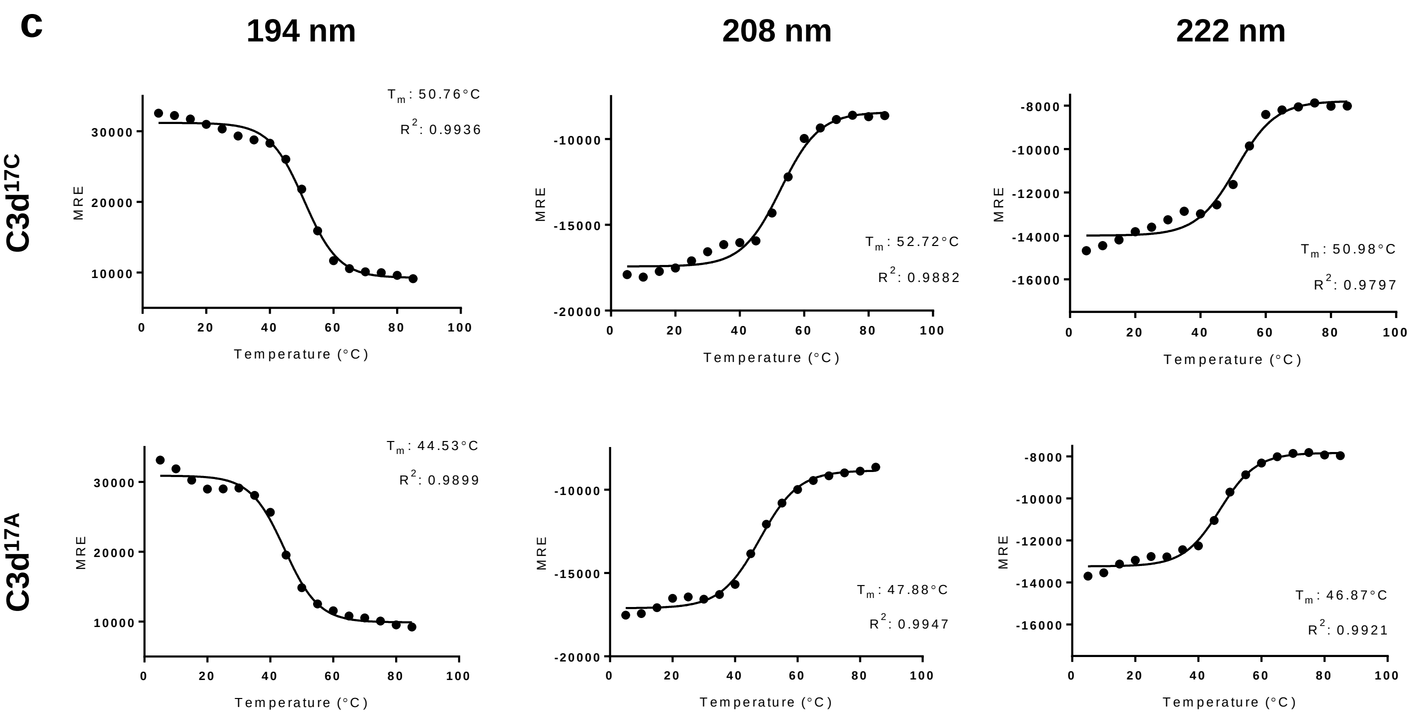


**c:** Boltzmann sigmoidal fits of thermal denaturation curves at 194, 208 and 222 nm used for Tm calculations of C3d$^{17C}$ (top) and C3d$^{17A}$ (bottom). MRE: mean residue ellipticity.

#### Supplementary Figure S7

a

b

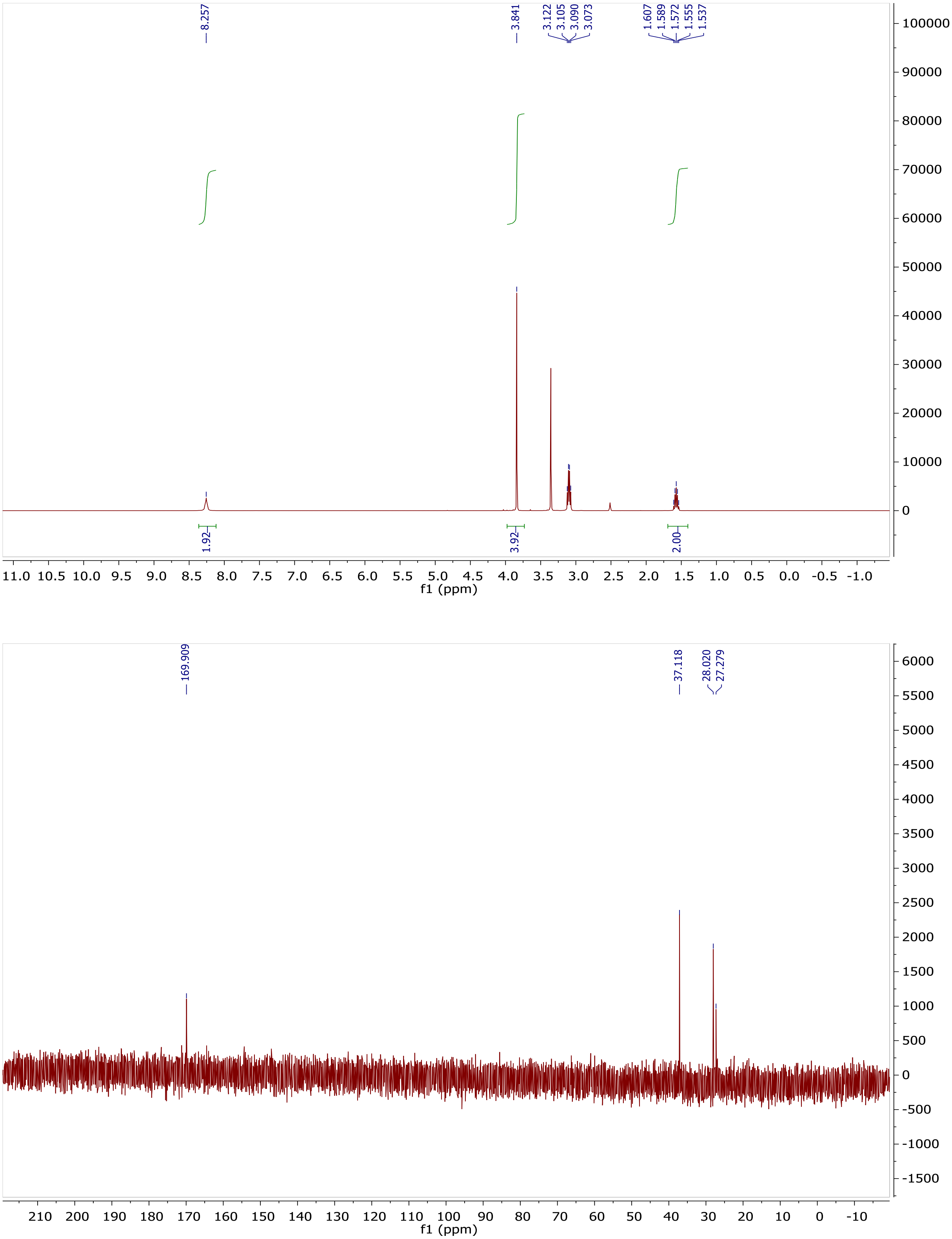


**Characterisation of *N,N′*-(propane-1,3-diyl) bis(2-bromoacetamide) linker. a:** $^{1}H$ NMR spectrum of *N,N′*-(propane-1,3-diyl) bis(2-bromoacetamide). $^{1}H$- NMR (400 MHz, d6-DMSO): δ 8.24 (s, 2H, NH), 3.83 (s, 4H, $CH_2Br$), 3.08 (m, $J$ = 6.9 Hz, 4H, $CH_2NH$), 1.56 (m, $J$ = 7.0 Hz, 2H, $CH_2CH_2CH_2$). **b**: $^{13}C$ NMR spectrum of *N,N′*-(propane-1,3-diyl) bis(2-bromoacetamide). $^{13}C$-NMR (400 MHz, $D_2O$): δ 169.91, 37.12, 28.02, 27.28.

#### Supplementary Figure S8

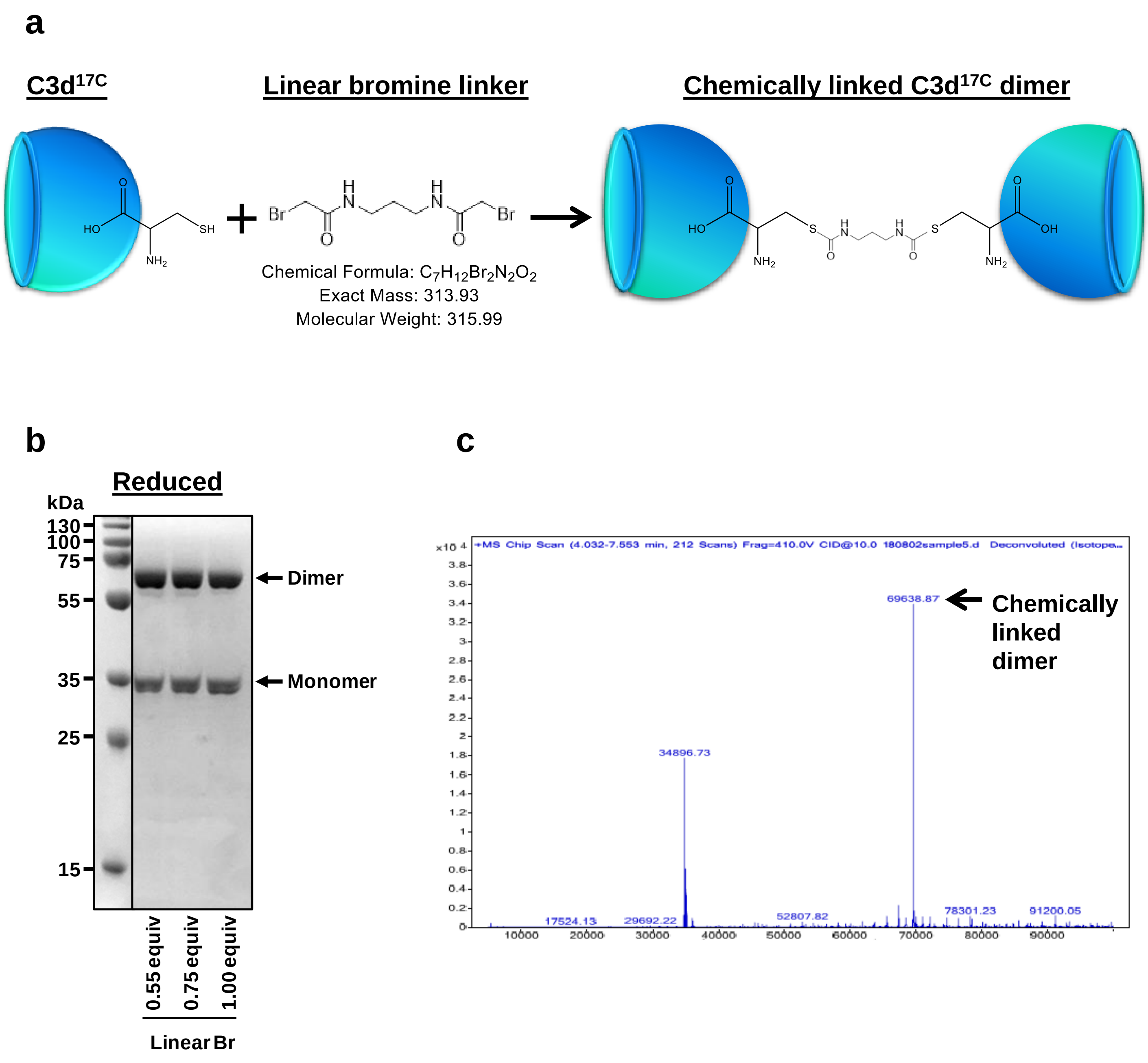


**Induced dimerisation of C3d$^{17C}$ using chemical linkage. a:** Schematic depicting formation of C3d$^{17C}$ dimers using chemical linkage with a linear bromine linker (*N,N'*-(propane-1,3-diyl) bis(2-bromoacetamide). When in close proximity to the sulfhydryl group of free cysteines (position 17 of C3d), the linker will link together two C3d$^{17C}$ monomers resulting in the formation of durable chemically-linked C3d$^{17C}$ dimers. During the reaction both terminal bromines from the linker and hydrogen atoms from the sulfhydryl groups of C3d free cysteines are lost. **b**: Reducing SDS-PAGE depicting the formation of an appreciable amount of chemically-linked C3d$^{17C}$ dimers resistant to reduction following conjugation with the linear bromine linker at different molar equivalences. **c**: Mass spectrometric analysis confirming the presence of a dimeric C3d species of a mass (69,638.87 Da) consistent with 2 C3d$^{17C}$ molecules (deficient in one proton each from the SH group) and the linear linker lacking both terminal bromines. The smaller peak at 34,896.73 Da indicates a monomeric C3d$^{17C}$ species that has also reacted with the linker but failed to form a dimer.

#### Supplementary Figure S9

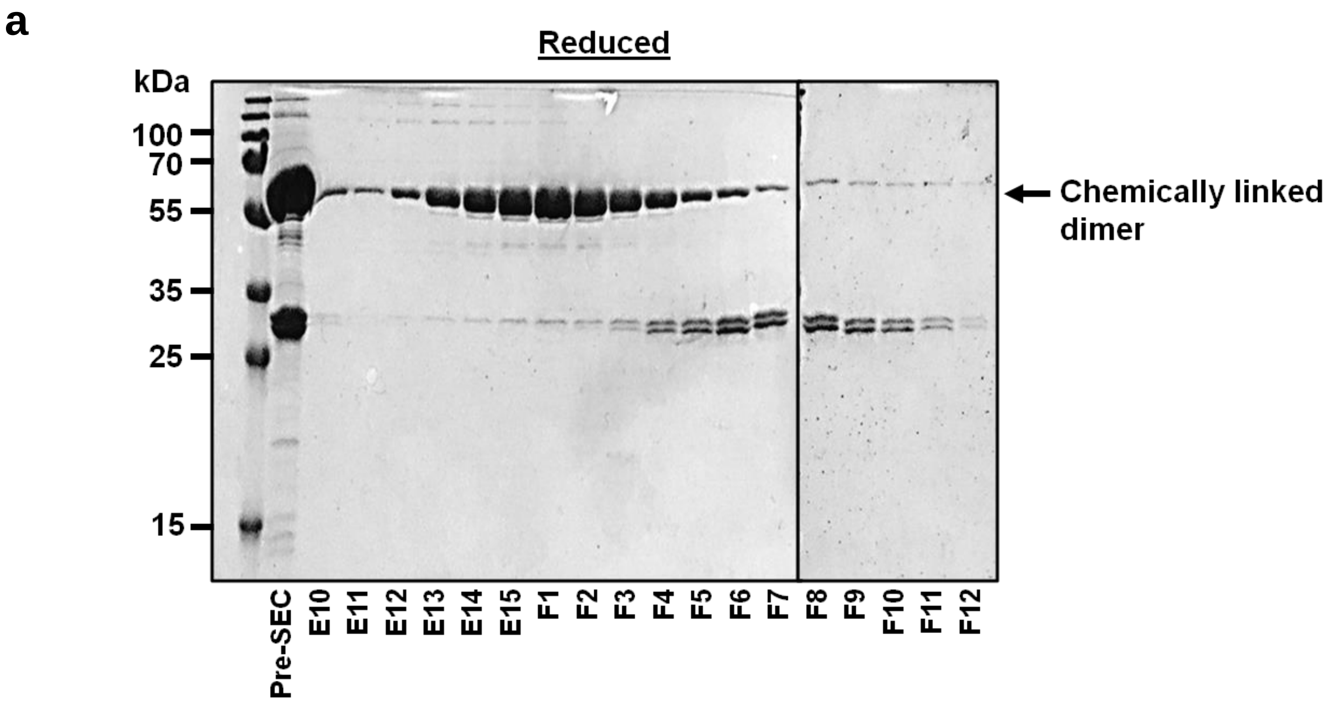


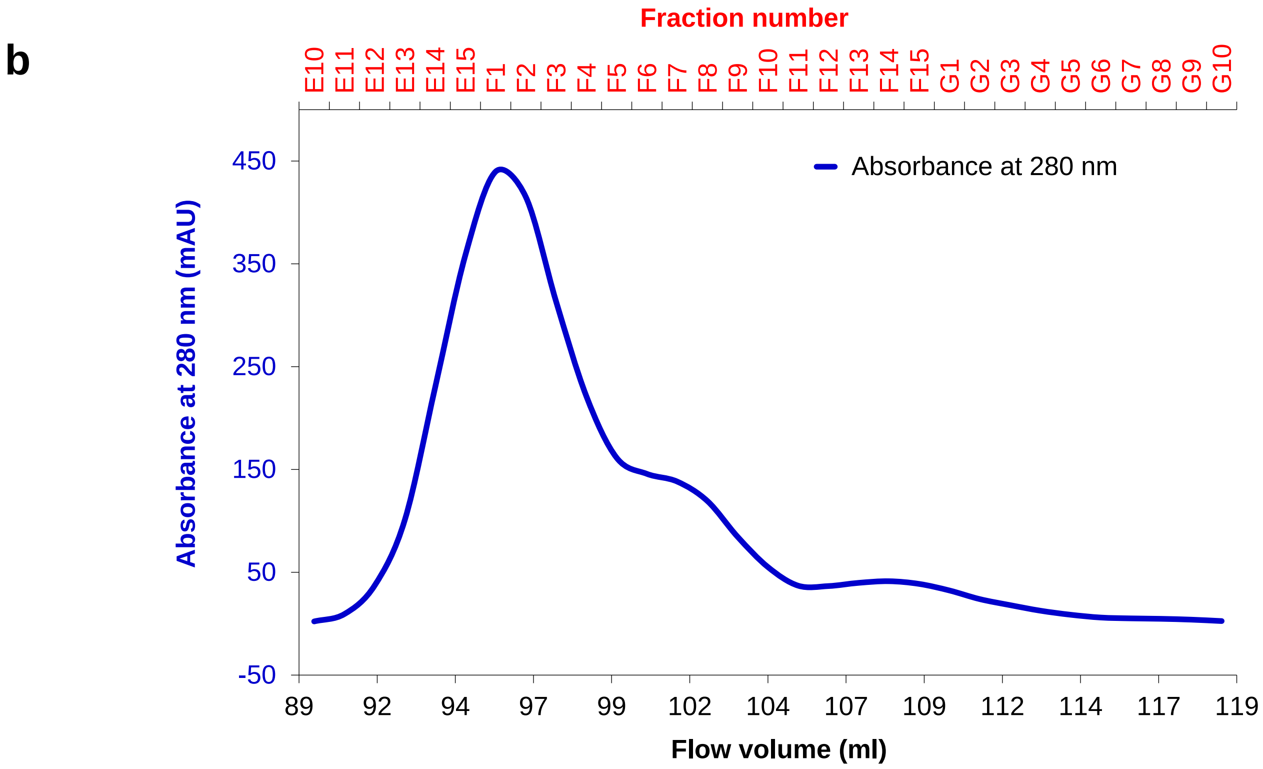


**Purification and characterisation of chemically-linked C3d$^{17C}$ dimers. a:** Reducing SDS-PAGE of elution fractions from size exclusion chromatography of chemically-linked dimeric C3d$^{17C}$. **b**: Chromatogram showing elution peak of chemically-linked dimeric C3d$^{17C}$ size exclusion chromatography. Pre-SEC: pre-size exclusion chromatography.

**c**

## 3.75 µM

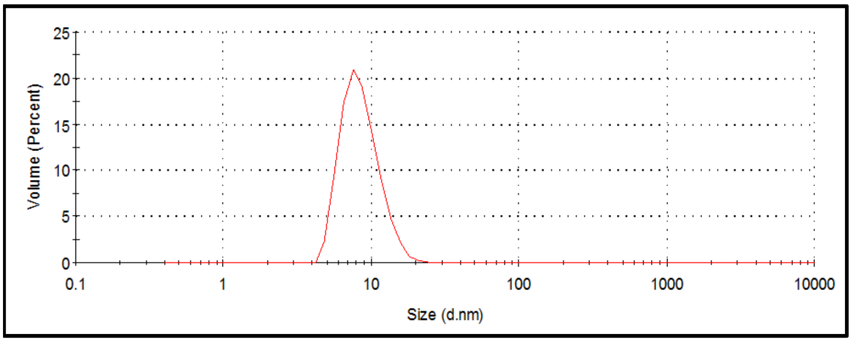


| Size (d.nm): | Volume (%): | St Dev (d.nm): | PdI |
|---|---|---|---|
| 8.623 | 99.9 | 2.529 | 0.172 |

## 7.5 µM

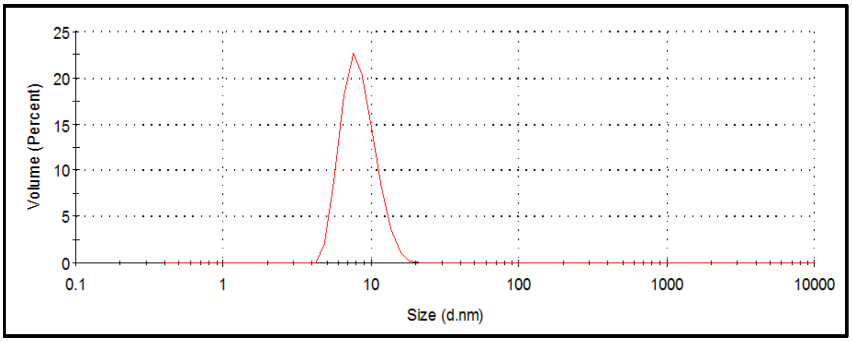


| Size (d.nm): | Volume (%): | St Dev (d.nm): | PdI |
|---|---|---|---|
| 8.421 | 99.8 | 2.209 | 0.30 |

## 15 µM

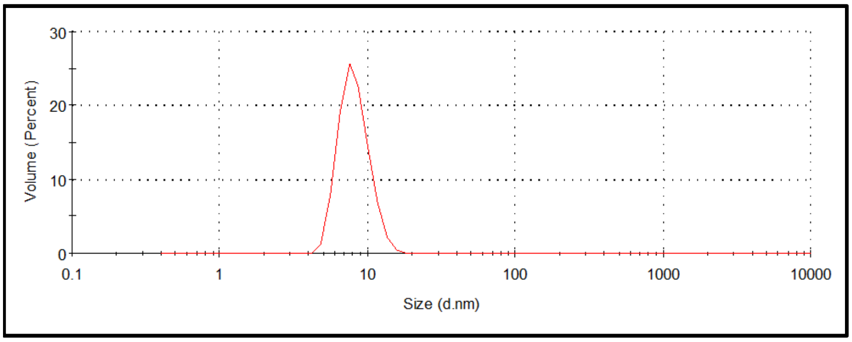


| Size (d.nm): | Volume (%): | St Dev (d.nm): | PdI |
|---|---|---|---|
| 8.223 | 100.0 | 1.873 | 0.204 |

**c:** Particle size analysis of chemically-linked dimeric C3d$^{17C}$. Dynamic light scattering size distribution by volume plots of chemically-linked dimeric C3d$^{17C}$ at concentrations of 3.75 µM (top), 7.5 µM (middle) and 15 µM (bottom) showing a single species of 8.422 $\pm$ 0.2 nm and lack of larger complexes or aggregates. Plots are representative of ≥11 measurements. PdI: polydispersity index.

**d**

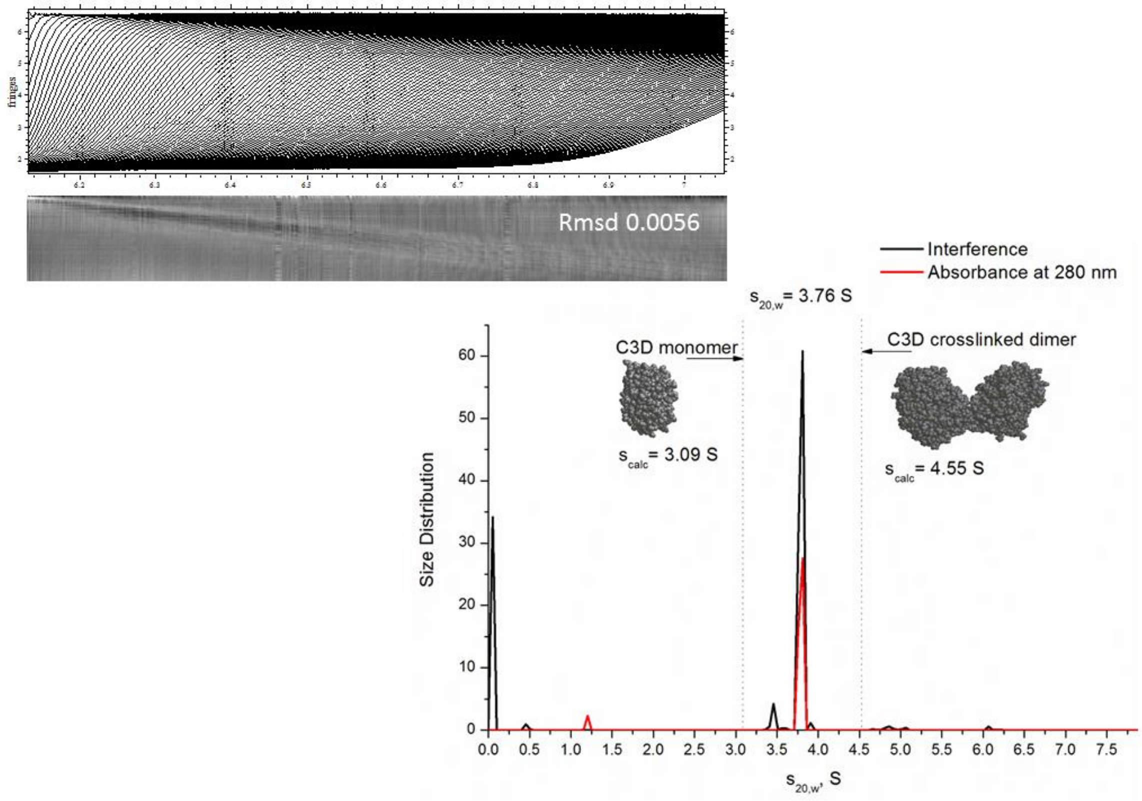


**d:** Analytical ultracentrifugation of chemically-linked C3d$^{17C}$. Comparison of the observed sedimentation coefficient with hydrodynamic calculations of the C3d$^{17C}$ dimer produced using SoMo and HYDROPRO confirms the main species is dimeric C3d$^{17C}$.

**e**

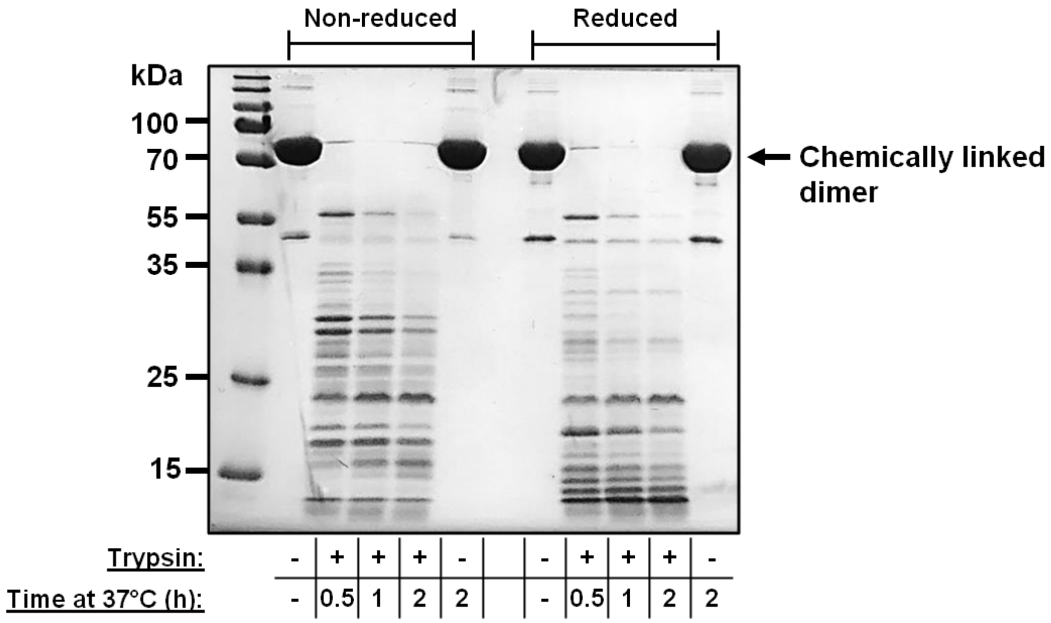


**e:** Trypsin digestion of chemically-linked dimeric C3d$^{17C}$ visualised using non-reducing and reducing SDS-PAGE. See Supplementary Tables S3 and S4 for mass spectrometric analysis of the digested fragments.

### Supplementary Figure S10

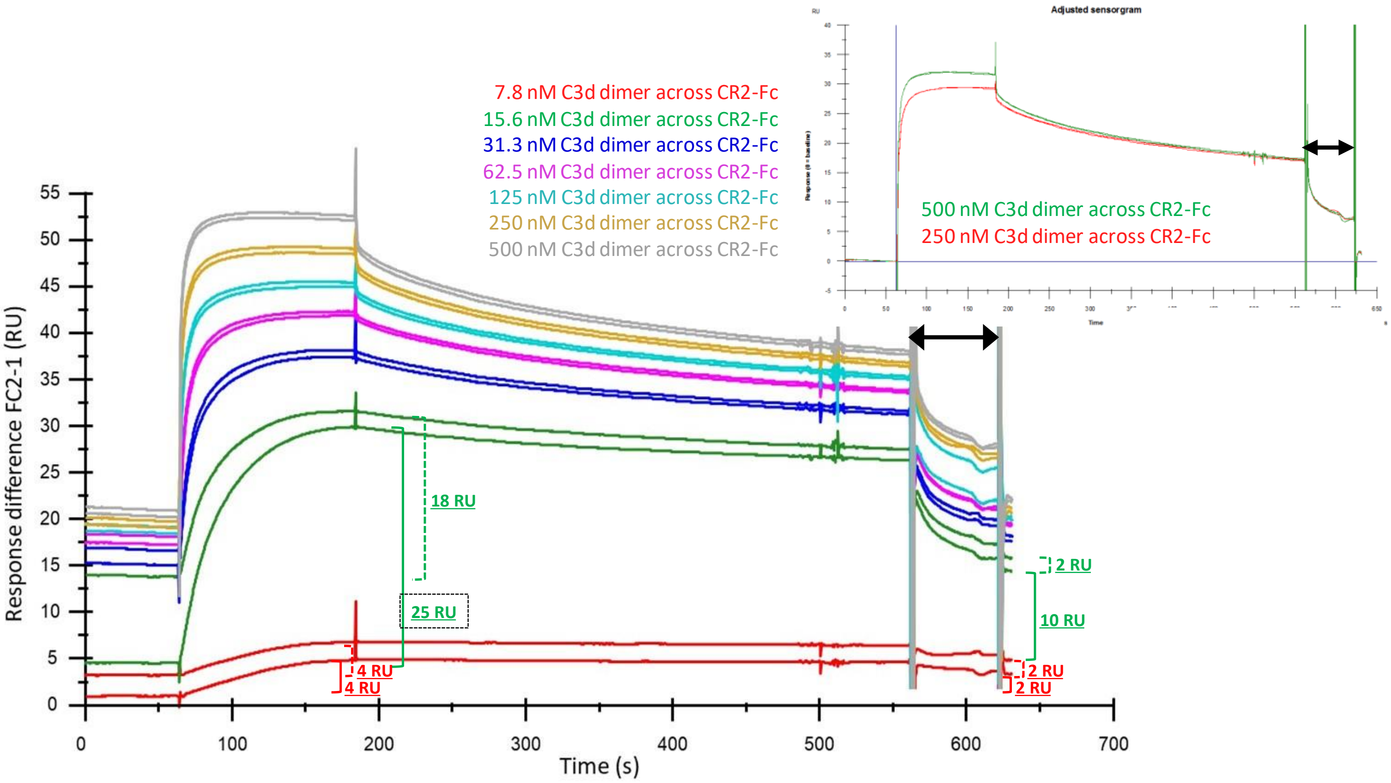


**Analysis of C3d$^{17C}$ dimer-CR2-Fc sensorgram.** Displayed is a sensorgram included in Figure 4a showing serially-diluted concentrations of 500 nM C3d$^{17C}$ dimer flowed in duplicate over a flow cell immobilised with CR2-Fc (SCR1-4). When 7.8 nM C3d$^{17C}$ dimer is flowed across the surface, 4 RU are bound at the end of the injection and 2 RU remain tightly bound after the regeneration, thus the new baseline is 2 RU higher for the duplicate sample. When the first injection of 15.6 nM is flowed across the surface, 25 RU bind at Req, 10 RU cannot be removed while 15 RU are readily eluted from the surface. When the second injection of 15.6 nM is flowed across the surface, 18 RU binds at Req, 16 RU are readily eluted and 2 RU remain tightly bound. It is clear that the majority of the high affinity sites are saturated during the first injection at 15.6 nM as the binding interactions observed during the duplicate injection are readily disrupted. In all subsequent cycles, from 31.3 nM, the majority of the binding interactions are readily disrupted with only 1 or 2 RU remaining bound after the cycle. At the highest concentrations (500 and 250 nM), all binding interactions are readily disrupted (baseline-adjusted sensorgram inset) and there is evidence of a minor very weak (fast on-off) interaction. Arrows depict the regeneration period.

### Supplementary Figure S11

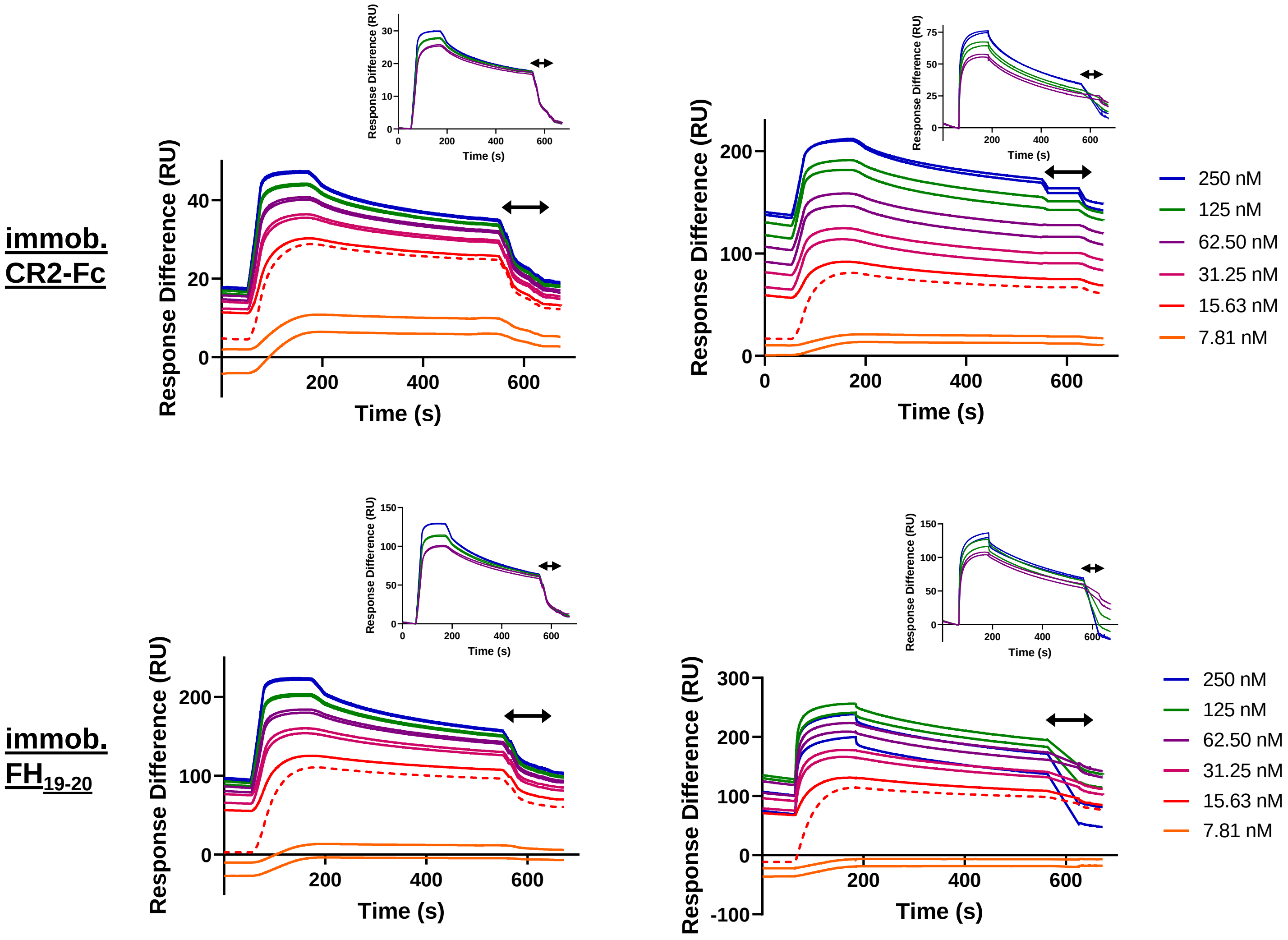


**SPR sensorgrams of C3d$^{17C}$ dimer binding interactions with immobilised CR2-Fc and FH$_{19\text{-}20}$.** Sensorgrams from two independent experiments showing serially-diluted concentrations of 250 nM C3d$^{17C}$ dimer flowed in duplicate over flow cells of a CM5 senor chip immobilised with CR2-Fc (SCR1-4) (top) or FH$_{19\text{-}20}$ (bottom). Similar to the pattern observed in Figure 4a, the curves suggest a two-state binding interaction with the formation of high avidity crosslinked complexes that fail to regenerate fully at low concentrations and less favourable, possible 1:1 interactions at higher concentrations. Baseline-adjusted sensorgrams displayed inset show the less favourable interactions at high concentrations that are readily eluted from the surface. Arrows depict the regeneration period.

### Supplementary Figure S12

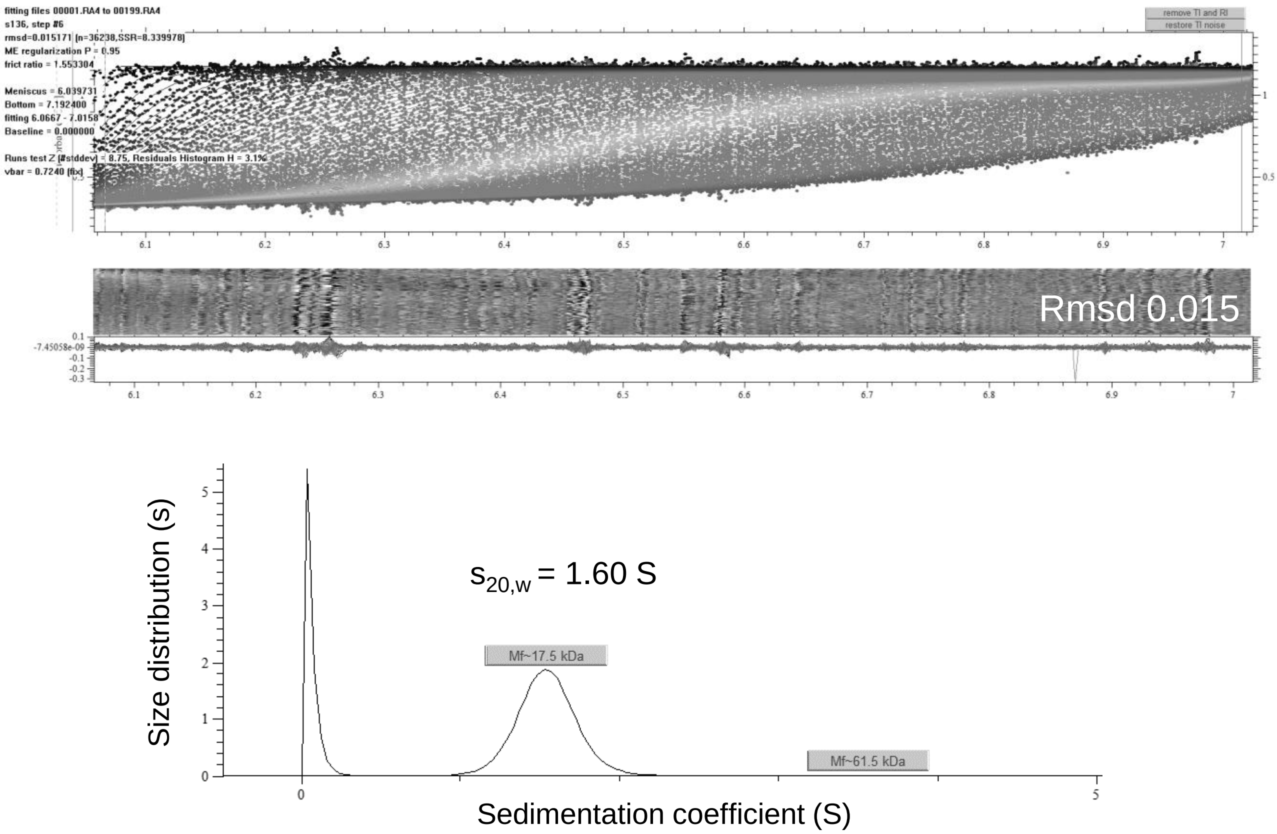


**Analytical ultracentrifugation analysis of $FH_{19\text{-}20}$.** The observed sedimentation coefficient of 1.60 S confirms the monomeric state of $FH_{19\text{-}20}$ (~17.5 kDa).

### Supplementary Figure S13

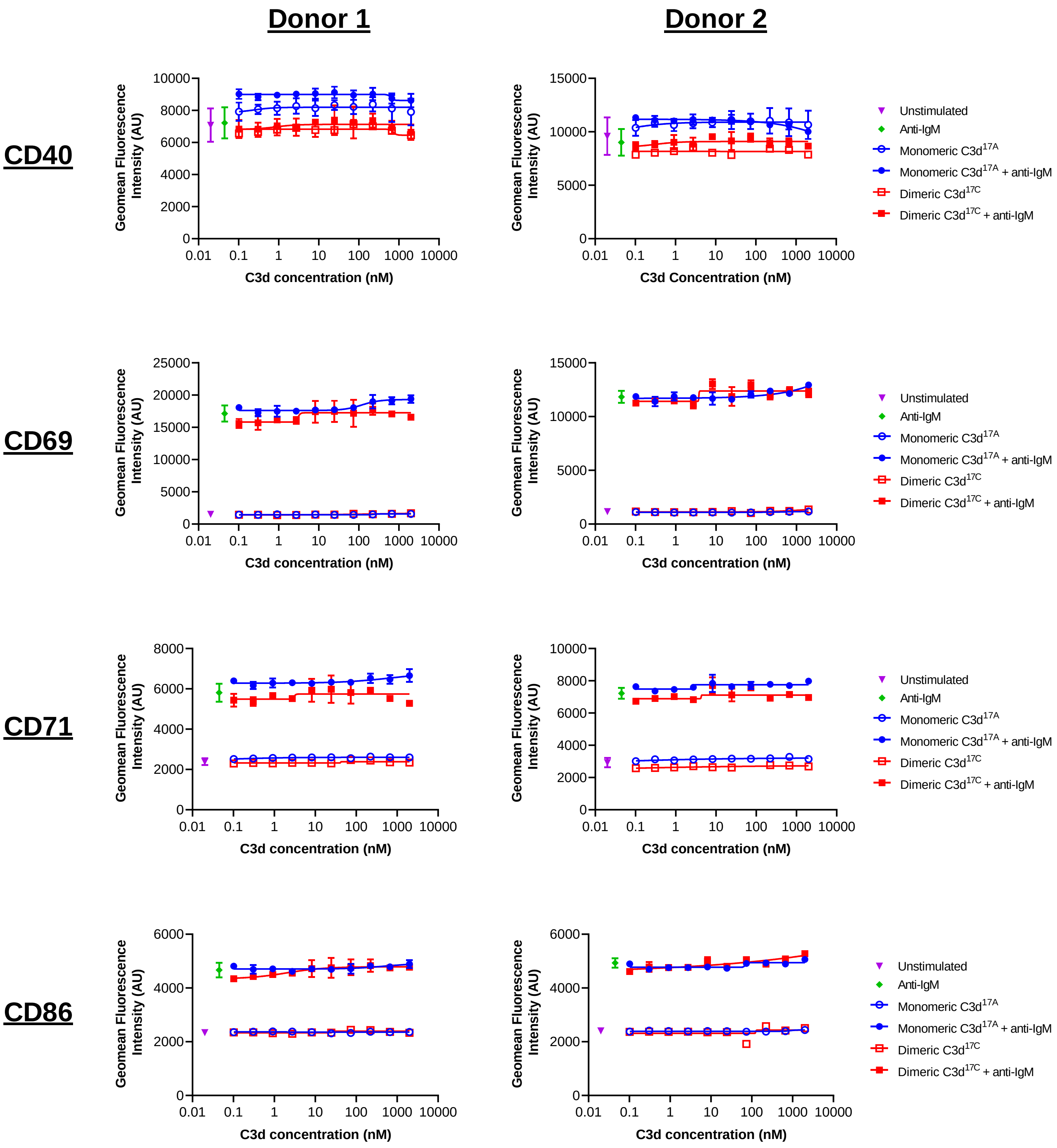


**Flow cytometric analysis of C3d-mediated changes in the activation state of purified B cells.** In contrast to experiments probing the activation of B cell populations in PBMCs (Figure 5, Supplementary Figure S17), incubation of purified B cells with monomeric C3d$^{17A}$ or dimeric C3d$^{17C}$ failed to induce any significant changes in expression of the four surface-associated activation markers examined. Data are of B cells purified from two representative donors and displayed as mean values (n=2) ± standard deviation from the mean with curves fitted using a non-linear regression model.

**Supplementary Figure S14**

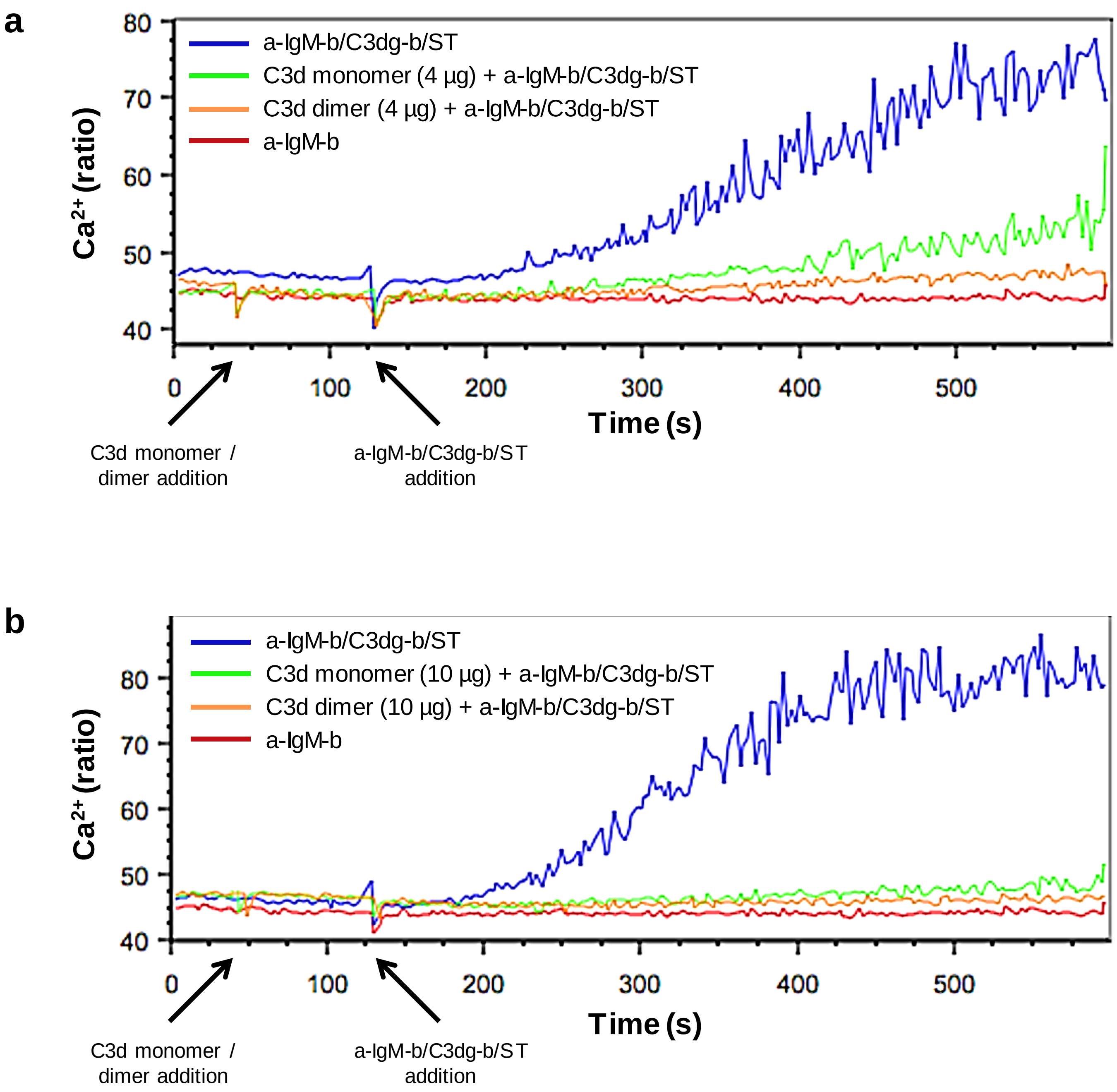


**Preligation of CR2 with monomeric C3d$^{17A}$ or dimeric C3d$^{17C}$ inhibits BCR/CR2-dependent Ca$^{2+}$ influx in murine B cells.** Incubation with 4 µg C3d$^{17A}$ monomer or C3d$^{17C}$ dimer (30 s) 90 seconds prior to the addition of BCR/CR2-crosslinking complexes (a-IgM-b/C3dg-b/ST) (120 s) significantly retards and reduces Ca$^{2+}$ influx in CD45R/B220-gated Indo 1-AM-loaded C57BL/6 mouse splenocytes with a more pronounced blocking effect apparent with dimeric C3d$^{17C}$ (**a**). 10 µg of monomeric C3d$^{17A}$ or dimeric C3d$^{17C}$ completely eliminates BCR/CR2-dependent Ca$^{2+}$ influx (**b**) suggesting the observed blocking effect is concentration dependent and likely a result of CR2 sequestration by monomeric C3d$^{17A}$/dimeric C3d$^{17C}$ reducing the proportion of CR2 available for crosslinking with BCR. BCR/CR2-crosslinking complexes were composed of a suboptimal dose (0.056 µg mL$^{-1}$) of biotinylated F(ab')$_2$ goat anti-mouse IgM (a-IgM-b), C3dg-biotin (C3dg-b) and streptavidin (ST). The C3d$^{17A}$ monomer/C3d$^{17C}$ dimer-mediated blocking of Ca$^{2+}$ influx was not evident when higher, more optimal concentrations of a-IgM-b/ST were used or when all the reaction components were added simultaneously.

#### Supplementary Figure S15

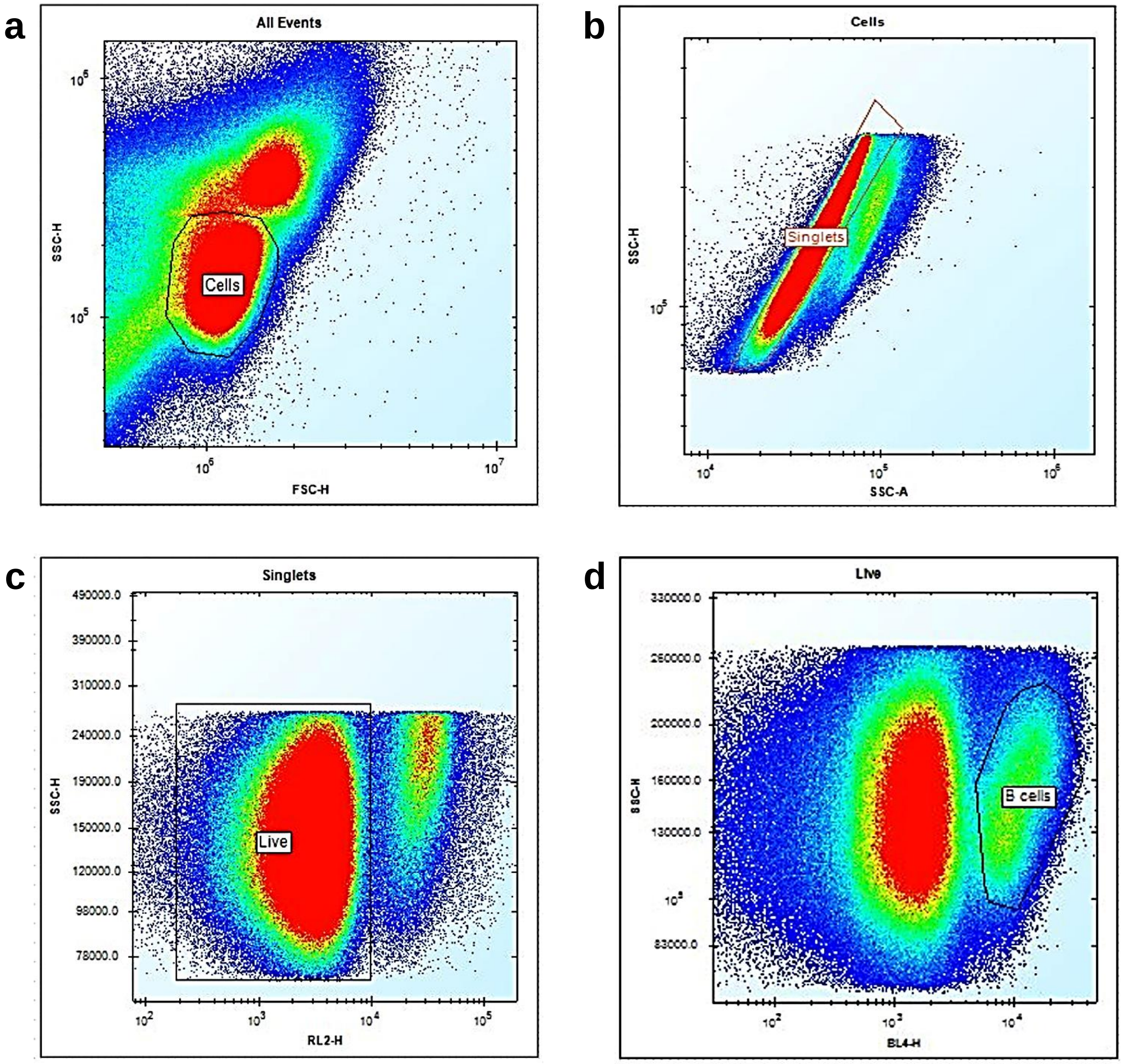


**Flow cytometry gating strategy for B cells in PBMC samples. a:** Identification of cells was performed on the basis of forward (FSC) and side scatter (SSC) profile. **b**: Singlets were gated from a plot showing the area (A) against the height (H) of the side scatter peak of cells. **c**: A live/dead gate was applied to sort for live singlet cells which tested negative for a fixable near-infrared dead cell stain (red laser 2 (RL2)). **d**: A Cy5-conjugated anti-human CD19 antibody (blue laser 4 (BL4)) was used on the live singlet cells to derive CD19$^{+}$ B cells.

#### Supplementary Figure S16

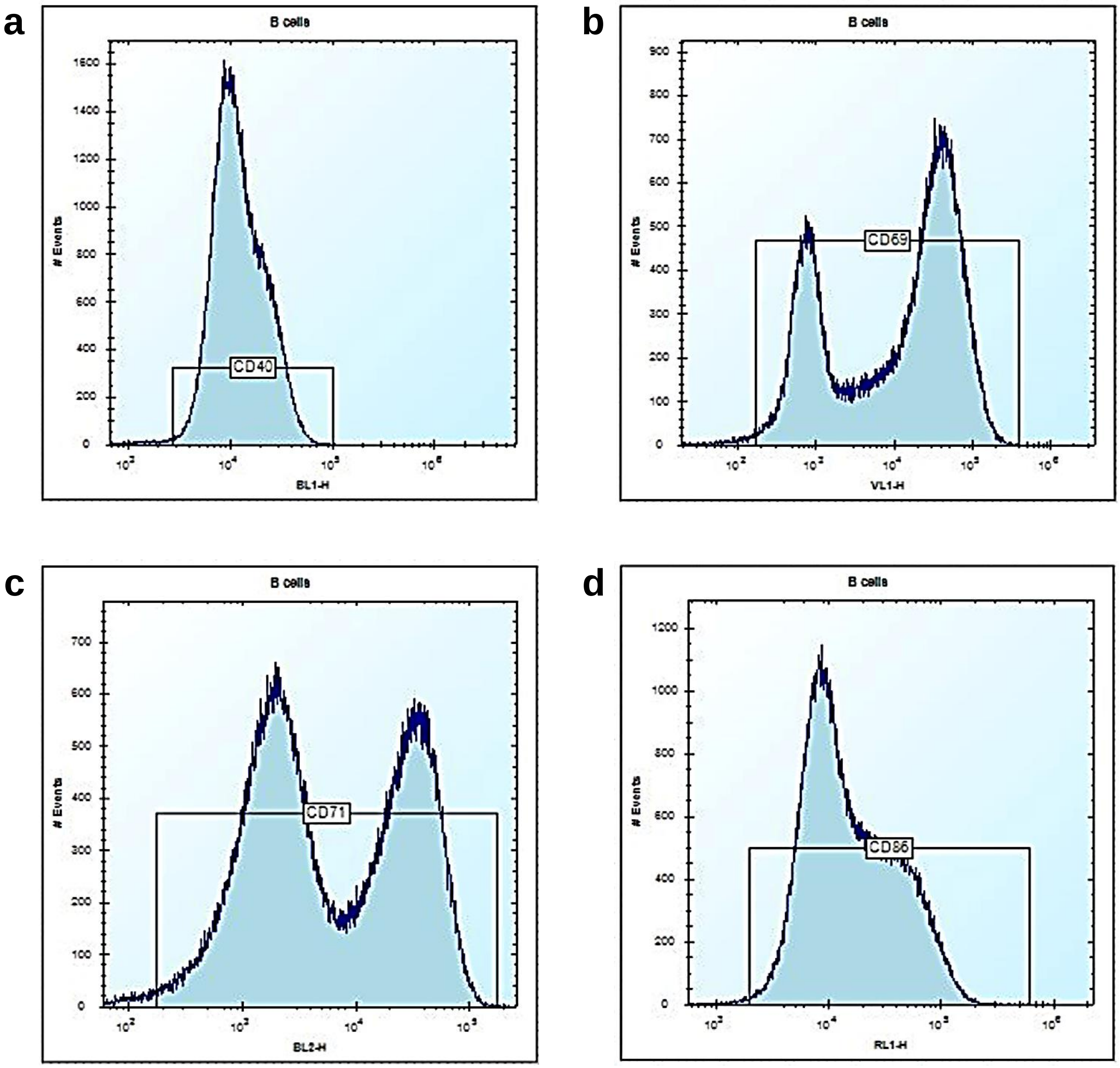


**Flow cytometry channel gating of B cell activation markers**. Representative histograms depicting gating applied to $CD19^{+}$ B cell populations for the following activation markers are shown: (**a**) CD40 (FITC anti-CD40: blue laser 1 (BL1)), (**b**) CD69 (BV421 anti-CD69: violet laser 1 (VL1)), (**c**) CD71 (PE anti-CD71: blue laser 2 (BL2)) and (**d**) CD86 (APC anti-CD86: red laser 1 (RL1)).

### Supplementary Figure S17

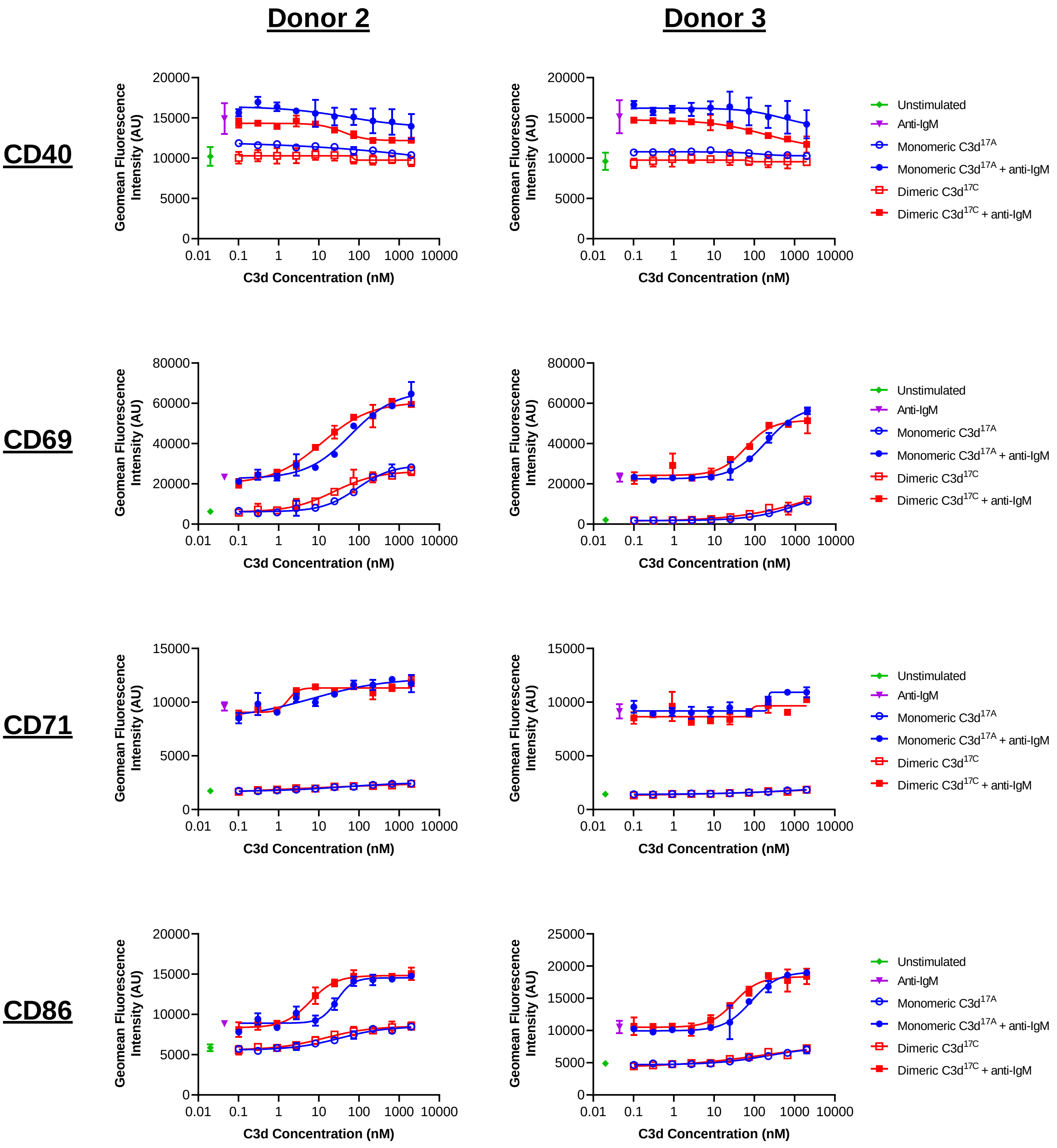


**Flow cytometric analysis of C3d-mediated changes in the activation state of PBMC B cell populations from an additional two donors.** Consistent with the data generated for donor 1 (Figure 5), the B cell populations from donors 2 and 3 display similar C3d dose-dependent trends in expression of the activation markers examined. Data are displayed as mean values (n=2) ± standard deviation from the mean with curves fitted using a non-linear regression model.

#### Supplementary Table S1

**Supplementary Table S1. Data collection statistics.**

| | C3d$^{17C}$ dimer | C3d$^{17C}$ dimer - Sbi-IV complex |
|---|---|---|
| **Space group** | P $2_1$ $2_1$ $2_1$ | P $2_1$ $2_1$ $2_1$ |
| **Unit cell dimensions** | | |
| **α, b, c (Å)** | 54.579, 59.914, 175.460 | 73.609, 115.159, 118.797 |
| **α, β, γ (˚)** | 90.0, 90.0, 90.0 | 90.0, 90.0, 90.0 |
| **Molecules/asymmetric unit** | 2 | 4 |
| **Wavelength (Å)** | 0.9795 | 0.9795 |
| **Resolution limits (Å)** | 58.49 - 1.87 (1.90 - 1.87) | 115.16 - 2.27 (2.31 - 2.27) |
| **Total reflections** | 1232513 (54652) | 597404 (27526) |
| **Unique reflections** | 48649 (2307) | 47004 (2149) |
| $R_{merge}$ | 0.424 (0.819) | 0.439 (5.161) |
| ***I/σI*** | 11.2 (3.4) | 7.9 (1.6) |
| **Completeness (%)** | 99.99 (96.77) | 99.75 (92.19) |
| **Multiplicity** | 25.3 (23.7) | 12.7 (12.8) |

Values in parentheses are for high resolution shell

### Supplementary Table S2

**Supplementary Table S2. Refinement statistics.**

| | C3d$^{17C}$ dimer | C3d$^{17C}$ dimer - Sbi-IV complex |
|---|---|---|
| **Resolution (Å)** | 2.0 | 2.4 |
| $R_{work}$ | 0.1422 | 0.1560 |
| $R_{free}$ | 0.1996 | 0.2107 |
| **Atoms (no.)** | 10263 | 11854 |
| **Residues (no.)** | 599 | 727 |
| **Ordered water molecules (no.)** | 534 | 260 |
| **Ligands / ions (no.)** | 4 | 2 |
| **R.m.s.d. bonds (Å)** | 0.008 | 0.012 |
| **R.m.s.d. angles (°)** | 0.948 | 1.102 |
| **Average *B*-factors (Å$^2$)** | 27.92 | 30.82 |

### Supplementary Table S3

#### Supplementary Table S3a. Mass spectrometry species of trypsin-digested dimeric C3d$^{17C}$ confirming chemical linkage of C3d$^{17C}$ at position 17C.

| Predicted formula of peptide | RT | Precursor | Mass | Score | Diff (ppm) | Ion Polarity | Ions | Height |
|---|---|---|---|---|---|---|---|---|
| $C_{356}H_{563}N_{91}O_{114}S_7$ | 6.878 | 1362.16 | 8160.92 | 77.86 | 1.27 | Positive | 79 | 210695 |

#### Supplementary Table S3b. Confirmatory ions detected for $C_{356}H_{563}N_{91}O_{114}S_7$

| m/z | Species | Abund |
|---|---|---|
| 1361.314 | (M+6H)+6 | 3296.6 |
| 1655.403 | (M+5Na)+5 | 3203.7 |
| 1383.163 | (M+6Na)+6 | 2780.1 |
| 1167.277 | (M+7H)+7 | 8300.9 |
| 1021.492 | (M+8H)+8 | 8418.2 |
| 1633.786 | (M+5H)+5 | 1413.8 |
| 1188.849 | (M+7Na)+7 | 1851.5 |

Chemically-linked dimeric C3d$^{17C}$ was digested with trypsin for 0.5 hours at 37°C (Supplementary Figure S9e) and subjected to electrospray ionization time-of-flight mass spectrometry. The predicted formula $C_{356}H_{563}N_{91}O_{114}S_7$ of a peptide containing two C3d trypsin-digested fragments linked by the linear bromine linker at position 17C was calculated using the two most abundant trypsin-digested peptide fragments surrounding position 17C of C3d (MLDAERLKHLIVTPSGCGEQNMIGMTPTVIAVHYLDETE: $C_{186}H_{303}N_{49}O_{60}S_4$, CGEQNMIGMTPTVIAVHYLDETEQWEKFGLEK: $C_{163}H_{250}N_{40}O_{52}S_3$) and the formula of the linker with bromines removed ($C_7H_{12}N_2O_2$) and loss of 2 H from the C3d fragments. A search for species consistent with this formula using the MassHunter Qualitative Analysis software (Agilent) yielded a species with multiple charge states providing validation that linkage of the bromine linker had occurred at position 17C of C3d. RT, retention time; Diff (ppm), difference (parts per million); m/z, mass-to-charge ratio; Abund, abundance.

#### Supplementary Table S4

**Supplementary Table S4. Trypsin digest mass spectrometry data confirming presence of an intact internal disulphide bond in chemically-linked dimeric C3d$^{17C}$.**

| Sequence of internal disulphide-containing intra-chain fragment (aa no.) | m/z | Mass | Height | Area | Missed cleavages | Matched MS/MS ions | FDR | Mod. |
|---|---|---|---|---|---|---|---|---|
| NLIAIDSQVLCGAVKWLILEKQKPDGVFQEDAPVIHQEMIGGLR (98-141) + KDMALTAFVLISLQEAKDICEEQVNSLPGSITK (146-178) | 1060.4418 | 8475.4652 | 391536 | 1818866 | 2+2 | 51 | 0 | Oxidn. |
| APSTWLTAYVVKVFSLAVNLIAIDSQVLCGAVKWLILEK (80-118) + NNNEKDMALTAFVLISLQEAKDICEEQVNSLPGSITK (142-178) | 1189.3475 | 8318.3645 | 289286 | 1214157 | 2+2 | 98 | 0 | |
| RAPSTWLTAYVVKVFSLAVNLIAIDSQVLCGAVK (79-112) + DMALTAFVLISLQEAKDICEEQVNSLPGSITKAGDFLE (147-184) | 966.6524 | 7725.1066 | 78310 | 306911 | 2+2 | 57 | 0.52 | |
| APSTWLTAYVVKVFSLAVNLIAIDSQVLCGAVKWLILE (80-117) + KDMALTAFVLISLQEAKDICEEQVNSLPGSITK (146-178) | 1103.8713 | 7719.0877 | 44584 | 214446 | 2+2 | 111 | 0 | |
| IAIDSQVLCGAVKWLILEKQKPDGVFQEDAPVIHQEMIGGLR (100-141) + ALTAFVLISLQEAKDICEEQVNSLPGSITK (149-178) | 1126.0388 | 7874.1921 | 36848 | 213979 | 2+1 | 80 | 0 | Oxidn. |
| AIDSQVLCGAVKWLILEKQKPDGVFQEDAPVIHQEMIGGLR (101-141) +DMALTAFVLISLQEAKDICEEQVNSLPGSITKAGDFL (147-183) | 1700.4973 | 8494.4345 | 40836 | 206829 | 2+2 | 43 | 0 | |
| APSTWLTAYVVKVFSLAVNLIAIDSQVLCGAVK (80-112) + NNNEKDMALTAFVLISLQEAKDICEEQVNSLPGSITK (142-178) | 1077.5705 | 7535.8996 | 45006 | 181032 | 1+2 | 60 | 0 | |
| AVNLIAIDSQVLCGAVKWLILEKQKPDGVFQEDAPVIHQEMIGGLR (96-141) + AFVLISLQEAKDICEEQVNSLPGSITKAGDFLEANYMNLQR (152-192) | 1202.1328 | 9609.035 | 43937 | 176583 | 2+2 | 23 | 0 | Oxidn. |
| APSTWLTAYVVKVFSLAVNLIAIDSQVLCGAVKWLILEK (80-118) + DICEEQVNSLPGSITK (163-178) | 999.0313 | 5988.1568 | 26246 | 96290 | 2+0 | 64 | 0 | |
| RAPSTWLTAYVVKVFSLAVNLIAIDSQVLCGAVK (79-112) + NNNEKDMALTAFVLISLQEAKDICEEQVNSLPGSITK (142-178) | 962.4995 | 7691.9935 | 20462 | 41715 | 2+2 | 47 | 0 | |

Electrospray ionization time-of-flight mass spectrometric data of chemically-linked dimeric C3d$^{17C}$ digested with trypsin for 0.5 hours at 37°C (Supplementary Figure S9e) analysed using the MassHunter Bioconfirm software. Dimeric C3d$^{17C}$ fragments corresponding to masses containing a disulphide bond between 108C and 165C of C3d are shown confirming the presence of an intact internal disulphide bond and hence indicating chemical linkage is unlikely to have occurred at this position. m/z, mass-to-charge ratio; MS/MS, tandem mass spectrometry; FDR, false discovery rate; Mod.; modification; Oxidn.; oxidation.
